# Atomically accurate de novo design of antibodies with RFdiffusion

**DOI:** 10.1101/2024.03.14.585103

**Authors:** Nathaniel R. Bennett, Joseph L. Watson, Robert J. Ragotte, Andrew J. Borst, DéJenaé L. See, Connor Weidle, Riti Biswas, Yutong Yu, Ellen L. Shrock, Russell Ault, Philip J. Y. Leung, Buwei Huang, Inna Goreshnik, John Tam, Kenneth D. Carr, Benedikt Singer, Cameron Criswell, Basile I. M. Wicky, Dionne Vafeados, Mariana Garcia Sanchez, Ho Min Kim, Susana Vázquez Torres, Sidney Chan, Shirley M. Sun, Timothy Spear, Yi Sun, Keelan O’Reilly, John M. Maris, Nikolaos G. Sgourakis, Roman A. Melnyk, Chang C. Liu, David Baker

## Abstract

Despite the central role that antibodies play in modern medicine, there is currently no method to design novel antibodies that bind a specific epitope entirely *in silico*. Instead, antibody discovery currently relies on animal immunization or random library screening approaches. Here, we demonstrate that combining computational protein design using a fine-tuned RFdiffusion network alongside yeast display screening enables the generation of antibody variable heavy chains (VHHs) and single chain variable fragments (scFvs) that bind user-specified epitopes with atomic-level precision. To verify this, we experimentally characterized VHH binders to four disease-relevant epitopes using multiple orthogonal biophysical methods, including cryo-EM, which confirmed the proper Ig fold and binding pose of designed VHHs targeting influenza hemagglutinin and *Clostridium difficile* toxin B (TcdB). For the influenza-targeting VHH, high-resolution structural data further confirmed the accuracy of CDR loop conformations. While initial computational designs exhibit modest affinity, affinity maturation using OrthoRep enables production of single-digit nanomolar binders that maintain the intended epitope selectivity. We further demonstrate the de novo design of single-chain variable fragments (scFvs), creating binders to TcdB and a Phox2b peptide-MHC complex by combining designed heavy and light chain CDRs. Cryo-EM structural data confirmed the proper Ig fold and binding pose for two distinct TcdB scFvs, with high-resolution data for one design additionally verifying the atomically accurate conformations of all six CDR loops. Our approach establishes a framework for the rational computational design, screening, isolation, and characterization of fully de novo antibodies with atomic-level precision in both structure and epitope targeting.

## Introduction

Antibodies are the dominant class of protein therapeutics, with over 160 antibody therapeutics currently licensed globally and a market value expected to reach $445 billion in the next five years^1^. Despite immense pharmaceutical interest, therapeutic antibody development still relies on animal immunization or screening of antibody libraries to identify candidate molecules that bind to a desired target. These methods are laborious, time-consuming, and can fail to produce antibodies that interact with the therapeutically relevant epitope^2^. Efforts at computational design of antibodies have grafted residues into existing antibody structures, sampled alternative native CDR loops to improve affinities^3,4^ and used Rosetta^5^ sequence design to improve the interacting regions. More recently, structure-based and sequence-based deep learning networks have been trained to design novel antibody sequences^6–8^, but de novo (no homology to an existing antibody targeting that epitope) design of structurally accurate antibodies has remained elusive. There has been recent progress in designing binding proteins (not antibodies) using RFdiffusion^9–11^ which, unlike previous methods, does not require pre-specification of the protein binder backbone, permitting the design of very diverse binders with inherent shape complementarity to the user-specified epitope^9,10^. However, as with other methods for de novo interface design^12–14^, these binders almost exclusively rely on regular secondary structure (helical or strand) based interactions with the target epitope, and the original (“vanilla”) RFdiffusion network is therefore unable to design antibodies de novo (Extended Data Fig. 1; and ref [^15^]).

An ideal method for designing de novo antibodies would enable 1) targeting of any specified epitope on any target of interest; 2) focusing of sampling on the CDR loops, keeping the framework sequence and structure close to a user-specified highly optimized therapeutic antibody framework; and 3) sampling of alternative rigid-body placements of the designed antibody with respect to the epitope. We hypothesized that it should be possible to develop a specialized RFdiffusion variant capable of designing de novo CDR-mediated interfaces, given the diversity and quality of de novo interfaces RFdiffusion can design and given that the underlying thermodynamics of interface formation are the same. RoseTTAFold2 (RF2) and RFdiffusion are trained on the entire Protein Data Bank (PDB^16^) which helps overcome the problem that the PDB contains relatively few antibody structures (∼8,100 antibody structures versus >200,000 total structures), complicating the training of large machine learning models. We set out to develop versions of RFdiffusion and RF2 specialized for antibody structure design and structure prediction by fine-tuning on native antibody structures. For simplicity, in this work, we henceforth refer to the original RFdiffusion network as “vanilla RFdiffusion”, and the antibody-specific variant we describe here simply as “RFdiffusion”.

### Fine-tuning RFdiffusion for antibody design

RFdiffusion uses the AlphaFold2^17^/RF2 frame representation of protein backbones comprising the Cɑ coordinate and N-Cɑ-C rigid orientation for each residue. During training, a noising schedule is used that, over a set number of “timesteps” (*T)*, corrupts the protein frames to distributions indistinguishable from random distributions (Cɑ coordinates are corrupted with 3D Gaussian noise, and residue orientations with Brownian motion on SO(3). During training, a PDB structure and a random timestep (*t*) are sampled, and t noising steps are applied to the structure. RFdiffusion predicts the de-noised (*pX_0_*) structure at each timestep, and a mean squared error (m.s.e.) loss is minimized between the true structure (*X_0_*) and the prediction (*pX_0_*). At inference time, starting from random residue distributions (*X_T_*), RFdiffusion iterative de-noising generates new protein structures.

To explore the design of antibodies, we fine-tuned RFdiffusion predominantly on antibody complex structures (Fig. 1; Methods). At each step of training, an antibody complex structure is sampled, and a randomly selected number (t) of noising steps are carried out to corrupt the antibody structure (but not the target structure). To permit specification of the framework structure and sequence at inference time, the framework sequence and structure are provided to RFdiffusion during training (Fig. 1B). Because it is desirable for the rigid body position (dock) between antibody and target to be designed by RFdiffusion along with the CDR loop conformations, the framework structure is provided in a global-frame-invariant manner during training (Fig. 1C). We utilize the “template track” of RF2/RFdiffusion to provide the framework structure as a 2D matrix of pairwise distances and dihedral angles between each pair of residues (a representation from which 3D structures can be accurately recapitulated)^18^, (Extended Data Fig. 1A). The framework and target templates specify the internal structure of each protein chain, but not their relative positions in 3D space (in this work we keep the sequence and structure of the framework region fixed, and focus on the design solely of the CDRs and the overall rigid body placement of the antibody against the target). In vanilla RFdiffusion, de novo binders can be targeted to specific epitopes at inference time through training with an additional one-hot encoded “hotspot” feature, which provides some fraction of the residues the designed binder should interact with. For antibody design, where we seek CDR-loop-mediated interactions, we adapt this feature to specify residues on the target protein with which CDR loops interact (Fig. 1D).

**Figure 1:**
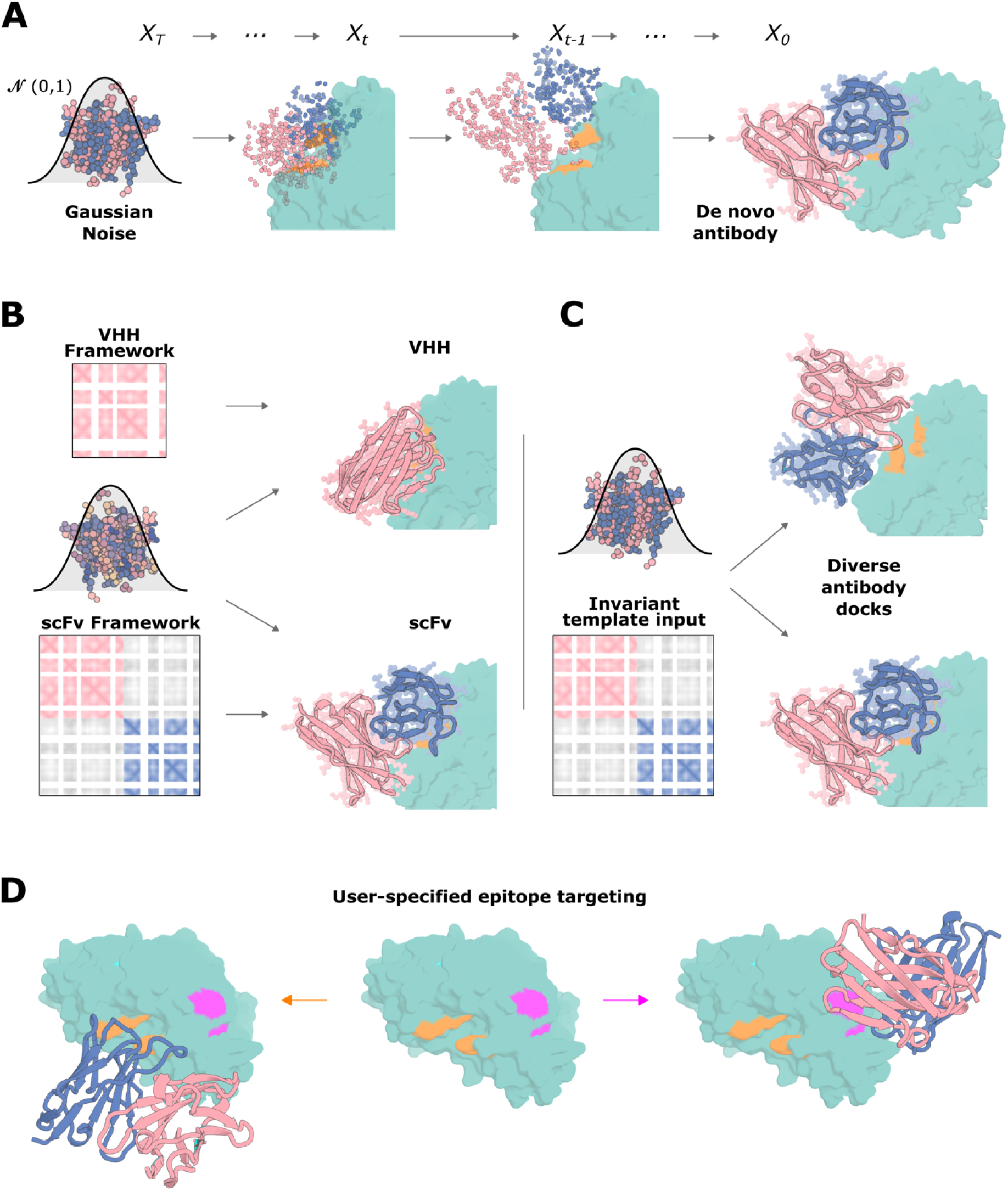
Overview of RFdiffusion for antibody design. **A**) RFdiffusion is trained such that at time *T*, a sample is drawn from the noise distribution (3D Gaussian distribution for translations, and uniform SO3 distribution for rotations), and this sampled noise is then “de-noised” between times *T* and 0, to generate an (in this case) scFv binding to the target structure through its CDR loops. **B**) Control over which framework is used is provided through input of a framework “template”, which specifies the pairwise distances and dihedral angles between residues in the framework. The sequence of the framework region is also included. For example, provision of a VHH framework generates a VHH (top row), whereas provision of an scFv framework generates a scFv (bottom row). **C**) Diversity in the antibody-target dock is achieved through the pairwise framework representation, which, because the framework structure is provided on a separate template to that of the target, does not provide information about the rigid body framework-target relationship. Hence, diverse docking modes are sampled by RFdiffusion. **D**) The epitope to which the antibody binds can be specified by provision of input “hotspot” residues, which direct the designed antibody (compare orange, left vs pink, right).

With this training regime, RFdiffusion is able to design antibody structures that closely match the structure of the input framework structure, and target the specified epitope with novel CDR loops (Extended Data Fig. 1). After the RFdiffusion step, we use ProteinMPNN to design the CDR loop sequences. The designed antibodies make diverse interactions with the target epitope and differ significantly from sequences in the training dataset (Fig. 2E).

**Figure 2:**
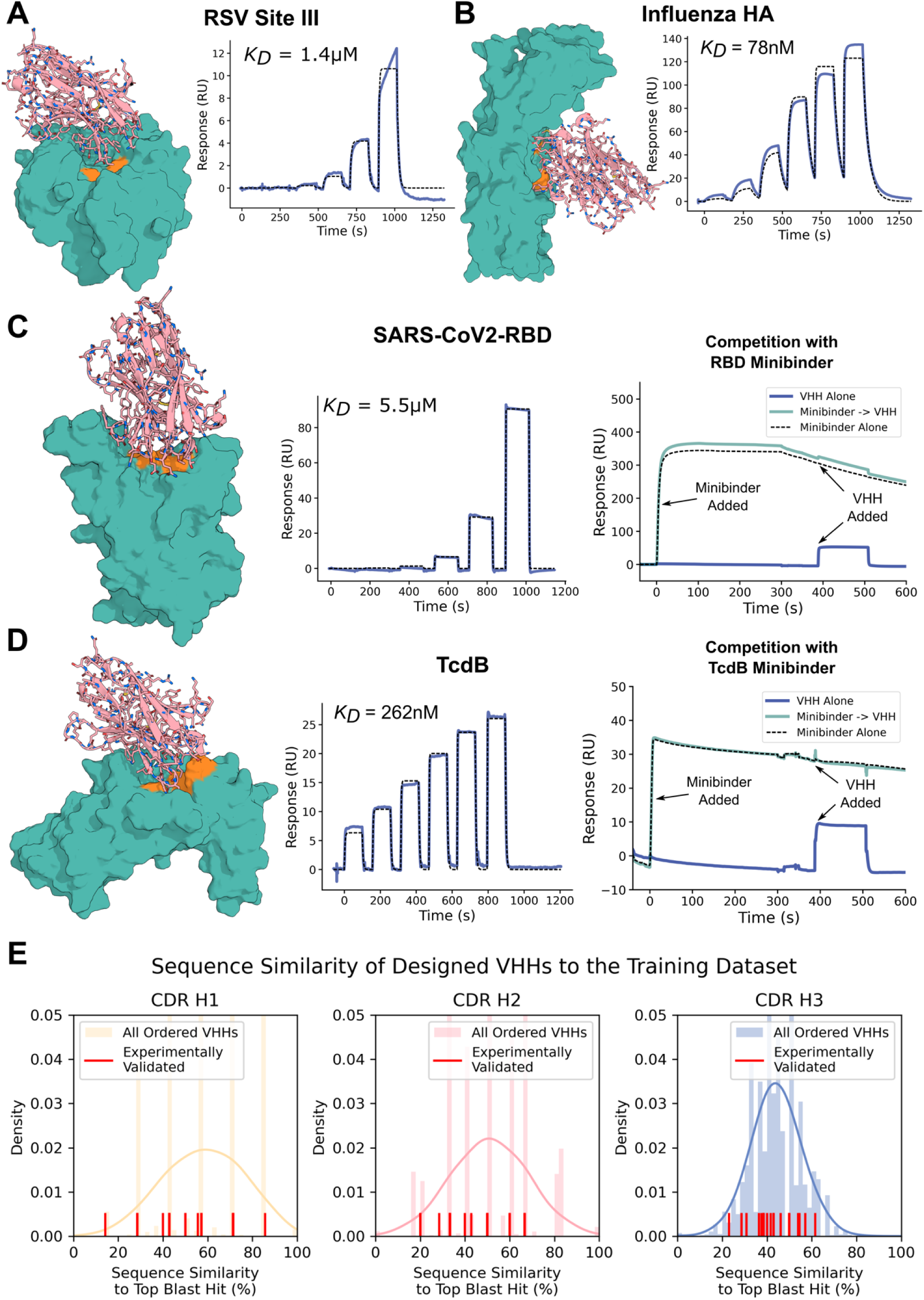
Biochemical characterization of designed VHHs. **A**-**B**) 9000 designed VHHs were screened against RSV site III (*VHH_RSV_01*) and influenza hemagglutinin (*VHH_flu_01*) with yeast surface display, before soluble expression of the top hits in *E. coli*. Surface Plasmon Resonance (SPR) demonstrated that the highest affinity VHHs to RSV site III and Influenza Hemagglutinin bound their respective targets with 1.4 μM and 78 nM respectively. **C**) 9000 VHH designs were tested against SARS-CoV-2 receptor binding domain (RBD), and after soluble expression, SPR confirmed an affinity of 5.5 μM to the target for design *VHH_RBD_D4*. Binding was to the expected epitope, confirmed by competition with a structurally confirmed de novo binder (AHB2, PDB ID: 7UHB). **D**) 95 VHH designs were tested against the *C. difficile* toxin TcdB. The highest affinity VHH, *VHH_TcdB_H2,* bound with 262 nM affinity, and also competed with a structurally confirmed de novo binder^30^ to the same epitope (right). See also Extended Data Fig. 7 for quantification of the competition shown in **C** and **D**. **E**) Designed VHHs were distinct from the training dataset. Blastp^55^ was used to find hits against the SAbDab^56^, and the similarity of the CDR loops in the top blast hit were reported for all VHHs experimentally tested in this study. Note also that the 28 VHHs confirmed to bind their targets by SPR do not show enhanced similarity to the training set (red lines).

### Fine-tuning RoseTTAFold2 for antibody design validation

Design pipelines typically produce a wide range of solutions to any given design challenge, and hence readily computable metrics for selecting which designs to experimentally characterize play an important role. An effective way to filter designed proteins and interfaces is based on the similarity of the design model structure to the AlphaFold2 predicted structure for the designed sequence (this is often referred to as "self-consistency"), which has been shown to correlate well with experimental success^19,20^. In the case of antibodies, however, AlphaFold2 fails to routinely predict antibody-antigen structures accurately^21^, preventing its use as a filter in an antibody design pipeline, and at the outset of this project, AlphaFold3^22^ was not available.

We sought to build an improved filter by fine-tuning the RoseTTAFold2 structure prediction network on antibody structures. To make the problem more tractable, we provide information during training about the structure of the target and the location of the target epitope to which the antibody binds; the fine-tuned RF2 must still correctly model the CDRs and find the correct orientation of the antibody against the targeted region. With this training regimen, RF2 is able to robustly distinguish true antibody-antigen pairs from decoy pairs and often accurately predicts antibody-antigen complex structures. Accuracy is higher when the bound (holo) conformation of the target structure is provided (Extended Data Fig. 2); this is available when evaluating design models, but not available in the general antibody-antigen structure prediction case. At monomer prediction, the fine-tuned RF2 performs better than previous models available at the time, especially at CDR H3 structure prediction (Extended Data Fig. 3).

When this fine-tuned RF2 network is used to re-predict the structure of RFdiffusion-designed VHHs, a significant fraction are confidently predicted to bind in an almost identical manner to the designed structure (Extended Data Fig. 4A). Further, in silico cross-reactivity analyses demonstrate that RFdiffusion-designed VHHs are rarely predicted to bind to unrelated proteins (Extended Data Fig. 4B). VHHs that are confidently predicted to bind their designed target are predicted to form high quality interfaces, as measured by Rosetta ddG (Extended Data Fig. 4C). The fact that many of the designed sequences generated by our RFdiffusion antibody design pipeline are predicted by RF2 to adopt the designed structures and binding modes suggested that RF2 filtering might enrich for experimentally successful binders.

### Design and biochemical characterization of designed VHHs

We initially focused on the design of single-domain antibodies (VHHs) based on the variable domain from heavy-chain antibodies produced by camelids^23^. The smaller size of VHHs makes genes encoding designs much easier to assemble and cheaper than single chain variable fragments (scFv; where linker choice can be a critical factor^24^) or fragment antigen-binding regions (Fab; where an interchain disulfide bond is required for proper folding^25^). VHHs are readily “humanized”; so far, two VHH-based therapies are approved by the FDA with many clinical trials ongoing^23^. Despite having fewer CDR loops (three) than conventional antibodies (six), the average interaction surface area of a VHH is very similar to that of an antibody^26^, suggesting a method capable of VHH design could also be suitable for antibody design. Indeed, in silico metrics for scFvs and VHHs showed similar qualities of interfaces, as assessed by Rosetta^5^ and fine-tuned RF2 (Extended Data Fig. 6).

We chose a widely-used humanized VHH framework (h-NbBcII10FGLA; [ref ^27^]) as the basis of our VHH design campaigns, and designed VHHs to a range of disease-relevant targets: *Clostridium difficile* toxin B (TcdB), influenza H1 hemagglutinin (HA), respiratory syncytial virus (RSV) sites I and III, SARS-CoV-2 receptor binding domain (RBD) and IL-7Rɑ. ProteinMPNN^28^ was used to design the sequences of the CDR loops (but not the framework) in the context of the target. We then filtered designs with the fine-tuned RoseTTAFold2 network described above. Designs were screened either at high-throughput by yeast surface display (9000 designs per target; RSV sites I and III, RBD, Influenza HA) or at lower-throughput with *E. coli* expression and single-concentration surface plasmon resonance (95 designs per target; TcdB, IL-7Rɑ and influenza HA–the latter was screened using both methods).

In the case of influenza HA, glycan N296, located along the HA-stem epitope, exhibited varying degrees of overlap with the approach angle of several of our designed VHHs. To best align the experimental design conditions with the computational parameters employed during design (i.e., excluding consideration of the glycan shield), affinity measurements were conducted using a commercially produced monomeric HA product expressed in insect cells (Extended Data Fig. 11). Insect cells express a truncated paucimannose glycan shield, which (relative to a natively expressed HA trimer) more closely resembles the fully deglycosylated HA monomeric PDB model used for VHH design. Of the HA binders tested against the insect-cell produced HA monomer, the highest affinity binder had a K_D_ of 78 nM, (Fig. 2B), with other binders having affinities of 546 nM, 698 nM, and 790 nM.

The highest affinity binders to RSV site III, Influenza HA, RBD, and TcdB are shown in Fig. 2A,B,C,E respectively (see also Extended Data Fig. 8 for all the SPR traces of confirmed VHH binders identified in this study). The CDR loops are distinct from VHHs observed in nature, indicating significant generalization beyond the training dataset (Fig. 2E, Extended Data Fig. 5). For TcdB, the target epitope was the Frizzled-7 interface for which there are no antibodies or VHHs targeting this site in the PDB. For the best designed VHH from both RBD (K_D_ = 5.5 μM; Fig. 2C) and TcdB (K_D_ = 260 nM; Fig. 2D) binding was confirmed to be to the desired epitope: binding was completely abolished upon addition of a previously designed, structurally characterized de novo binder to that epitope (AHB2, PDB ID: 7UHB^29^ for RBD and Fzd48^30^ for TcdB) (Fig. 2C,D; Extended Data Fig. 7). This TcdB VHH also neutralized TcdB toxicity in CSPG4 knockout cells (an alternative TcdB receptor) with an EC50 of 460 nM (Extended Data Fig. 9). For TcdB, the interactions were specific, with no binding observed to the highly related (70% sequence homology) *Clostridium sordellii* lethal toxin L (TcsL) (Extended Data Fig. 7B). These data demonstrate the ability of RFdiffusion to design VHHs that make specific interactions with the target epitope.

### Cryo-EM reveals atomically accurate VHH design against a natively glycosylated influenza viral glycoprotein

Given the success of RFdiffusion at generating moderate affinity VHHs against diverse epitopes, we sought to evaluate design accuracy by cryo-EM structure determination of the designed anti-HA VHHs in complex with natively glycosylated, trimeric influenza HA glycoprotein (strain A/USA:Iowa/1943 H1N1), which retains the conserved stem epitope used during computational VHH design and upstream biochemical screening. The VHHs were combined with Iowa43 HA at a 3:1 molar excess ratio (VHH:HA monomer) at a concentration of 15 μM and promptly prepared for cryo-EM grid freezing. Cryo-EM data processing revealed one VHH design effectively bound to the fully glycosylated HA trimer (out of the four tested), denoted hereafter as *VHH_flu_01* (Fig. 3). 2D classification of all particles in the dataset (Fig. 3A) and the determined 3.0 Å structure of the complex (Fig. 3B, Supplementary Methods Table 9) identified approximately 66% of HA particles bound to a maximum of two VHHs per trimer (Fig. 3A-H). This partial occupancy is likely attributable to the N296 glycan, which in unbound subunits partially occludes the target epitope but reorients when bound to *VHH_flu_01* (see Fig. 3H).

**Figure 3:**
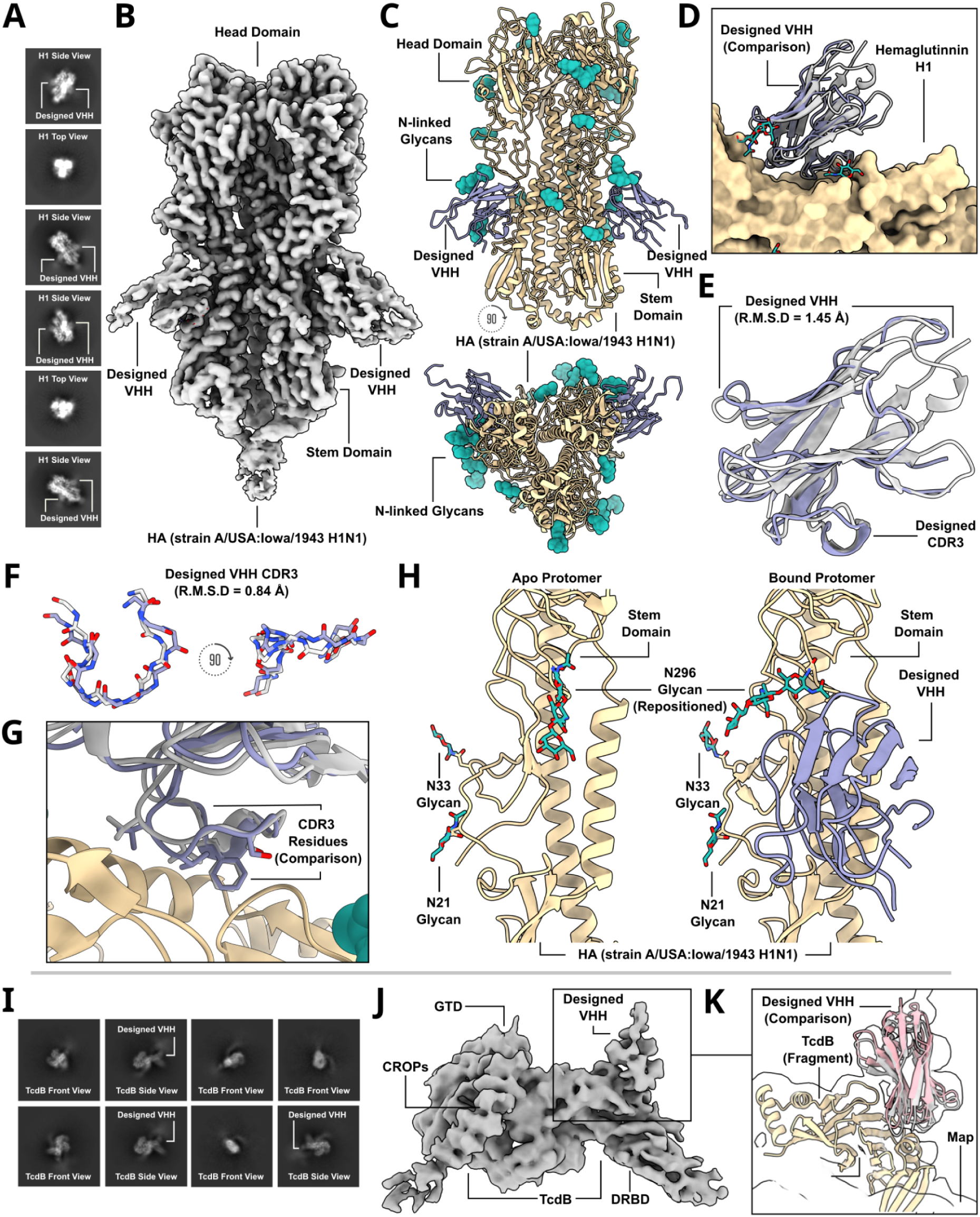
Cryo-EM structural characterization of two de novo designed VHH binding to influenza hemagglutinin and TcdB. **A**) Labeled cryo-EM 2D class averages of a designed VHH, *VHH_flu_01*, bound to influenza HA, strain A/USA:Iowa/1943 H1N1. **B**) A 3.0 Å cryo-EM 3D reconstruction of the complex shows *VHH_flu_01* bound to H1 along the stem in two of the three protomers. **C**) Cryo-EM structure of *VHH_flu_01* bound to influenza HA. **D**) The cryo-EM structure of *VHH_flu_01* in complex with HA closes matches the design model. **E**) cryo-EM reveals the accurate design of *VHH_flu_01* using RFdiffusion (RMSD to the RFdiffusion design of the VHH is 1.45 Å). **F**) Superposition of the designed VHH CDR3 predicted structure as compared to the built cryo-EM structure (RMSD = 0.84 Å). **G**) Comparison of predicted CDR3 rotamers compared to the built 3.0 Å cryo-EM structure. **H**) Examination of apo HA protomers juxtaposed with those bound to the designed VHH unveils a notable repositioning and accommodation of glycan N296 to allow for binding of the designed VHH to the HA stem. In each structural depiction panel, the designed VHH predicted structure is showcased in gray, while the cryo-EM solved structure of the designed VHH is depicted in purple. Additionally, the HA glycoprotein is represented in tan, and the HA glycan shield is illustrated in green. **I**) Labeled cryo-EM 2D class averages of the designed and affinity-matured VHH, *VHH*_*TcdB_H2_ortho*, bound to full-length TcdB. **J**) A 5.7 Å cryo-EM 3D reconstruction of the complex shows *VHH*_*TcdB_H2_ortho* bound to the target epitope as predicted. **K**) Due to the modest resolution, a fragment of TcdB was first docked into the cryo-EM density map, and the full design model—including both the TcdB fragment and the designed VHH—was then aligned to the pre-fitted TcdB fragment. This approach demonstrates that the predicted design closely matches the experimentally determined complex in structure, epitope targeting, and overall conformation. (Yellow = HA; purple = VHH (cryo-EM); gray = VHH (computational design prediction)

The structure of influenza HA bound to two copies of *VHH_flu_01* (Figure 3B,C, Extended Data Fig. 12) reveals a VHH approach angle which closely matches the predicted model (Fig. 3D), and a VHH backbone which is very close to the RFdiffusion design, with a calculated RMSD of 1.45 Å (Fig. 3E). The CDR3 structure is also very similar between the cryo-EM structure and the computational model (RMSD = 0.8 Å) (Fig. 3F), with residues V100, V101, S103, and F108 in the de novo designed CDR3 loop interacting with the influenza HA stem epitope in the cryo-EM structure, as designed by RFdiffusion and re-predicted with RF2 (Fig. 3G). Notably, the design is highly dissimilar from the closest antibody/VHH binding to this epitope in the PDB (Extended Data Fig. 5G,H). Taken together, these results demonstrate VHH design with atomic level precision.

### Cryo-EM characterization of affinity-matured VHHs targeting *C. difficile* TcdB and SARS-CoV-2 RBD

To improve the binding affinity of *de novo* designed VHHs, we utilized the OrthoRep system for continuous hypermutation of target genes *in vivo*.^31,32^ OrthoRep has been shown to drive the rapid affinity maturation of yeast surface-displayed antibodies. We used this capability to affinity mature VHHs targeting *Clostridioides difficile* toxin B (TcdB), Influenza H1 Hemagglutinin, and the SARS-CoV-2 receptor binding domain (RBD). In all cases, affinity-matured VHHs acquired several mutations relative to the parent designs and improved binding affinities by approximately two orders of magnitude (Extended Data. Fig. 10), making them suitable candidates for downstream cryo-EM structural characterization. Structural predictions of the affinity-matured complexes suggested that the mutations retained the original target site specificity. Notably, for the SARS-CoV-2 RBD-targeting VHH, the affinity-enhancing mutations were not localized to the predicted binding interface (Extended Data Fig. 10D).

For *Clostridioides difficile* toxin B (TcdB), our design campaign targeted the Frizzled-7 epitope located on the receptor binding domain that lacks known natural antibody or VHH binding partners in the Protein Data Bank (PDB). TcdB consists of four functional domains: an N-terminal glucosyltransferase domain (GTD), a cysteine protease domain (CPD), a C-terminal domain of combined repetitive oligopeptides (CROPs), and a central delivery and receptor-binding domain (DRBD). Cryo-EM characterization of the original parent design, *VHH_TcdB_H2*, confirmed that the VHH engages the target Frizzled-7 DRBD epitope. Analysis via 2D and 3D classification revealed a mix of bound and unbound TcdB particles (Extended Data Figs. 14), a result observed only after adding glycine to prevent nonspecific aggregation of the VHH^33–35^ (Extended Data Fig. 13). In contrast, subsequent cryo-EM studies of the affinity-matured VHH, *VHH_TcdB_H2_ortho*—conducted without glycine—showed a high proportion of TcdB particles bound by the VHH, consistent with its enhanced affinity (Figure 3I,J,K, Extended Data Fig. 10B). To address TcdB flexibility and improve map quality at the VHH-TcdB interface, we performed extensive 3D classifications and local refinements. This approach identified multiple structural states of TcdB within the dataset: a compressed apo state, a compressed bound state, an extended apo state, and an extended bound state (Extended Data Figs. 15-16). By isolating the bound conformations, we were able to resolve the VHH-TcdB complex to a modest 5.7 Å resolution, enabling us to confidently dock the designed VHH into the cryo-EM density, confirm that the VHH folded into the expected architecture and bound to the target epitope as designed (Fig. 3J-K, Extended Data Fig. 16). These results underscore the capability of RFdiffusion to design accurate *de novo* VHHs capable of targeting previously unexplored and structurally flexible epitopes.

We next used cryo-EM to characterize an affinity-matured VHH (*VHH_RBD_D4_ortho19)*) targeting the SARS-CoV-2 spike RBD, where competition experiments indicated that the parental VHH bound the intended epitope (Fig. 2C, Extended Data Fig. 7C, Extended Data Fig. 10B, Extended Data Fig. 17). The RBD transitions between "up" and "down" conformations, with the "up" state enabling receptor binding and viral entry^36^. Cryo-EM 2D class averages and 3D classification reconstructions of the VHH-bound complex revealed a mixture of RBD conformations (1–2 "up"), with VHH density observed exclusively in the "up" state. This is consistent with its design, as the target epitope is occluded in the "down" conformation (Extended Data Fig. 17A–B). A global refinement with an average estimated resolution 3.9 Å provided well-defined density for the lower portion of the spike protein (local resolution = ∼2.5 Å). However, the relative flexibility of the RBD resulted in substantial signal averaging, causing density loss at higher contour levels which precluded assessment of VHH design accuracy (Extended Data Fig. 17C–E). Symmetry expansion and local refinements helped improve the resolution of the RBD–VHH interface, confirming the intended VHH fold and accurate epitope targeting following rigid-body docking of the design model into the density map (Extended Data Fig. 17F–G), a result in agreement with our biochemical competition data (Fig. 2C). However, while the VHH bound the correct RBD epitope, its binding mode surprisingly deviated significantly from the design model, instead adopting a predominantly framework-mediated interaction that more closely matched retrospective AlphaFold3 predictions (Extended Data Fig. 17G–H). Due to the significant deviation between the designed docks and the experimentally-determined dock, we classified this as a design *failure*.

### Design of scFvs with six designed CDRs

Given the success of RFdiffusion at designing VHHs with three de novo CDRs, we next tested its ability to design both heavy and light chains in single-chain Fv (scFv) format. RFdiffusion was used to generate scFvs targeting specific epitope sites, following a strategy similar to the VHH design approach. However, unlike VHHs, where only three CDRs were built de novo, scFv design involved constructing all six CDRs on both the heavy and light chains in addition to the docking mode.

The gene synthesis problem is more formidable for scFvs than for VHHs as they are too long to be simply assembled from pairs of conventional oligos synthesized on oligonucleotide arrays, and are challenging to uniquely pair due to high sequence homology between scFvs. We developed stepwise assembly protocols that enable the construction of libraries with heavy and light chains either specifically paired as in the design models (Extended Data Fig. 19 and 20) or combinatorially mixed within subsets of designs specifically with similar target binding modes (Extended Data Fig. 21). The latter approach helps overcome the greater challenge of accurate design of six CDRs de novo, which increases the possibilities for error compared to the VHH problem as only one suboptimal CDR can compromise binding. We found that given sets of nearly superimposable designs targeting the same site with the same binding mode, new scFvs generated by combining pairs of heavy and light chains from different designs were confidently predicted to bind to the target site in the designed binding mode at similar frequencies as the original designs (Extended Data Fig. 22A). This is not surprising given the extended nature of the scFv interface: within the sets of designs with the same binding orientation, the heavy chain and light chain CDRs make interactions with different regions and hence can be combined without loss of structural precision. In contrast, random, structure-agnostic pairing rarely led to predicted binders (Extended Data Fig. 22A). Hence by mixing CDRs from different designs that bind in the same orientation we can effectively overcome failures due to single imperfectly designed CDRs–a combinatorial solution to a combinatorially more complex problem (two-chain ScFv design vs one-chain VHH design). This strategy highlights a key advantage of structure-based design: “intelligent” pairing of heavy and light chains is possible with a structural model of every antibody, and allows de novo designed antibody libraries to reach scales attainable by traditional library assembly methods, even despite current limitations in gene synthesis.

We succeeded in identifying epitope specific scFvs from the heavy-light combinatorial libraries but not the fixed pairing libraries (Extended Data Fig. 23). Following expression and purification, SPR analysis of 6 distinct scFvs originating from two unique docks targeting the Frizzled epitope of TcdB revealed a range of affinities; the highest affinity binder, *scFv6*, had a K_D_ of 72 nM (Fig. 4C). There are no antibodies binding to this epitope in the PDB, hence this success cannot be attributed to memorization. Subsequent prediction of the structure of the scFv with AlphaFold3 showed a binding mode identical to that of the two nearly superimposable parent designs that contributed the light and heavy chains (Extended Data Fig. 29C-D). Competition with a native binder, Frizzled-7 to this epitope confirmed that the binding was on target (Fig. 4D). In contrast, no competition was seen in the presence of CSPG4, an alternative receptor that interacts with an epitope at the toxin core. Thus, scFvs targeting user-specified epitopes can be identified from structure-aware designed combinatorial libraries.

**Figure 4:**
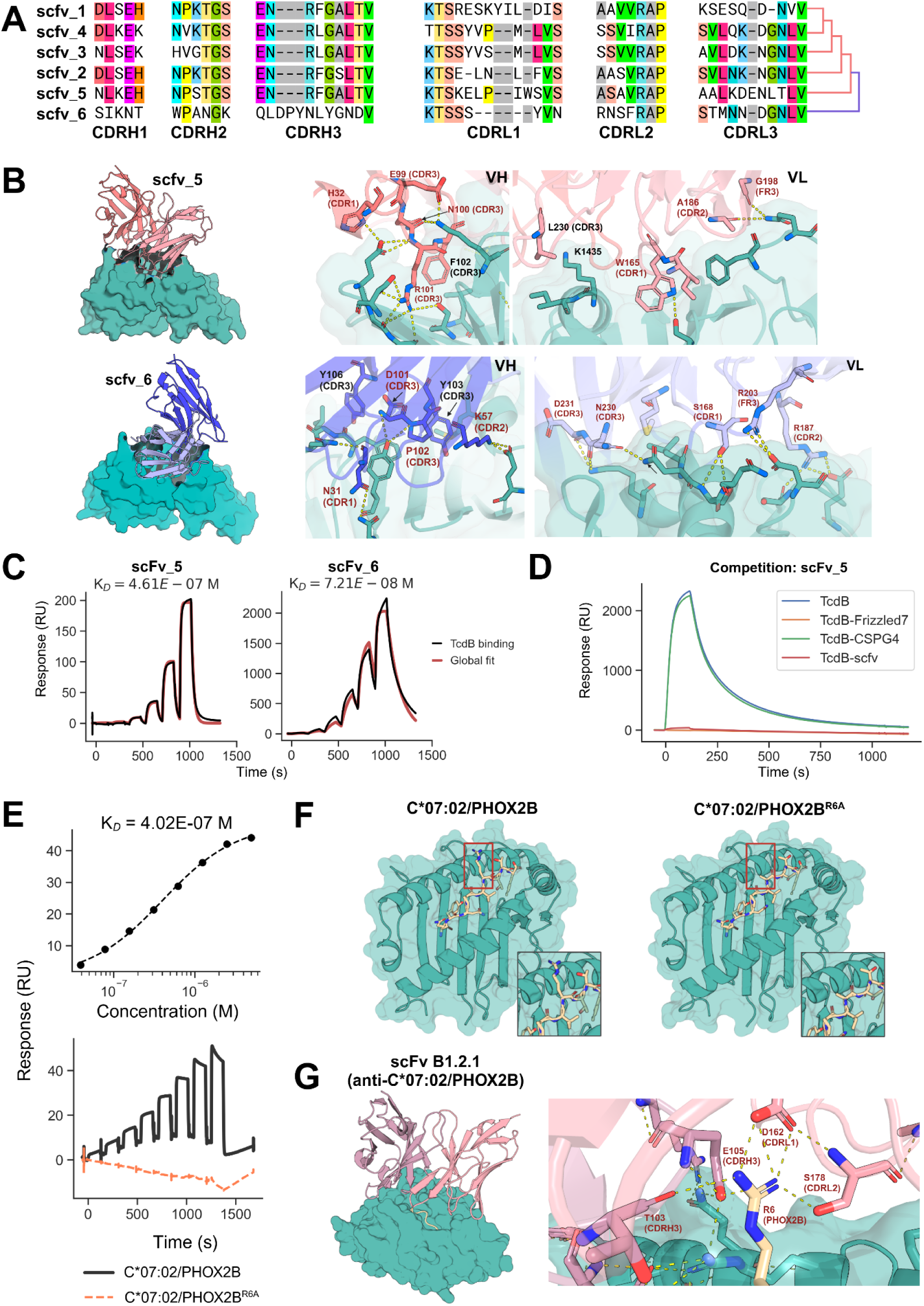
Biochemical characterization of combinatorially-assembled scFvs with six designed CDRs. **A**) Multiple sequence alignment of 6 scFvs that bind TcdB. The first five scFvs originate from the same structural cluster, which *scFv6* originates from a distinct cluster. **B**) AlphaFold3 predictions of *scFv5* and *scFv6* in complex with TcdB receptor binding domain. Both *scFv5* and *scFv6* are predicted to bind a similar but not identical epitope. The predicted orientation of *scFv6* relative to TcdB is rotated in comparison to *scFv5* (left). CDRH3 of both scFvs are predicted to make several polar contacts with the target (right). VL of *scFv6* is also predicted to make several polar contacts via all 3 CDR loops. VL of *scFv5* exhibits more hydrophobic packing to the target. **C**) SPR analysis of scFv binding to the receptor binding domain of TcdB. Each scFv was conjugated to a CM5 chip and TcdB RBD was titrated across a 6-step 4-fold dilution curve with an upper concentration of 1 μM **D**) scFv was conjugated to a CM5 chip and then TcdB RBD was flowed over at 50 nM either alone or mixed with 1 μM of Frizzled-7, CSPG4 or the same scFv as was conjugated to the chip. **E**) SPR comparative analysis of B1.2.1 binding C*07:02/PHOX2B versus C*07:02/PHOX2B^R6A^. scFv was immobilized to a CM5 chip and then on- and off-target binding was measured across an 8-step 2-fold titration with an upper concentration of 5 μM. Steady state kinetic analysis (top) and raw SPR trace of on- and off-target binding (lower) indicate specific binding to the intended target. **F**) AlphaFold3 predictions of HLA-C*07:02 with peptide PHOX2B (left) and PHOX2B^R6A^ (right). R6 of PHOX2B is predicted to be solvent-exposed. **G**) AlphaFold3 prediction of scFv B1.2.1 in complex with C*07:02/PHOX2B (left). Predicted polar contacts with R6 of the PHOX2B peptide (right), mediated by CDRH3, CDRL1, and CDRL2. Polar contacts were visualized in PyMOL.

We next targeted a clinically-relevant epitope – the QYNPIRTTF peptide derived from the *PHOX2B* neuroblastoma-dependency gene and master transcriptional regulator in complex with the major histocompatibility complex (MHC) allotype HLA-C*07:02 (we refer to this peptide below simply as Phox2b). The Phox2b peptide was originally discovered by immunopeptidomics of neuroblastoma patient-derived samples, and has been targeted with peptide-centric chimeric antigen receptors (PC-CARs) for treating high-risk neuroblastoma^37^. However, the PC-CARs identified previously are restricted to recognizing Phox2b presented on HLAs of the A9 serological group, excluding the common allotype HLA-C*07:02^38^. Targeting the Phox2b:HLA-C*07:02 complex could significantly increase the addressable patient population for these immunotherapies, and has been the focus of ongoing therapeutics development.

Recently, computationally designed binders for Phox2b/HLA-C*07:02 have been developed, using the TRACeR-I system^39^, while high-affinity TCRs have been identified for targeting peptides on the common HLA-C*08:02/HLA-C*05:01 allotypes^40^. An scFv (C3) targeting an HIV epitopic peptide presented on HLA-C*07:02 with picomolar-range binding affinity can kill infected cells i*n vitro*, highlighting the feasibility of using antibodies to target peptide antigens on HLA-C*07:02^41^. A benefit of structure-based design is the ability to target specific peptide residues to achieve binding specificity (rather than binding only to the MHC), and we therefore used RFdiffusion to target the R6 residue, known to be important for binding in the PC-CAR^38^. Given the low stability of the HLA-C*07:02/Phox2b complex (T_m_ of 44.2°C, ref [^42^]), we leveraged our disulfide-stabilized approach to prepare a stabilized form of the pHLA target^43^. Using the combinatorial assembly approach described above, we identified modest affinity (400 nM as measured by SPR and 1 μM as measured by ITC; Fig. 4E, Extended Data Figure 25D) scFv binders to Phox2b:HLA-C*07:02. Binding was specific to the peptide, with no detectable binding to the R6A point mutant Phox2b peptide (Phox2b^R6A^:HLA-C*07:02) (Fig. 4E). Attempts were made to incorporate scFv binders into a 4-1BB-CAR, but T cell cytotoxicity assays demonstrated no detectable killing of a range of neuroblastoma cell lines (Extended Data Fig. 26), likely because of the modest binding affinity or low levels of antigen density expressed on the tumor cells. Although there is still considerable room for improvement in affinity, this demonstrates the ability of structure-based antibody design, paired with appropriate library assembly methods, to design specific binders to challenging and clinically important target epitopes.

### Cryo-EM reveals atomically accurate scFv design against *C. difficile* Toxin B

To evaluate the accuracy of de novo scFv design, we determined the cryo-EM structures of two combinatorially-assembled scFvs, *scFv5* and *scFv6*, both targeting the Frizzled-7 epitope of TcdB. Cryo-EM analysis confirmed that both scFvs bound the Frizzled-7 epitope as designed (Fig. 5). High-resolution 2D class averages of *scFv6* revealed clear density for both TcdB and the bound scFv, further supported by a 3.6 Å 3D reconstruction (Fig. 5A-B). The resolved structure showed that *scFv6* engaged the Frizzled-7 epitope along its DRBD domain with the predicted binding orientation (Fig. 5C, Supplementary Methods Table 9). Superposition of the cryo-EM structure with the design model demonstrated remarkable agreement, with both heavy and light chains interacting with the epitope as intended (Fig. 5D, Extended Data Fig. 29A-B). The overall fold closely matched the computational model (RMSD = 0.9 Å), and each of the six complementarity-determining regions (CDR) exhibited near-atomic precision (backbone RMSDs: CDRH1 = 0.4 Å; CDRH2 = 0.3 Å; CDRH3 = 0.7 Å; CDRL1 = 0.2 Å; CDRL2 = 1.1 Å; CDRL3 = 0.2 Å) (Fig. 5E-F). This agreement extended to the rotameric conformations of CDR side chains and their interactions with the Frizzled-7 epitope, underscoring the accuracy of RFdiffusion in designing de novo scFv-target interactions (Fig. 5G).

**Figure 5:**
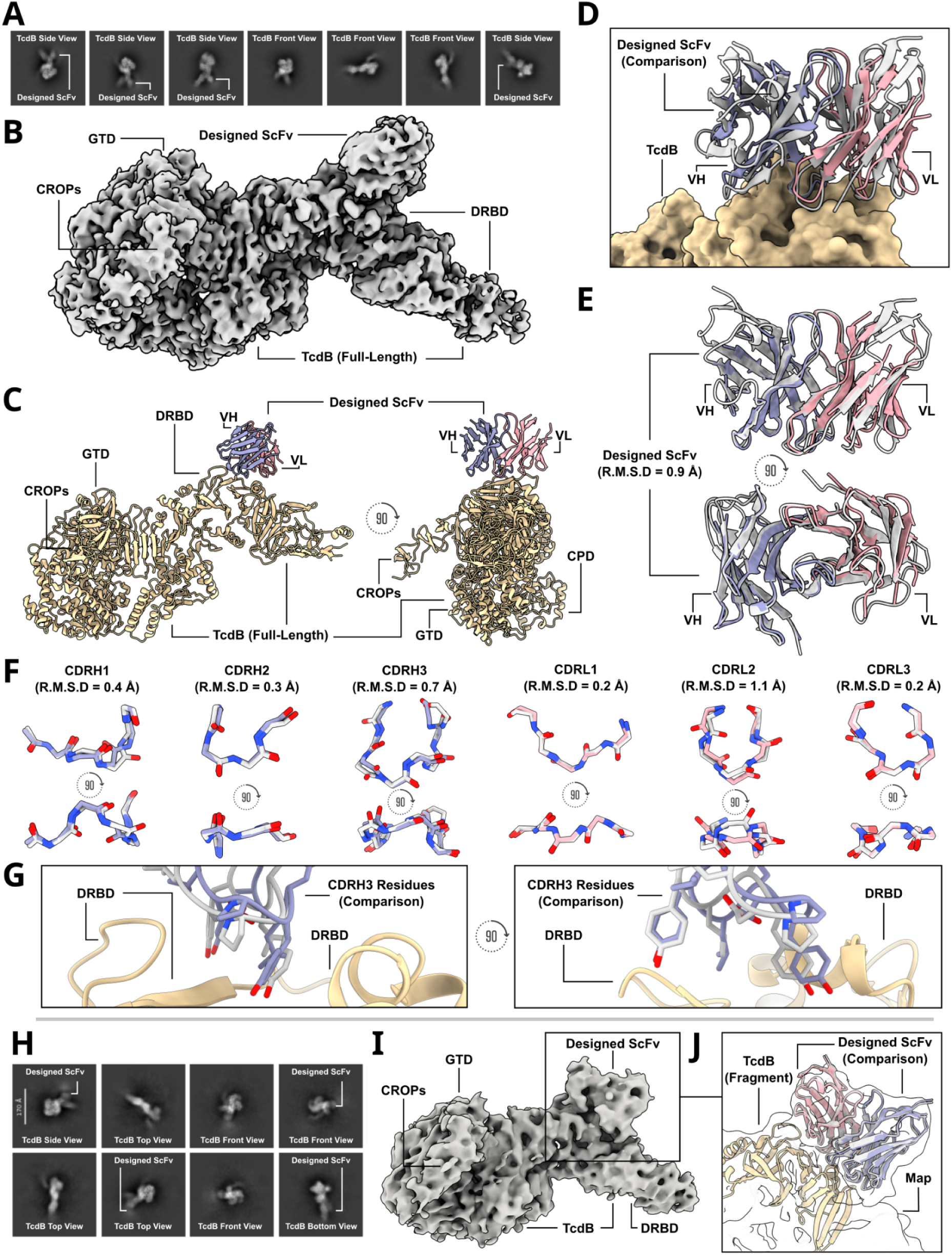
Cryo-EM structural characterization of two TcdB-binding scFvs. **A**) Labeled cryo-EM 2D class averages of a designed scFv, *scFv6*, bound to TcdB. **B**) A 3.6 Å cryo-EM 3D reconstruction of the complex shows *scFv6* bound to TcdB along the Frizzled-7 epitope. **C**) Cryo-EM structure of *scFv6* bound to TcdB. **D**) The cryo-EM structure of *scFv6* in complex with TcdB closes matches the design model. **E**) Cryo-EM reveals the accurate design of *scFv6* using RFdiffusion (RMSD to the RFdiffusion design of the scFV is 0.9 Å). **F**) Superposition of each of the six designed *scFv6* CDR loop predicted structures as compared to the built cryo-EM structure (RMSD values: CDRH1 = 0.4 Å; CDRH2 = 0.3 Å; CDRH3 = 0.7 Å; CDRL1 = 0.2 Å; CDRL2 = 1.1 Å; CDRL3 = 0.2 Å). **G**) Comparison of predicted CDRH3 rotamers compared to the built 3.6 Å cryo-EM structure. **H**) Labeled cryo-EM 2D class averages of the designed scFv, *scFv5*, bound to full-length TcdB. **I**) A 6.1 Å cryo-EM 3D reconstruction of the complex shows the *scFv5* bound to the target epitope as predicted. **J**) Due to the modest resolution, a fragment of TcdB was first docked into the cryo-EM density map, and the full design model—including both the TcdB fragment and the designed scFv—was then aligned to the pre-fitted TcdB fragment. This approach demonstrates that the predicted design closely matches the experimentally determined complex in structure, epitope targeting, and overall conformation. (Yellow = TcdB; purple = variable heavy chain fragment (cryo-EM); pink = variable light chain fragment (cryo-EM); gray = computational design prediction)

*scFv5* was designed to bind the same epitope but with an orthogonal approach angle relative to *scFv6* (Fig. 4B). A 6.1 Å cryo-EM reconstruction confirmed *scFv5* binding to the TcdB Frizzled-7 epitope, with 2D class averages showing clear density for the complex (Fig. 5H). Rigid-body docking of the designed model into the cryo-EM density revealed close agreement between the predicted and experimentally determined binding modes (Fig. 5I-J).

### Improved structure prediction oracles will increase success rate

While our results demonstrate that the de novo design of antibodies is possible, the quite low experimental success rates currently necessitate the relatively high-throughput screening methods used in this study. A key contributor to previous successes in de novo binder design was improved filters (primarily AlphaFold2^44^), which enriched for experimental success in the subset of designs that are tested experimentally^9,20^. At the outset of this study, we sought to build such a filter by fine-tuning RoseTTAFold2 (Extended Data Figs. 2-4), but the filtering power of this model is limited (at least with the settings used; providing 100% of interface “hotspots”). This likely accounts for the low experimental success rates and the inaccurate SARS-CoV-2 design, where the overall fold and epitope targeting were correct, but the binding orientation was not.

Subsequent to the design work in this study, AlphaFold3^22^ was released that has improved antibody structure prediction accuracy^22,45^. Retrospectively, we can assess how filtering with AlphaFold3 would have improved experimental success rates.

First, AlphaFold3 accurately predicts the experimentally validated structure of the inaccurately designed SARS-CoV-2 VHH (Extended Data Fig. 17). Had AlphaFold3 been used as an initial filter, this design would have been rejected due to the discrepancy between the predicted and intended structures, thereby preventing its experimental testing.

Secondly, we predicted the structures of the SARS-CoV-2, influenza HA, TcdB and IL-7R designs using AF3 with a multiple sequence alignment (MSA) and template for the target and only a template for the VHH (since CDRs are de novo, we reasoned the MSA would be of limited utility). We analyzed the predictions for libraries with at least one structurally validated VHH (TcdB, influenza HA and SARS-CoV-2). These results are dominated by anti-HA VHHs as the majority of successful binders came from this library. We find that the AF3 iPTM score is predictive of binding success (AUC = 0.86, Extended Data Fig. 18A-B). Overall, only 9% of our ordered VHH designs have an iPTM score > 0.6 suggesting that success rate will be improved by incorporation of an iPTM filter. We ran a similar analysis for the combinatorially-assembled scFv libraries; we predicted the structures of the parental scFv designs (before combinatorial assembly) and the experimentally-confirmed scFv designs (combinatorially-assembled) using AF3 with an MSA for the target sequence and a template for the target as well as the heavy and light chains, taking the max iPTM score over 10 seeds. We find that successful designs cluster to higher AF3 iPTM scores than the parental designs (Extended Data Fig. 18C). Only 4% of the initial design library has iPTM > 0.85 while 5 out of the 6 experimentally-confirmed designs pass this threshold, again suggesting that filtering by AF3 iPTM should increase success rates (Extended Data Fig. 18D).

## Discussion

Our results demonstrate that de novo design of antibody domains targeting specific epitopes on a target is possible. The cryo-EM structural data for the designed VHHs to influenza HA and TcdB reveals very close agreement to the computational design models, showing that our approach can design VHH complexes with atomic accuracy – including the highly variable H3 loop and the overall binding orientation – that are highly dissimilar from any known structures in the PDB. Moreover, cryo-EM structural data of designed scFvs bound to TcdB demonstrate the ability of RFdiffusion to design two-chain scFvs accurately. To our knowledge, these are the first experimentally validated cases of de novo-designed antibodies with structurally accurate binding. Furthermore, the success of this approach cannot be attributed to memorization, as no antibodies targeting the TcdB epitope exist in the PDB, reinforcing the model’s capability to design truly de novo binders.

Our computational method synergizes with experimental screening approaches developed for retrieving antibodies from large random libraries in several ways. First, yeast display selection methods widely used for antibody library screening enable the retrieval of the highest affinity binders amongst large sets of designs, which is currently necessary due to the quite low design success rate. Second, screening combinatorial libraries that mix heavy and light chains from designs with similar binding modes allows for the identification of scFvs targeting specific epitopes, as demonstrated here for TcdB and Phox2b-peptide MHC. In our case, this approach is particularly advantageous because a large fraction of recombinants are likely functional due to their structural compatibility. Third, affinity maturation using OrthoRep^46^ improves the measured affinity of initial VHH designs down to the single digit nanomolar or subnanomolar range, while preserving the original designed binding mode. From a practical standpoint, the key advance of this work is not merely the ability to generate nanobodies and scFvs against a target—something often achievable through purely experimental methods—but rather the ability to accurately target specific binding epitopes. Indeed, cryo-EM studies unambiguously confirmed that our computationally designed antibodies engage their intended epitopes with remarkable precision. This high degree of design accuracy is critical for therapeutic applications such as antagonists that block receptor-ligand interactions, antibodies that avoid competing with endogenous molecules, modulators that induce conformational changes to trigger signaling, or antibodies targeting conserved or evolutionarily restricted viral epitopes.

Although our results demonstrate successful de novo design of antibodies, there is considerable room for improvement. For the backbone design step, incorporating recent architectural improvements^47^ and new advances in generative modeling^48–50^ may yield design models with higher designability and diversity. RoseTTAFold2 and RFdiffusion have also recently been extended to model all biomolecules (rather than just proteins)^51^, and incorporating this capability into the antibody design RFdiffusion should permit the accurate design of antibodies to epitopes containing non-protein atoms, such as glycans. Indeed, the sub-stoichiometric binding observed for *VHH_flu_01* could be explained by the presence of nearby glycan N296, which was not considered during the initial design of this VHH. ProteinMPNN was not modified in this current work, but designing sequences that more closely match human CDR sequences would be expected to reduce the potential immunogenicity and developability of designed antibodies^52,53^ Finally, further improvements in antibody prediction methods should allow better in silico benchmarking of upstream design methods, and improve experimental success rates. Converting scFvs to full antibodies should be straightforward, as the variable CDR sequences remain identical. Alternatively, the methods used here can be directly applied to Fab design by simply omitting the linker between the heavy and light chains. With these advancements, computational antibody design can be expected to become routine.

Ultimately, computational de novo design of antibodies using our RFdiffusion and related approaches could revolutionize antibody discovery and development. Our RFdiffusion approach allows for the precise targeting of specific epitopes on a given antigen. As the method improves and success rates increase, it has the potential to be significantly faster and more cost-effective than immunizing animals or screening random libraries, which require extensive follow-up for epitope mapping and structural characterization. A structure-based approach to antibody design enables the optimization of key pharmaceutical properties, such as aggregation, solubility, and expression levels, in a structurally informed manner. This allows for targeted modifications while avoiding mutations that could disrupt the antibody-target interface or compromise antibody stability. Furthermore, the ability to explore the full space of CDR loop sequences and structures from the start, particularly for CDR1 and CDR2 which are natively limited to the space of sequences encoded by germline V genes prior to somatic hypermutation, should simplify both the optimisation of the developability features and the targeting of non-immunodominant epitopes^54^. Altogether, we expect that rational design of antibodies will dramatically increase the number of tractable clinical targets and conditions accessible to antibody therapeutics.

## Acknowledgements

We thank Phil Bradley for use of the TCR distillation dataset. We thank Minkyung Baek and Frank DiMaio for training RoseTTAFold2. We thank Brian Coventry for the use of the de novo miniprotein binder dataset. We also thank Jacob Gershon for early contributions to this project, and Justas Dauparas and Hetu Kamisetty for helpful discussions. We thank Jean-Philippe Julien and Iga Kucharska (The Hospital for Sick Children) for providing recombinant Frizzled-7 and CSPG4. We thank Yoann Aldon and Rogier Sanders (Amsterdam University Medical Center) for providing recombinant HIV env protein. We thank Nicole Roullier for help with sample preparation for next-generation sequencing. We also thank Twist Biosciences for access to their 400 bp oligo synthesis, which was invaluable for the high-throughput VHH experiments. We thank Anne Dosey for providing target protein, and Abishai Ebenezer, Amir Motmaen and Bingxu Lu for helpful discussions. We thank Ian Haydon for help with graphics. Finally, we thank Lance Stewart, Lynda Stuart, Kandise VanWormer and Luki Goldschmidt for supporting the running of the Institute for Protein Design.

This work was supported by gifts from Microsoft (D.L.S., D.B.), The Donald and Jo Anne Petersen Endowment for Accelerating Advancements in Alzheimer’s Disease Research (N.R.B.), Amgen (J.L.W.), grant DE-SC0018940 MOD03 from the U.S. Department of Energy Office of Science (A.J.B., D.B.), the National Institute of General Medical Sciences of the National Institutes of Health under Award Number T32GM008268 (D.L.S.), the National Eye Institute of the National Institutes of Health under Award Number T32EY032448 (Y.Y.), grant 5U19AG065156-02 from the National Institute for Aging (D.B.), grant R01CA260415 from the National Cancer Institute (C.C.L.), grant R35GM136297 from the National Institute of General Medical Sciences (C.C.L.), the Institute for Rapid Antibody Engineering and Evolution as part of the Engineering+Health Initiative of the UCI Samueli School of Engineering (C.C.L.), the Open Philanthropy Project Improving Protein Design Fund (R.J.R., D.B.), a grant (INV-010680) from the Bill and Melinda Gates Foundation (J.L.W. C.W., E.L.S., K.D.C., D.B.), an EMBO Postdoctoral Fellowship (grant number ALTF 292-2022; J.L.W.), Howard Hughes Medical Institute COVID-19 Initiative (C.E.W.), Defense Threat Reduction Agency grant HDTRA1-21-1-0007 (B.H.), a National Science Foundation Training Grant (EF-2021552; P.J.Y.L.), NERSC award BER-ERCAP0022018 (P.J.Y.L.), a Grants for Resident Innovation and Projects award from the Children’s Hospital of Philadelphia (R.A.), as part of the NexTGen team supported by the Cancer Grand Challenges partnership funded by Cancer Research UK (CGCATF-2021/100002), the National Cancer Institute (CA278687-01) and The Mark Foundation for Cancer Research (J.M.M, N.G.S.), a grant (U19 AG065156) from the National Institute for Aging (S.V.T.), Washington Research Foundation Postdoctoral Fellowship program (R.J.R.), Defense Threat Reduction Agency Grant HDTRA1-21-1-0038 (I.G.), the Howard Hughes Medical Institute (N.R.B., R.J.R., D.B.), a grant from the Institute for Basic Science IBS-R030-C1 (H.M.K.), the Bill and Melinda Gates Foundation for Adjuvant Research (C.C), the Audacious Project Project at the Institute for Protein Design (K.D.C., D.B.)

## Author Contributions

N.R.B., J.L.W. and R.J.R conceived the study, and may change the order of their names for personal pursuits to best suit their own interests. N.R.B. and J.L.W. trained RFdiffusion and fine-tuned RoseTTAFold2. R.J.R., D.L.S., and R.B led the experimental work, with help from E.L.S., P.J.Y.L., B.H., I.G., R.A., S.V.T., S.M.S., T.S., and K.O.. A.J.B. led the nsEM and cryo-EM structural characterization work, with help from C.W. and K.D.C.. J.L.W., N.R.B, D.L.S., R.A., C.C., H.M.K. made designs. D.L.S. and B.S. contributed additional code. D.L.S, R.B. and R.J.R. did the retrospective AF3 analysis. J.L.W., R.J.R. and B.I.M.W. devised the library assembly strategy. S.C. and Y.S. purified target proteins. Y.Y. performed the OrthoRep experiments under C.C.L.’s guidance and supervision. J.L.W., R.J.R. and D.B. co-managed the project. J.L.W., D.B., R.J.R., N.R.B. and A.J.B. wrote the manuscript. All authors read and contributed to the manuscript.

## Extended Data Figures

**Extended Data Figure 1:**
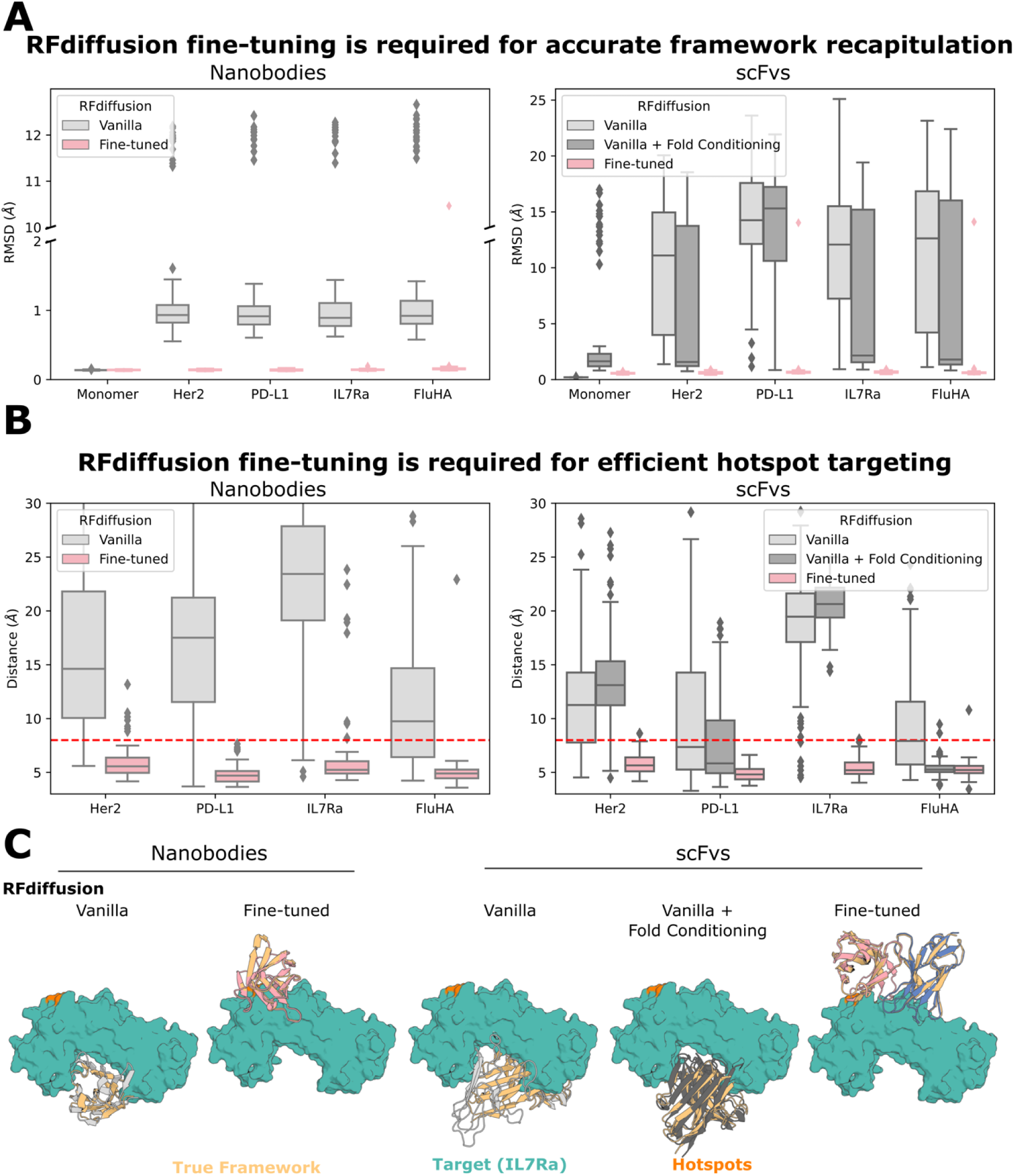
Fine-tuning is required for antibody design with RFdiffusion. **A**) To test whether existing vanilla RFdiffusion models were capable of designing VHHs/scFvs, we explored means of providing the antibody template. For VHHs (left), we used RFdiffusion variant trained to condition on sequence alone^10^ and provided the VHH framework sequence (gray). This version, as compared to the fine-tuned version described in this work (pink), was significantly worse at recapitulating the native VHH framework structure. For scFvs (right), we additionally tried providing fold-level information into the appropriate vanilla RFdiffusion model^9^ (dark gray), but found that this was also insufficient to get accurate recapitulation of the scFv framework. Fine-tuning (pink) yields near-perfect recapitulation of the scFv framework structure. **B**) Although vanilla RFdiffusion is trained to respect “hotspots”, for VHHs (left) and scFvs (right) we find this to be less robust (grays) than after fine-tuning on antibody design (pink). **C**) Examples depicting the results of (**A**) and (**B**). In all cases, the “median” accuracy example (by framework recapitulation) is shown. Left to right: i) without fine-tuning, vanilla RFdiffusion does not target “hotspot” residues (orange) effectively, and does not recapitulate the VHH framework accurately (gray vs yellow). ii) After fine-tuning on antibody design, RFdiffusion targets “hotspots” with accurately recapitulated VHHs (pink vs yellow). iii) Providing only the scFv sequence, vanilla RFdiffusion does not target “hotspots” (orange) accurately nor accurately recapitulates the VHH framework (gray vs yellow). iv) Providing additional fold-level information is insufficient to get perfect framework recapitulation (dark gray vs yellow). v) After fine-tuning on antibody design, RFdiffusion can design scFvs with accurate framework structures (blue/pink vs gray) targeting the input “hotspots” (orange).

**Extended Data Figure 2:**
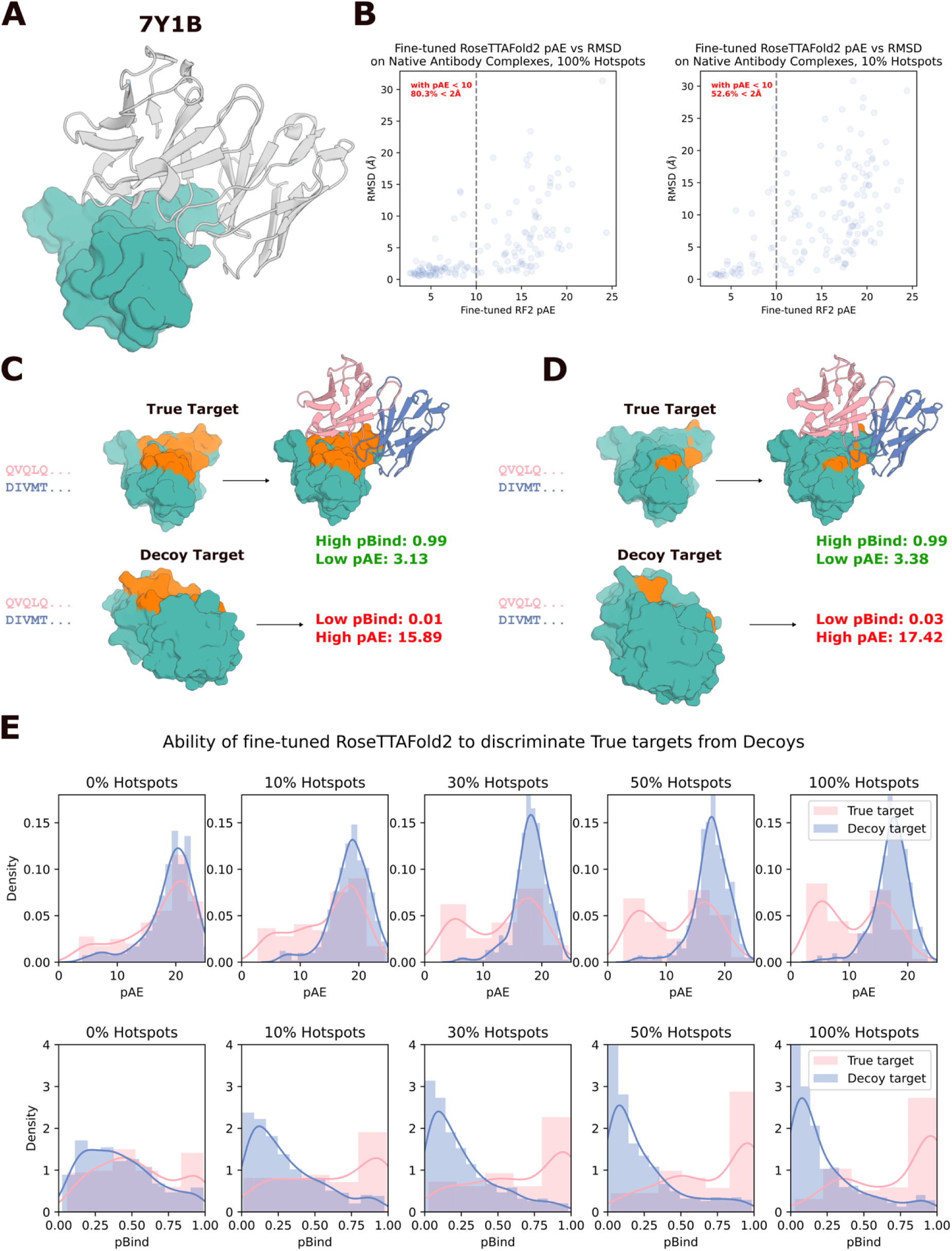
Fine-tuned RoseTTAFold2 can distinguish true complexes from decoy complexes. **A**) An example antibody structure from the validation set used in this figure, which shares < 30% sequence similarity on the target (teal) to anything in the RoseTTAFold2 fine-tuning training dataset. **B**) Fine-tuned RoseTTAFold2 quite reliably predicts its own accuracy. Correlation between RF2 predicted aligned error (pAE) and RMSD to the native structure with 100% (left) or 10% (right) of “hotspot” residues provided. With pAE < 10, 80.3% of structures are within 2 Å when 100% of “hospots” are provided (along with the holo target structure), with this falling to 52.6% when only 10% of hotspots are provided. **C**-**D**) Cherry-picked example of RoseTTAFold2 correctly distinguishing a “true” from a “decoy” complex. The sequence of antibody 7Y1B was provided either with the correct (PDB ID: 7Y1B) or decoy (PDB ID: 8CAF) target. Both with 100% (**C**) or 10% (**D**) of “hotspots” provided, RF2 near-perfectly predicts binding (top row) or non-binding (bottom row). **E**) Quantification of the fine-tuned RF2’s ability to distinguish true targets from decoy targets with both pAE (top row) and pBind (bottom row). Note that this ability depends on the proportion of “hotspots” provided. Without any “hotspots” provided, RF2 is hardly predictive, because RF2 without privileged information is quite rarely confident or accurate in its predictions.

**Extended Data Figure 3:**
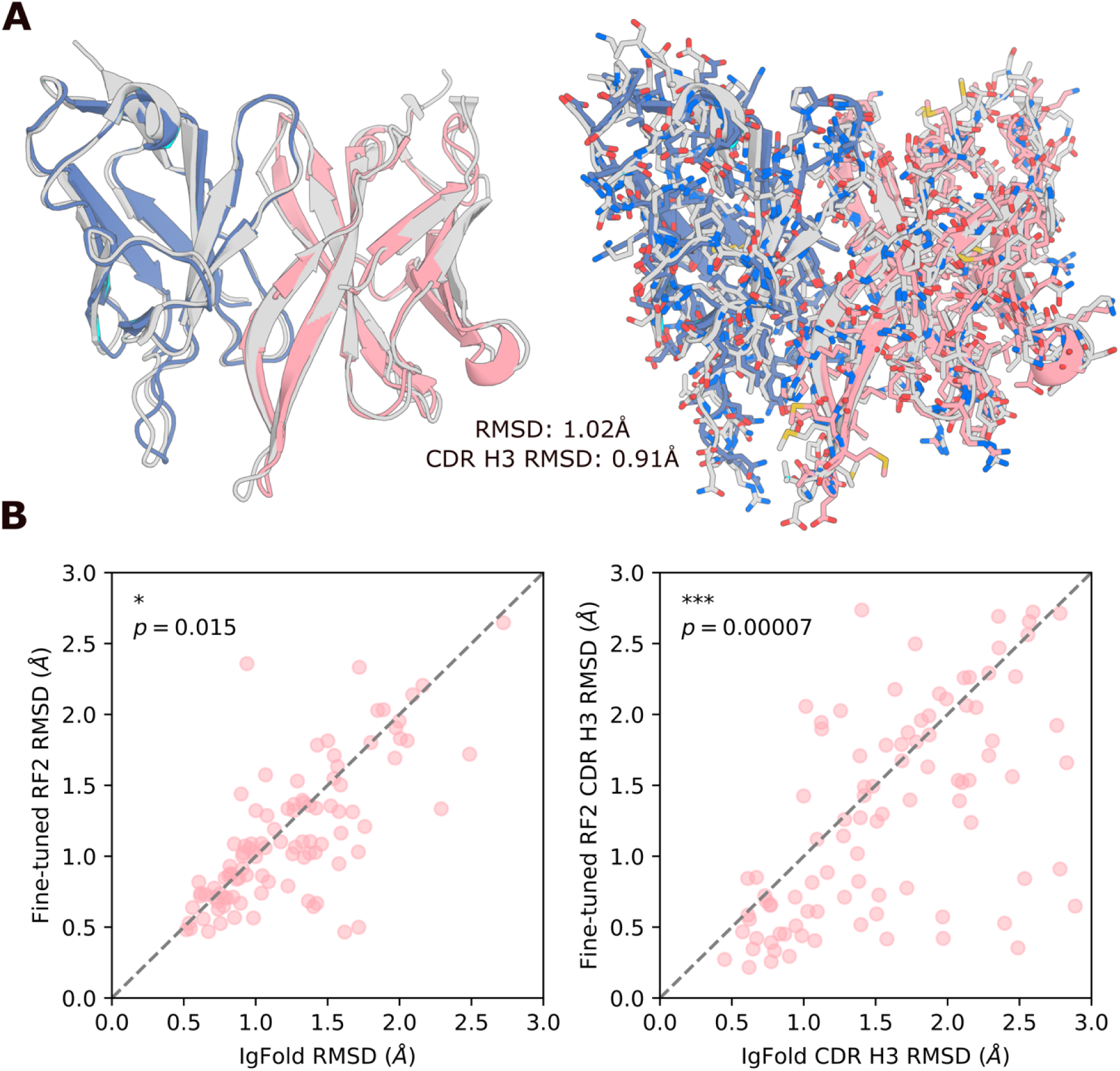
Comparison of fine-tuned RoseTTAFold2 to IgFold on antibody monomer prediction. **A**) 104 antibodies released after the RF2 (and IgFold) training dataset date cutoff (January 13th, 2023) that share < 30% target sequence similarity to any antibody complex released prior to this date were predicted as monomers with either fine-tuned RF2 or IgFold (IgFold cannot predict antibody-target complexes). Shown is the median Fv quality prediction (by overall RMSD) of fine-tuned RF2, of (PDB ID: 8GPG), with (right) and without (left) sidechains shown (gray: native; colors: prediction). While the backbone RMSD is close to the true structure, some sidechains are incorrectly positioned. **B**) Fine-tuned RF2 slightly outperforms IgFold at prediction accuracy. Overall prediction accuracy is slightly improved in fine-tuned RF2 vs IgFold (*p=0.015*, Wilcoxon Paired Test), with greater improvements in CDR H3 prediction accuracy (*p=0.00007*, Wilcoxon Paired Test).

**Extended Data Figure 4:**
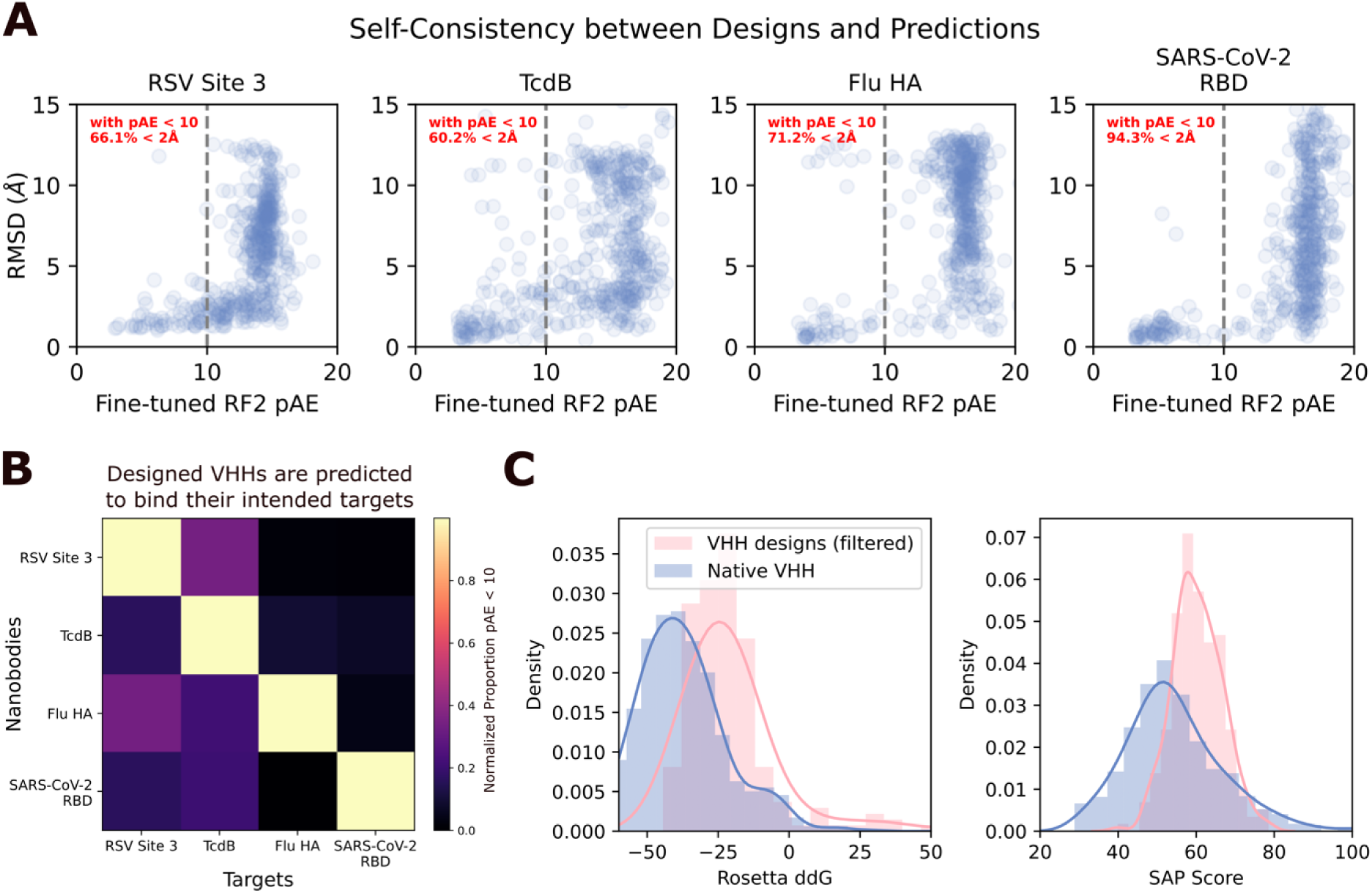
Fine-tuned RoseTTAFold2 recapitulates design structures and computationally demonstrates specificity of VHHs for their targets. **A**) Comparison of RF2 pAE and RMSD of the prediction to the design model. A significant fraction of designs are re-predicted by RF2 (given 100% of “hotspots”), and pAE correlates well with accuracy to the design model. **B**) RF2 can be used to assess quality of designed VHHs. Providing the VHH sequence with the true target structure (used during design) leads to higher rates of high-confidence predictions than predicting the same sequence with a decoy structure (not used in design), as assessed by the fraction of predictions with pAE < 10 (normalized to the fraction of predictions with pAE < 10 for that target with its “correct” VHH partners). In these experiments, the true or decoy target was provided along with 100% of hotspot residues, with those hotspot residues derived from the target with its “true” designed VHH bound. **C**) Orthogonal assessment of designed VHHs with Rosetta demonstrates that the interfaces of RF2-approved (RMSD < 2 Å to design model, pAE < 10) VHH designs have low change in free energy (ddG) (top; only slightly worse than native VHHs) and slightly higher spatial aggregation propensity (SAP) score as compared to natives (bottom).

**Extended Data Figure 5:**
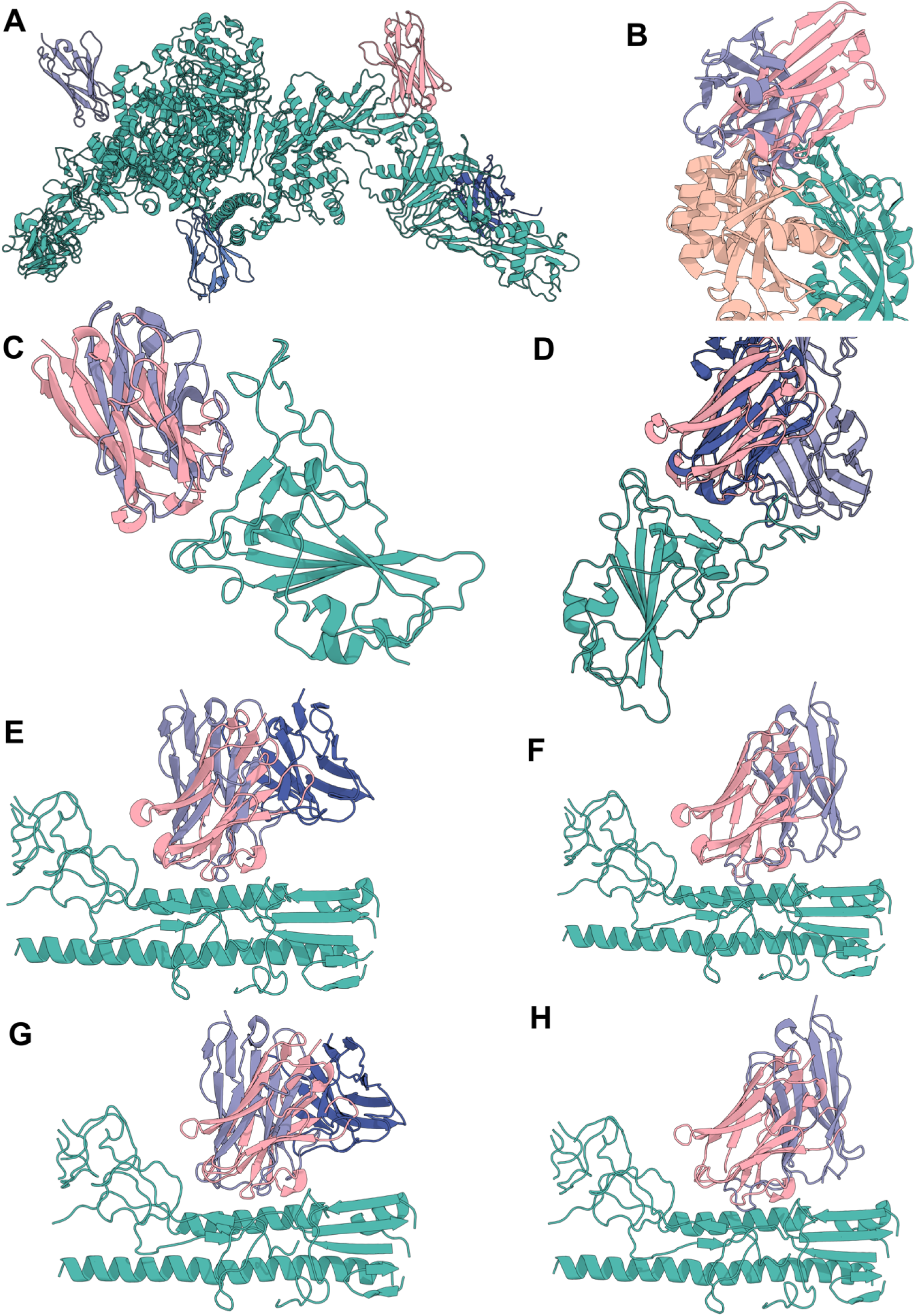
Alignment of VHH Design Models to Complexes in the PDB. For each of the highest affinity VHHs identified for each target, and the structurally characterized influenza HA VHH, the closest complex in the PDB is shown. Designed VHHs (pink) are shown in complex with their designed target (teal and tan). The closest complex was identified visually (Methods). **A**) Designed TcdB VHH aligned against 3 VHHs from(PDB ID: 6OQ5) (shades of blue). The designed TcdB VHH binds to a site for which no antibody or VHH structure exists in the PDB. **B**) Designed RSV Site III VHH aligned against VHH from (PDB ID: 5TOJ) (blue). **C**) Designed SARS-CoV-2 VHH aligned against VHH from (PDB ID: 8Q94) (blue). **D**) Designed SARS-CoV-2 VHH aligned against Fab from 7FCP (shades of blue). **E**) Highest affinity designed influenza HA VHH aligned against Fv from (PDB ID: 8DIU) (shades of blue). **F**) Highest affinity designed influenza HA VHH aligned against VHH from (PDB ID: 6YFT) (blue). **G**) Structurally characterized (cryoEM) designed influenza HA VHH aligned against Fv from (PDB ID: 8DIU) (shades of blue). **H**) Structurally characterized (cryoEM) designed influenza HA VHH aligned against VHH from (PDB ID: 6YFT) (blue).

**Extended Data Figure 6:**
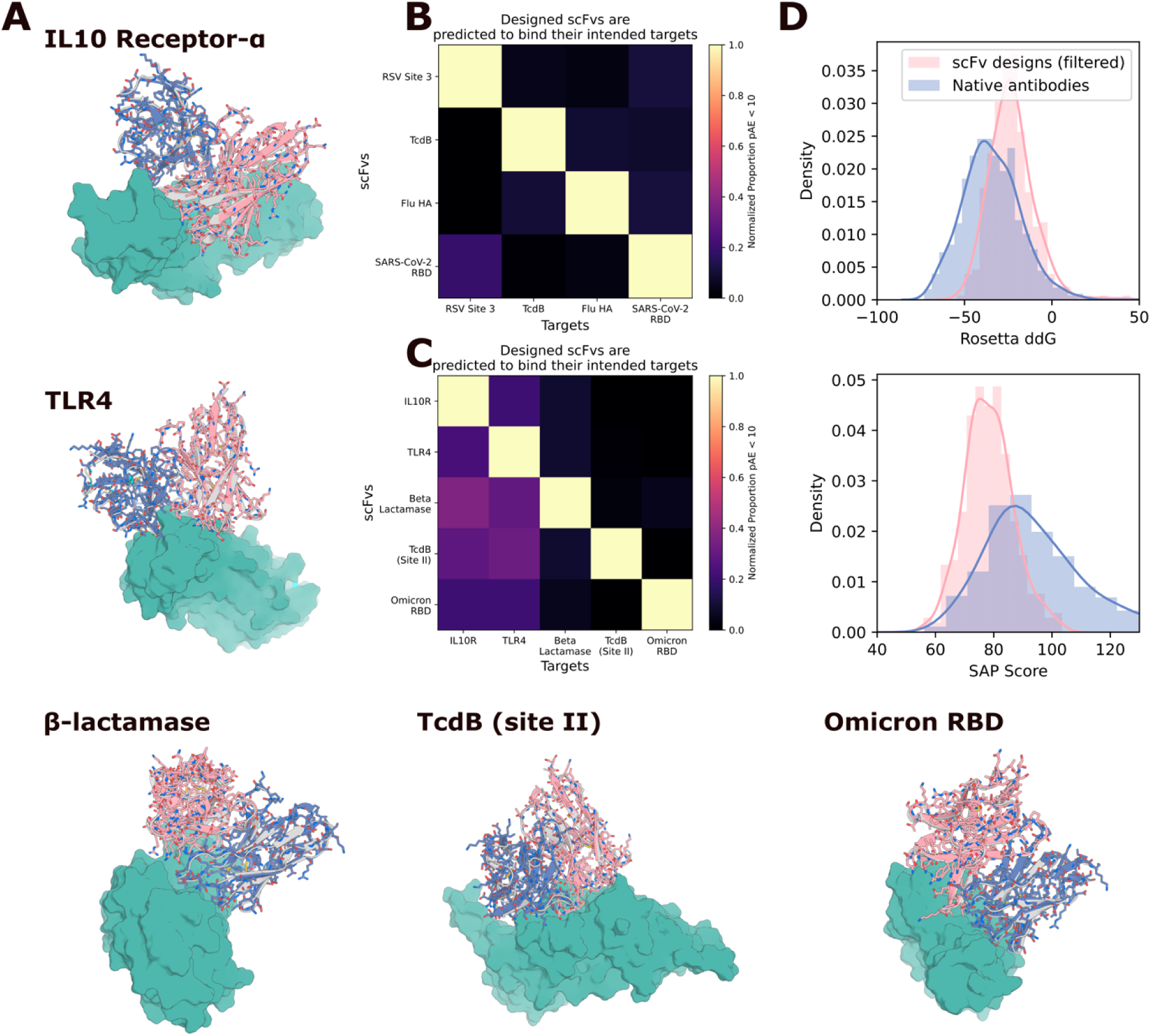
In silico evaluation of RFdiffusion scFv designs. **A**) RFdiffusion was used to generate scFv designs using the framework from Herceptin (hu4D5-8), which has been used to make scFvs previously^57^. Five targets were chosen (IL10 Receptor-ɑ, TLR4, β-lactamase, TcdB and SARS-CoV-2 (omicron) RBD (PDB IDs: 6X93, 4G8A, 4ZAM, 7ML7, 7WPC). Shown are five examples with close agreement between the design model and the fine-tuned RF2 prediction (RMSD (Å): 0.60, 0.56, 0.46, 0.43, 0.61; pAE: 4.73, 4.10, 4.49, 3.52, 3.65). Gray: designs, Pink: RF2 prediction. **B**) Against the four targets to which VHHs were successfully designed, fine-tuned RF2 predicts good specificity to the designed target vs decoy targets. **C**) Against the five targets shown in (**A**), fine-tuned RF2 similarly predicts high specificity to the designed target vs decoy targets. **D**) Orthogonal assessment of designed scFvs with Rosetta demonstrates that the interfaces of RF2-approved (RMSD < 2 Å to design model, pAE < 10) scFv designs have low ddG (top; only slightly worse than native Fabs) and lower spatial aggregation propensity (SAP) score as compared to natives (bottom).

**Extended Data Figure 7:**
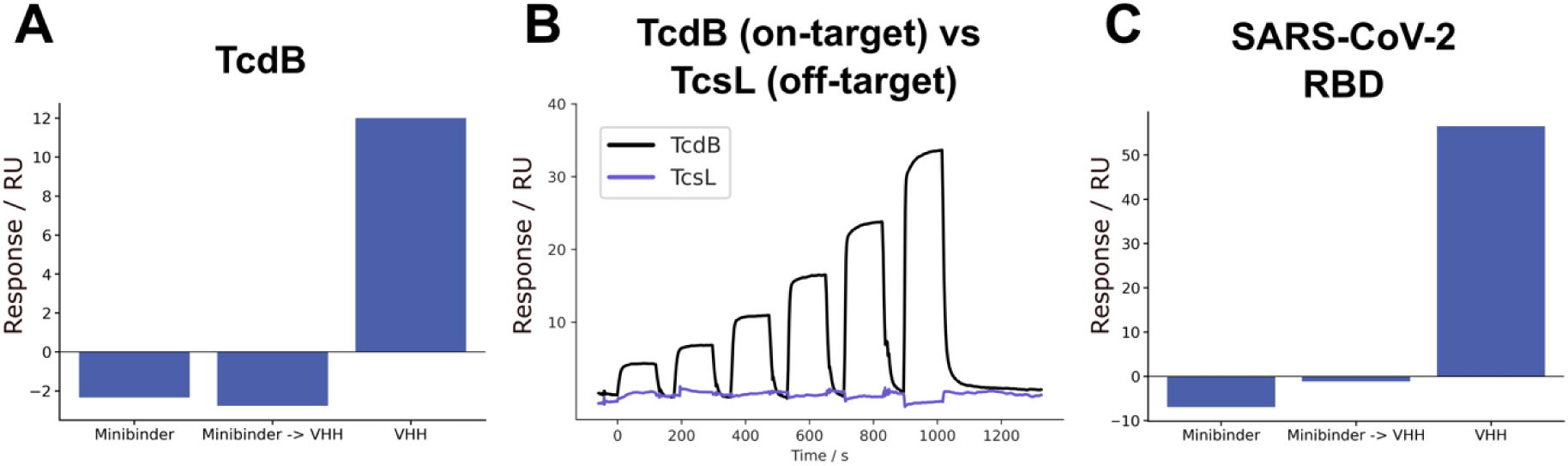
Analysis of SPR Competition Assays. The average response during VHH injection normalized to the response immediately preceding VHH injection for **A**) TcdB VHH competition with Fzd48. **B**) TcdB VHH does not bind to the closely related *Clostridium sordellii* TcsL toxin, indicating that it is binding through specific interactions. **C**) SARS-CoV-2 RBD VHH competition with AHB2. For the competition experiments, in the miniprotein binder-only trace, no VHH is injected and the average response over the corresponding period is plotted as a baseline. (**A**) and (**C**) are the quantification from the rightmost panels of Fig. 2C-D.

**Extended Data Figure 8:**
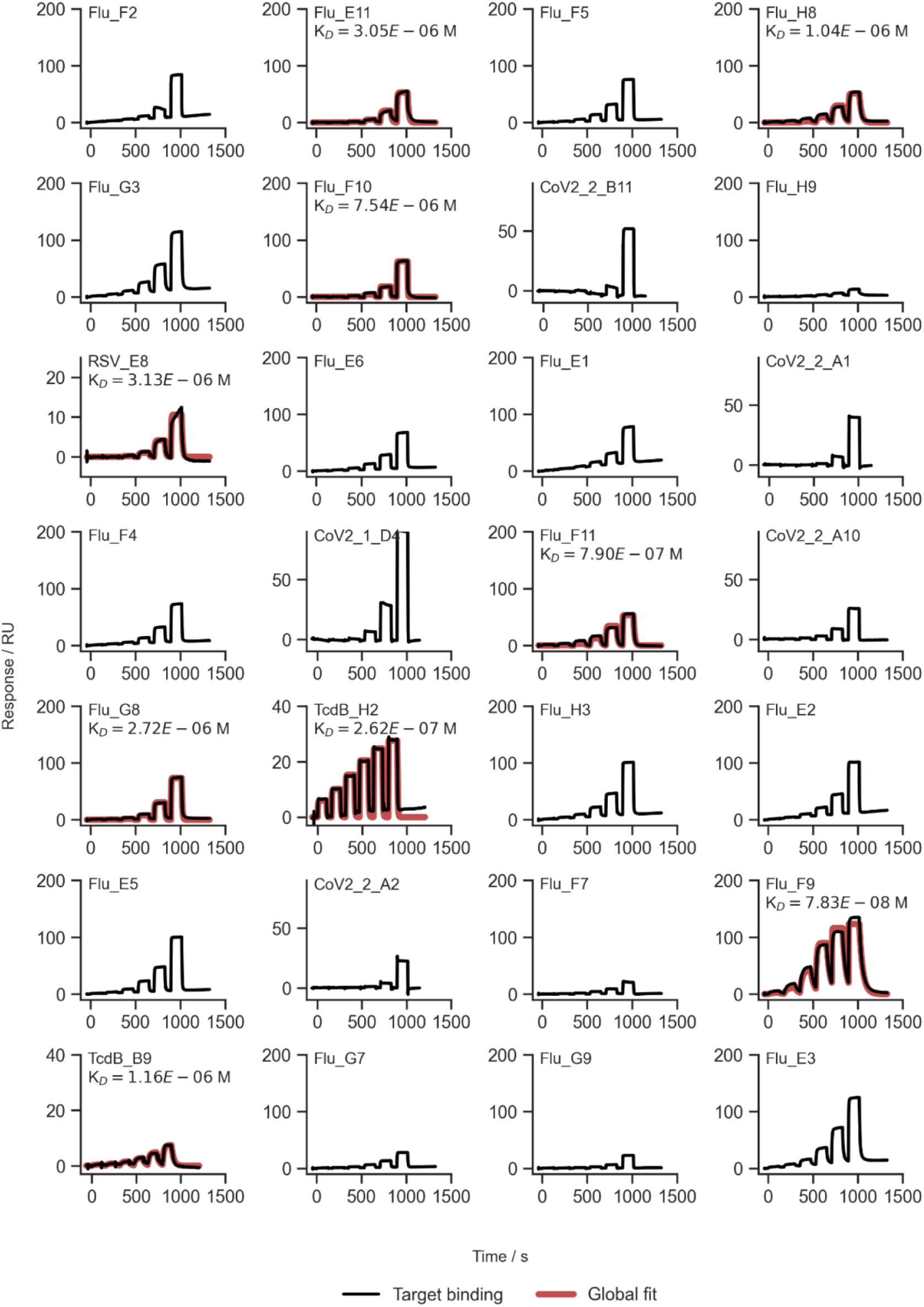
SPR traces of experimentally validated VHHs. SPR traces of the experimentally validated VHH hits described in this study. For traces where confident K_D_ estimates could be fit, we display these on the figure panels.

**Extended Data Figure 9:**
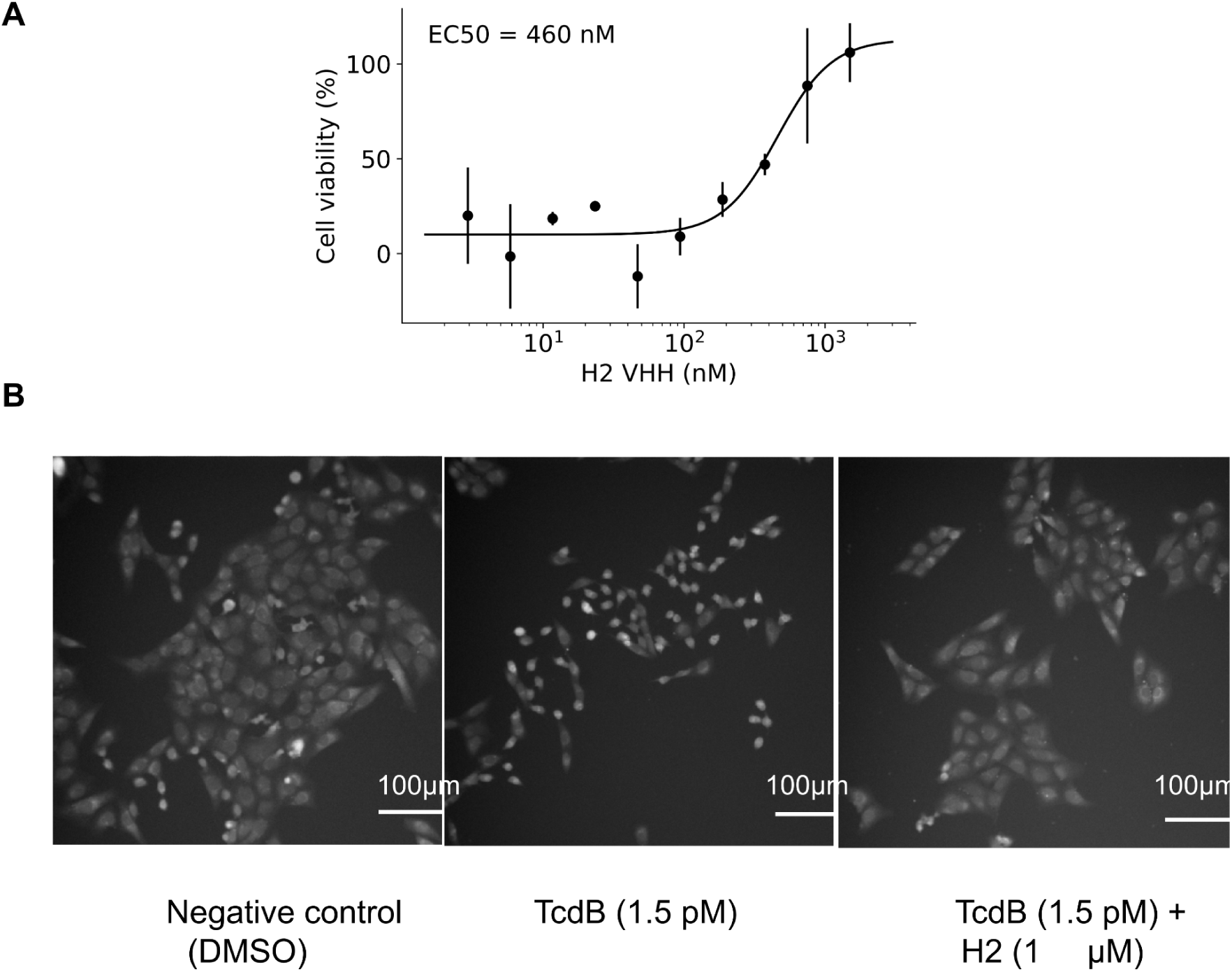
TcdB neutralization by *VHH_TcdB_H2*. **A**) Neutralization of TcdB by *VHH_TcdB_H2* in CSPG4 KO Vero cells. Cell viability is measured after 48 hours. Points indicate the mean and error bars are the standard deviation across two independent replicates. **B**) Vero CSPG4 KO cells treated with vehicle, TcdB alone or TcdB + VHH after 24 hours.

**Extended Data Figure 10:**
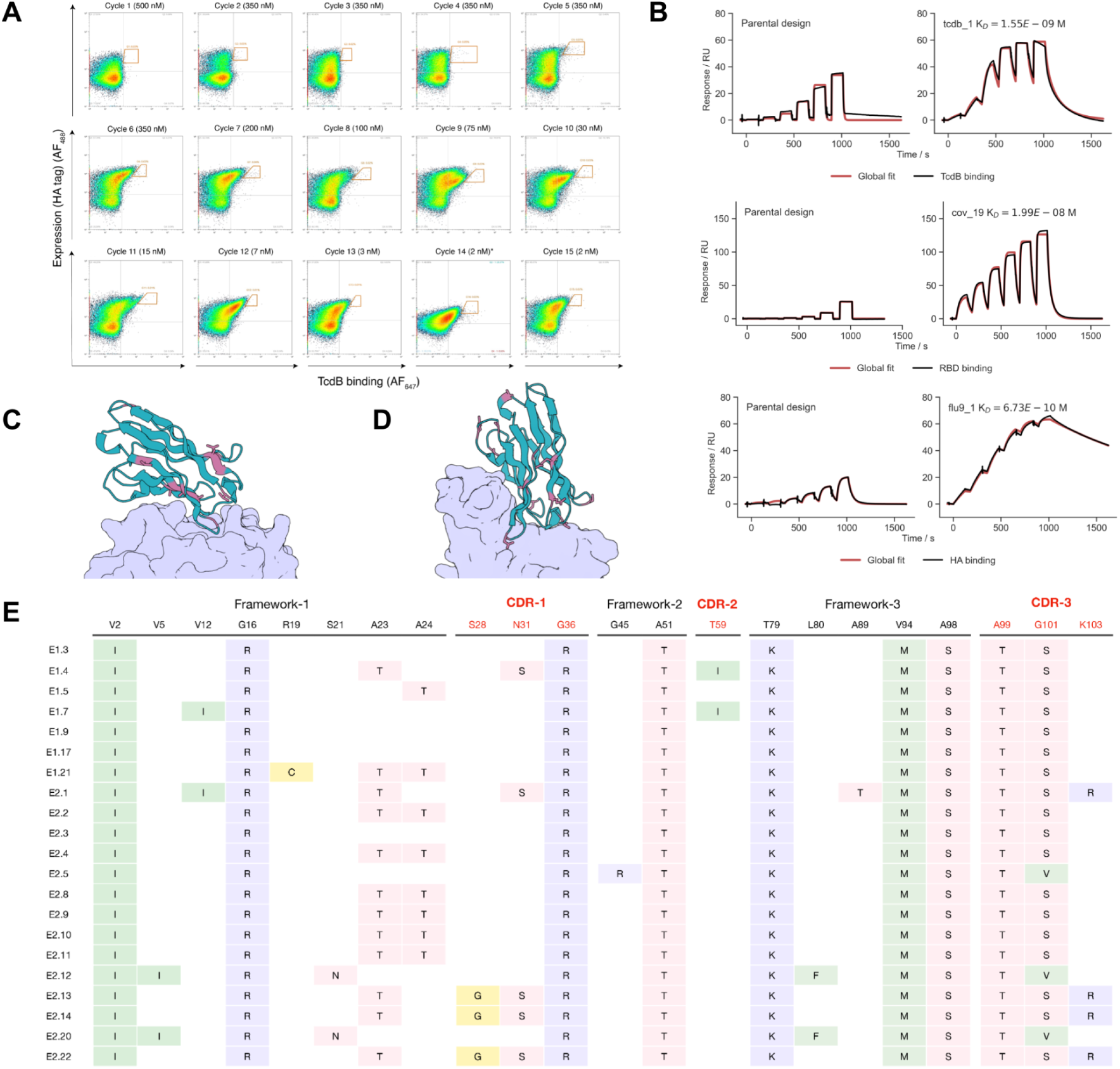
Affinity maturation of VHH binders with OrthoRep. **A**) FACS-plots showing OrthoRep-driven affinity maturation over 15 rounds for a TcdB-targeting VHH where red gates are the population of binders selected for the next round of FACS. The asterisk in the Cycle 14 plot indicates the use of less antibody for detecting expression (Y-axis), explaining why the average expression level of Cycle 14 measures lower than other cycles. FACS plots for OrthoRep-driven affinity maturation campaigns for VHHs targeting SARS-CoV-2 RBD and influenza HA are not shown but follow similar trends. **B**) K_D_ measurements for VHHs affinity-matured against TcdB (first row), SARS-CoV-2 RBD (second row), and influenza HA (third row) in comparison to the parental designs (left column). Dilution series for parental designs (left column) are 5-fold with an upper concentration of 5 μM. Affinity-matured VHHs (right column) were run as titration series as follows: TcdB (200 nM, 4-fold), RBD (200 nM 2-fold) and HA (50 nM, 2-fold). **C-D**) Location of mutations in E2.11 (*TcdB_H2_ortho*), the affinity matured version of *TcdB_H2* (**C**), and *cov_19_ortho* the affinity matured version of the SARS-CoV-2 RBD nanobody *cov_19* (**D**). **E**) Multiple sequence alignment of highly enriched TcdB nanobody clones isolated at cycle 15 of the OrthoRep-driven TcdB nanobody affinity maturation campaign.

**Extended Data Figure 11:**
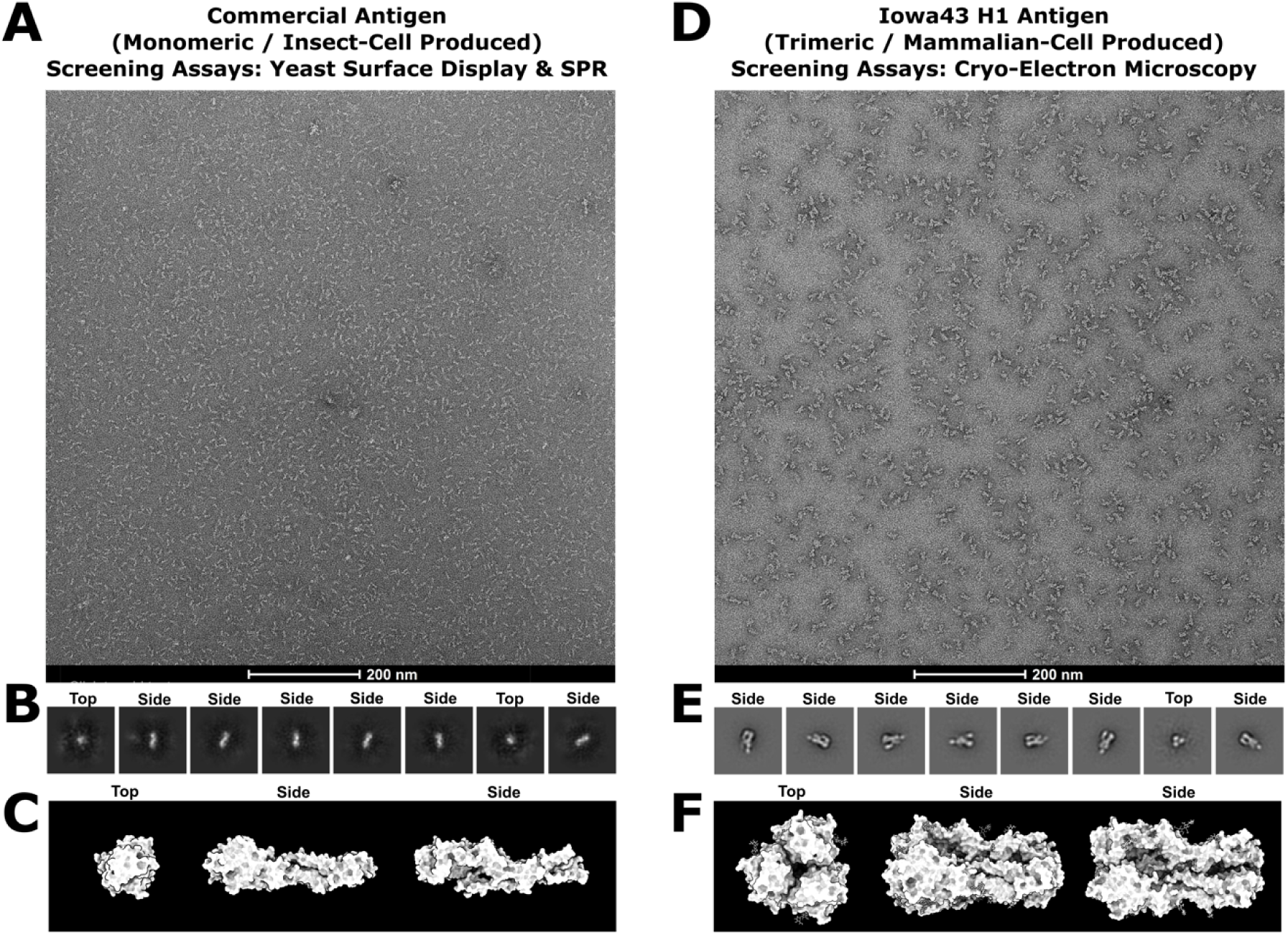
Negative-stain electron microscopy analysis of influenza HA antigens. **A**) Raw nsEM micrograph, **B**) 2D class averages showing a predominance of HA monomer species in the sample, and **C**) a representative predicted 3D model of this commercially produced monomeric HA antigen expressed in insect cells (adapted from PDB ID: 8SK7). This construct was used for screening VHH binders via yeast surface display and surface plasmon resonance. Insect-cell-produced glycoproteins exhibit a truncated glycan shield compared to those produced in mammalian cells. **D**) Raw nsEM micrograph, **E**) 2D class averages showing a clear abundance of HA trimers, and **F**) a representative 3D model of this in-house produced, trimeric Iowa43 HA antigen expressed in mammalian cells (adapted from (PDB ID: 8SK7). This antigen is fully and natively glycosylated, and is the trimeric form of HA. Together these features make Iowa43 suitable for Cryo-EM structural studies of de novo designed VHHs and their capacity to bind to natively glycosylated glycoproteins of therapeutic interest.

**Extended Data Figure 12:**
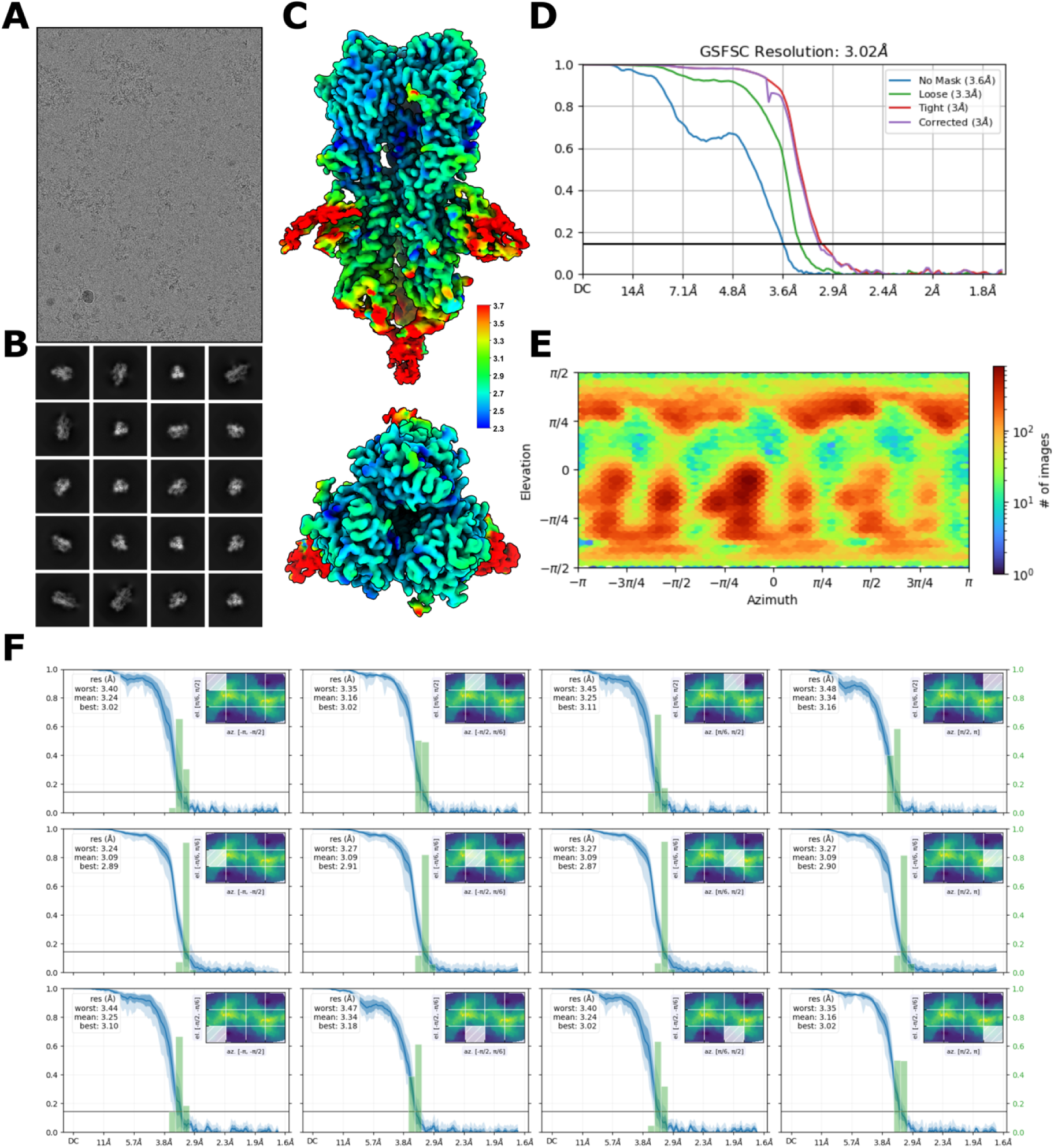
Cryo-EM structure determination statistics for a de novo designed VHH bound to an influenza HA trimer. **A**) Representative raw micrograph showing ideal particle distribution and contrast. **B**) 2D Class averages of Influenza H1+designed VHH with clearly defined secondary structure elements and a full-sampling of particle view angles. **C**) Cryo-EM local resolution map calculated using an FSC value of 0.143 viewed along two different angles. Local resolution estimates range from ∼2.3 Å at the core of H1 to ∼3.7 Å along the periphery of the designed VHH. **D**) Global resolution estimation plot. **E**) Orientational distribution plot demonstrating complete angular sampling.

**Extended Data Figure 13:**
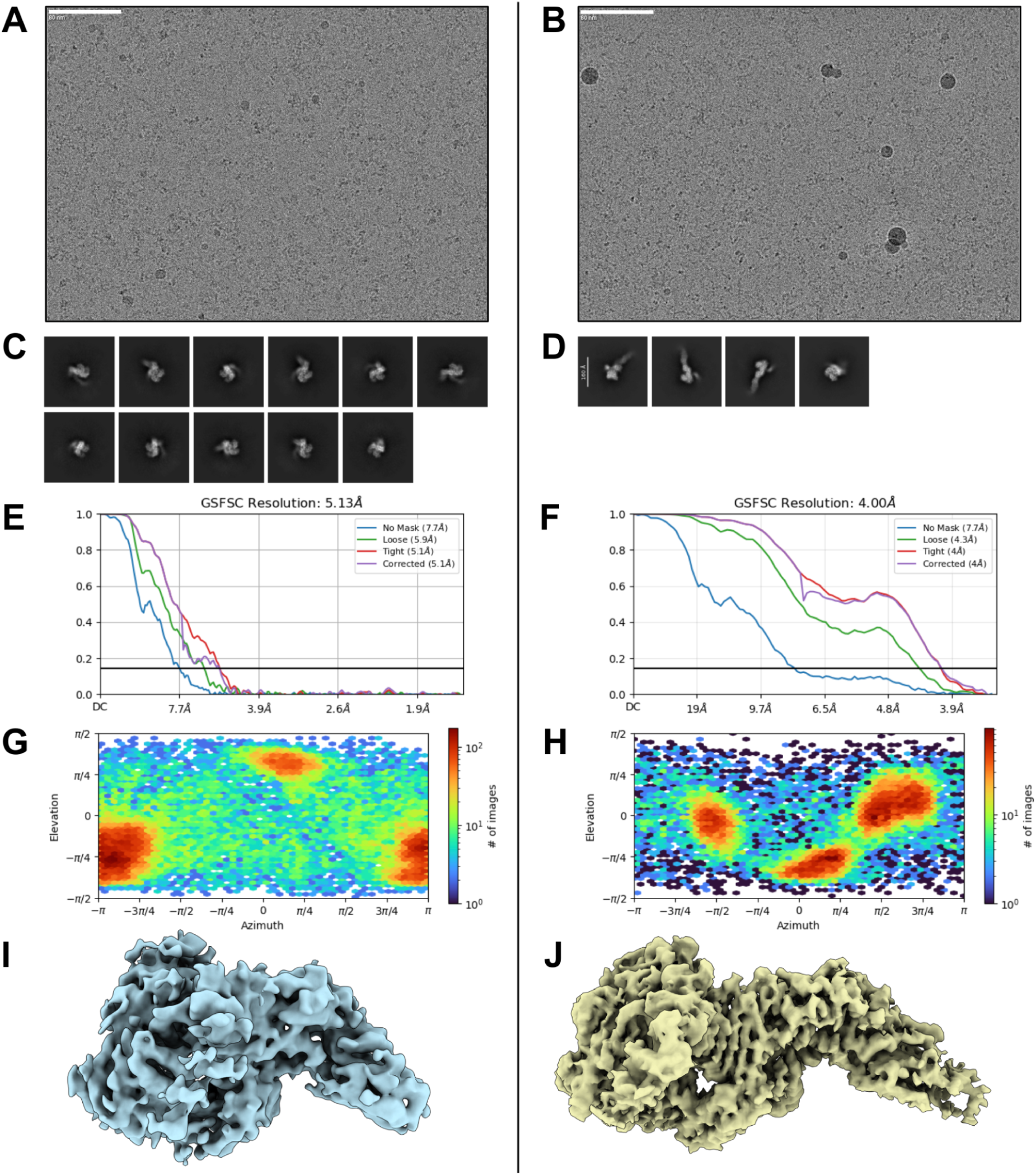
CryoEM statistics for the observed apo states of TcdB, *VHH*_*TcdB_H2*. Early experiments with pre OrthoRep *VHH*_*TcdB_H2* nanobody resulted in 2 apo state structures with no nanobody bound. Representing two states of TcdB observed from two different cryoEM grids collected on Titan Krios with K3 detector; compressed TcdB/ thick ice (**A, C, E, G, I**) 2335 movies and extended TcdB/ thin ice (**B, D, F, H, J**) 3430 movies. **A, B)** Representative Micrographs. **C, D**) 2D Class Averages. **E, F**) Global FSC non uniform refinement. **G, H**) Orientational distribution plots. **I, J**) Sharpened maps, non uniform refinement

**Extended Data Figure 14:**
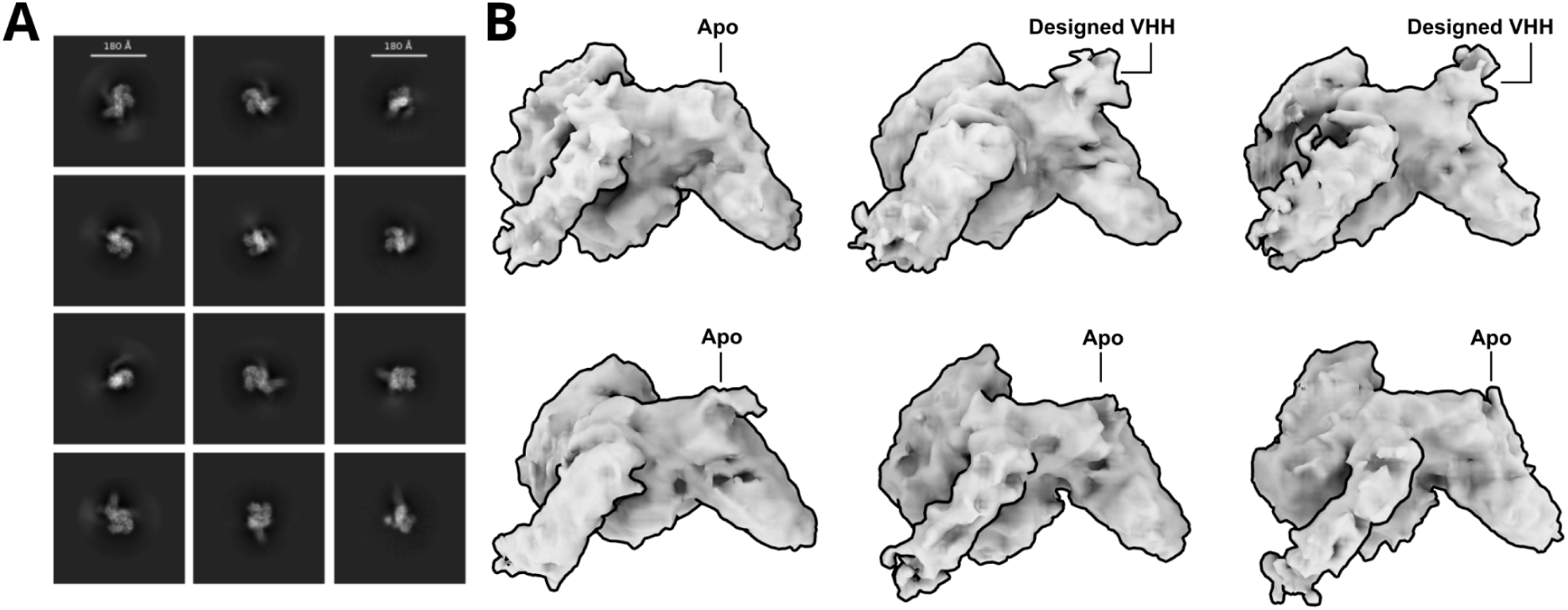
Cryo-EM structural characterization TcdB in complex with a de novo designed VHH, *VHH*_*TcdB_H2* with glycine added. **A)** 2D class averages of TcdB in complex with the de novo designed VHH, *VHH*_*Tcdb_H2*. **B)** 3D classification reveals multiple classes lacking bound density, while two classes exhibit clear density near the Frizzled-7 epitope, the intended target of this design.

**Extended Data Figure 15:**
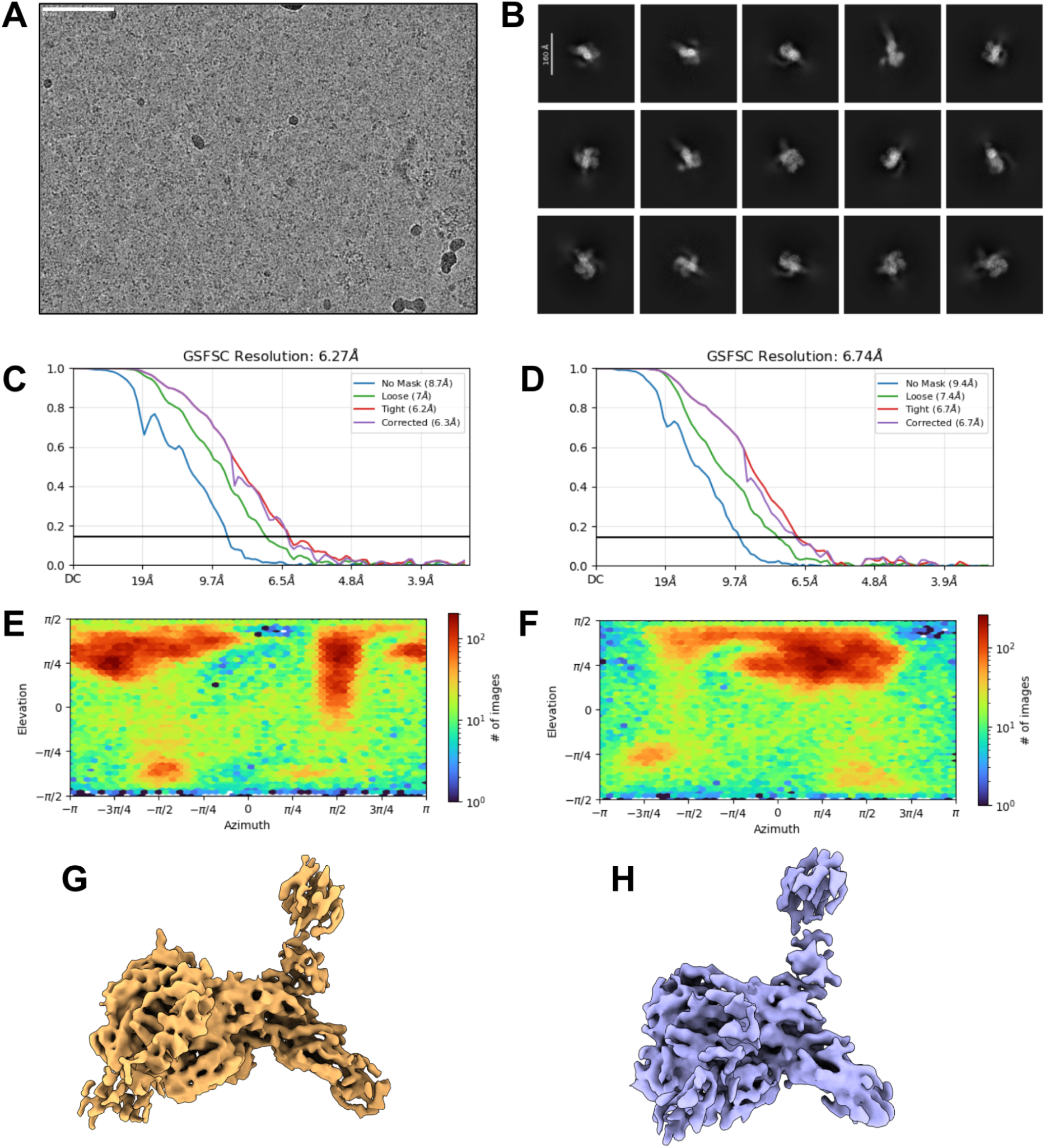
Preliminary CryoEM structures of OrthoRep Affinity Matured TcdB VHH, *VHH*_*TcdB_H2_ortho*; multiple TcdB conformation observed. Thin and thick ice was targeted on one grid. 7,546 movies collected on Titan Krios with K3 detector. 4 major classes were observed and sorted with heterogeneous refinement and previously solved apo state structures: compressed TcdB apo unbound 43,943 particles (not shown), compressed TcdB VHH bound 76,997 particles (**C, E, G**), TcdB extended apo unbound 16,074 particles (not shown), TcdB extended VHH bound 72,014 particles (**D, F, H**) **A**) Representative micrographs **B**) 2D Class Averages. **C, D**) Global Fourier Shell Correlation plot, Non Uniform Refinement **E, F)** Orientational distribution plots. **G, H**) Sharpened maps, non uniform refinement (aggregation from His tag visible).

**Extended Data Figure 16:**
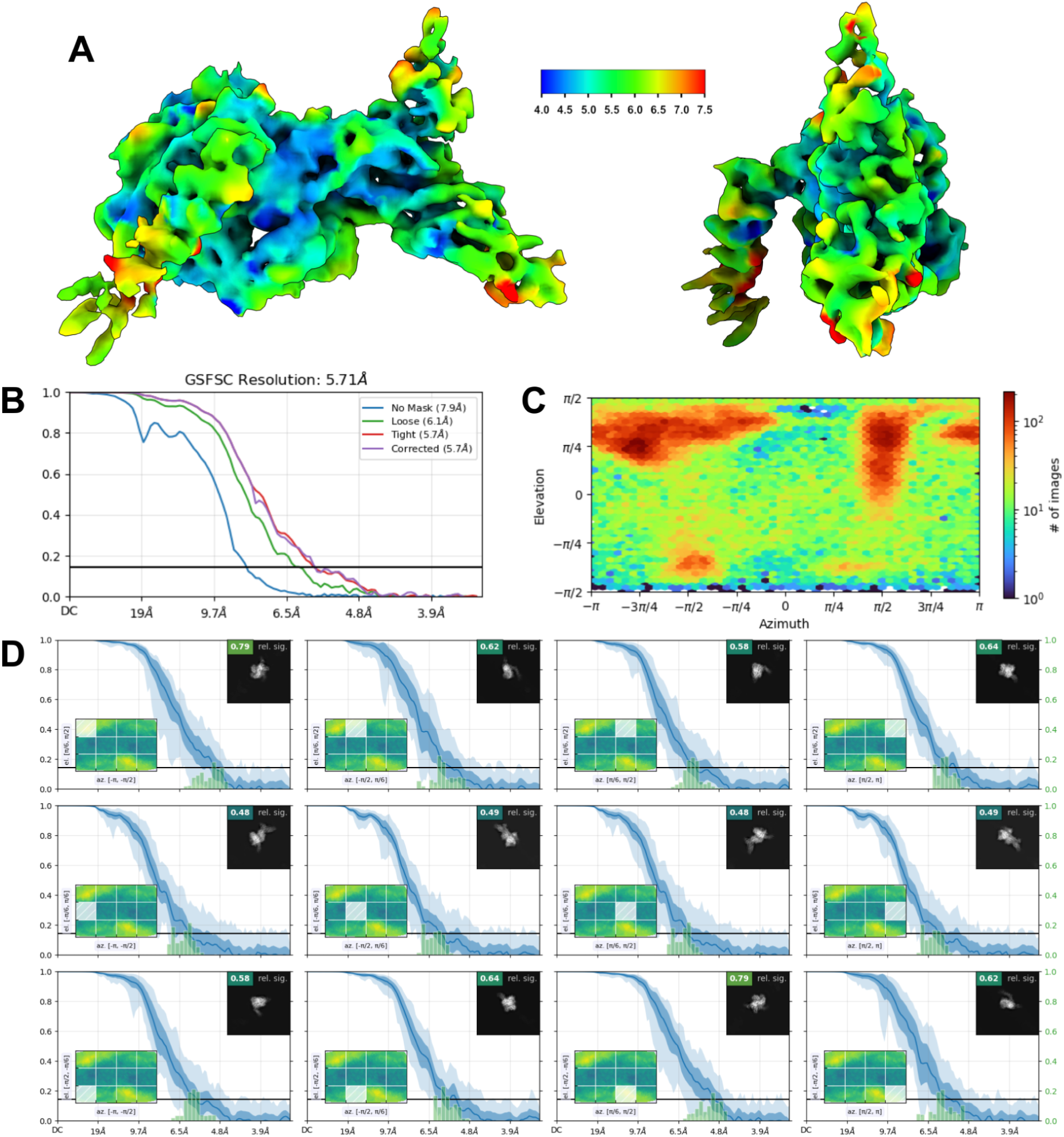
Final Local Refinement CryoEM statistics for OrthoRep Affinity Matured TcdB VHH, *VHH*_*TcdB_H2_ortho* in complex with TcdB. Local refinement and masking used to reduce noise from His tag and improve resolution. **A**) Local Resolution map (Å), calculated using an FSC value of 0.143 viewed along two different angles **B**) Global Fourier Shell Correlation plot, Local Refinement **C**) Orientational distribution plot **D**) Orientational diagnostics data.

**Extended Data Figure 17:**
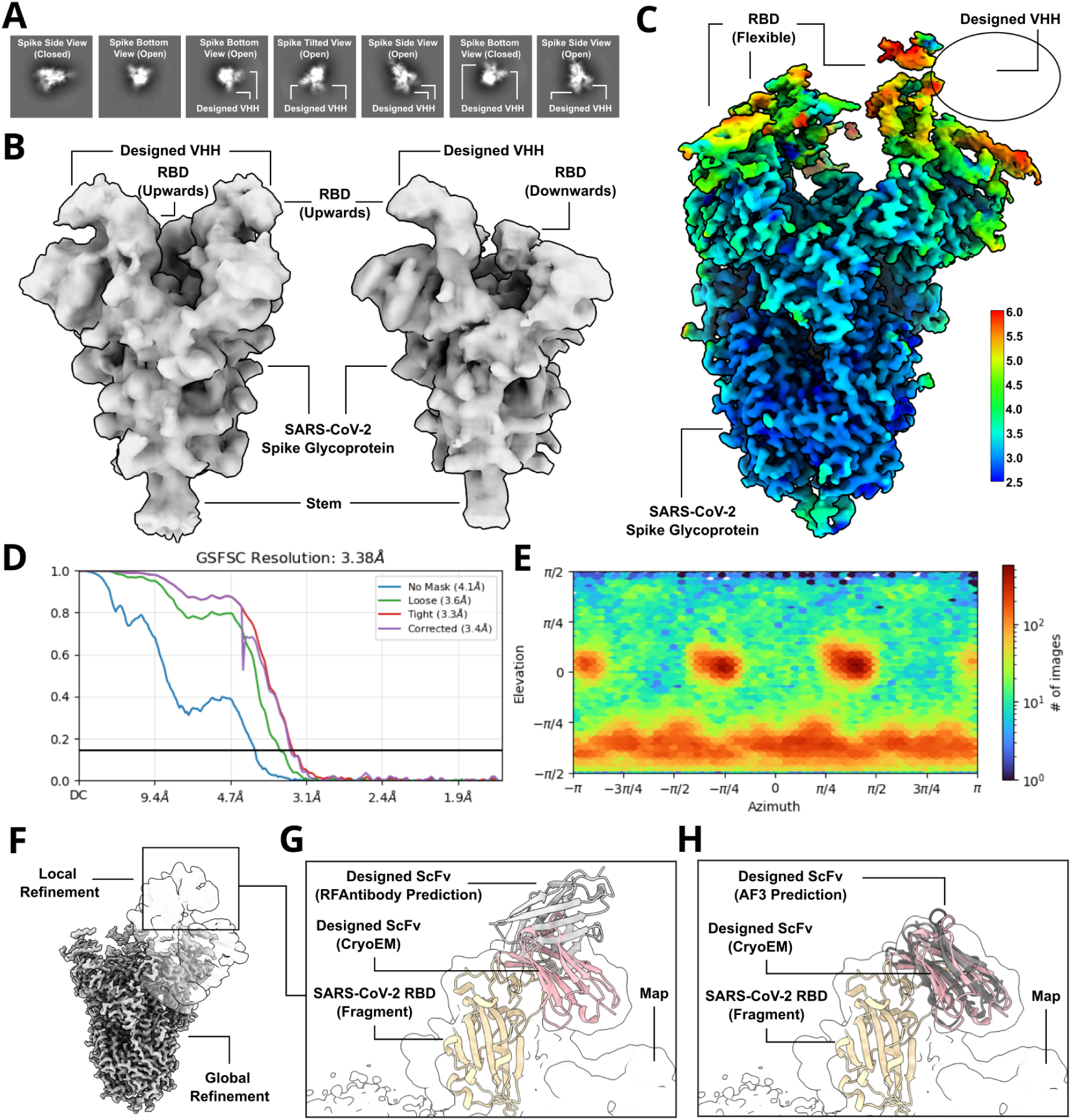
Cryo-EM structural characterization of an inaccurately designed VHH against SARS-CoV-2. **A)** Representative 2D class averages of the designed VHH, *cov_19*, bound to the SARS-CoV-2 RBD. **B**) 3D class averages reveal the SARS-CoV-2 spike population adopting conformations with either one or two RBDs in the up position, with additional density corresponding to the designed VHH. **C**) A globally refined ∼3.4 Å cryo-EM reconstruction of the complex highlights substantial flexibility in the RBDs, leading to poor resolution in these regions and preventing direct visualization of the designed VHH. **D**) Global Fourier shell correlation (FSC) plot of the 3.38 Å cryo-EM map. **E**) Orientational distribution plot showing complete angular sampling of the SARS-CoV-2/designed VHH complex. **F**) Symmetry expansion, 3D classification, and local refinement focused on the RBD region improve the resolution of the bound VHH compared to the globally refined map. (**G**-**H**) Due to the modest resolution of the local refinement, an initial docking of a SARS-CoV-2 RBD fragment into the cryo-EM density was performed, followed by alignment of the full design model—including both the RBD fragment and the designed VHH—to the pre-fitted RBD. This approach confirmed that the RFdiffusion design closely matches the experimentally determined complex in both structure and epitope targeting; however, the binding angle deviates significantly from the design prediction (**G**). In contrast, a retrospective analysis revealed that the corresponding AlphaFold3 (AF3) prediction accurately recapitulated the docking within the cryo-EM density, suggesting that AF3 outperforms RFdiffusion as a predictor of successful binding and should be considered as an additional filter in future design efforts (**F**).

**Extended Data Figure 18:**
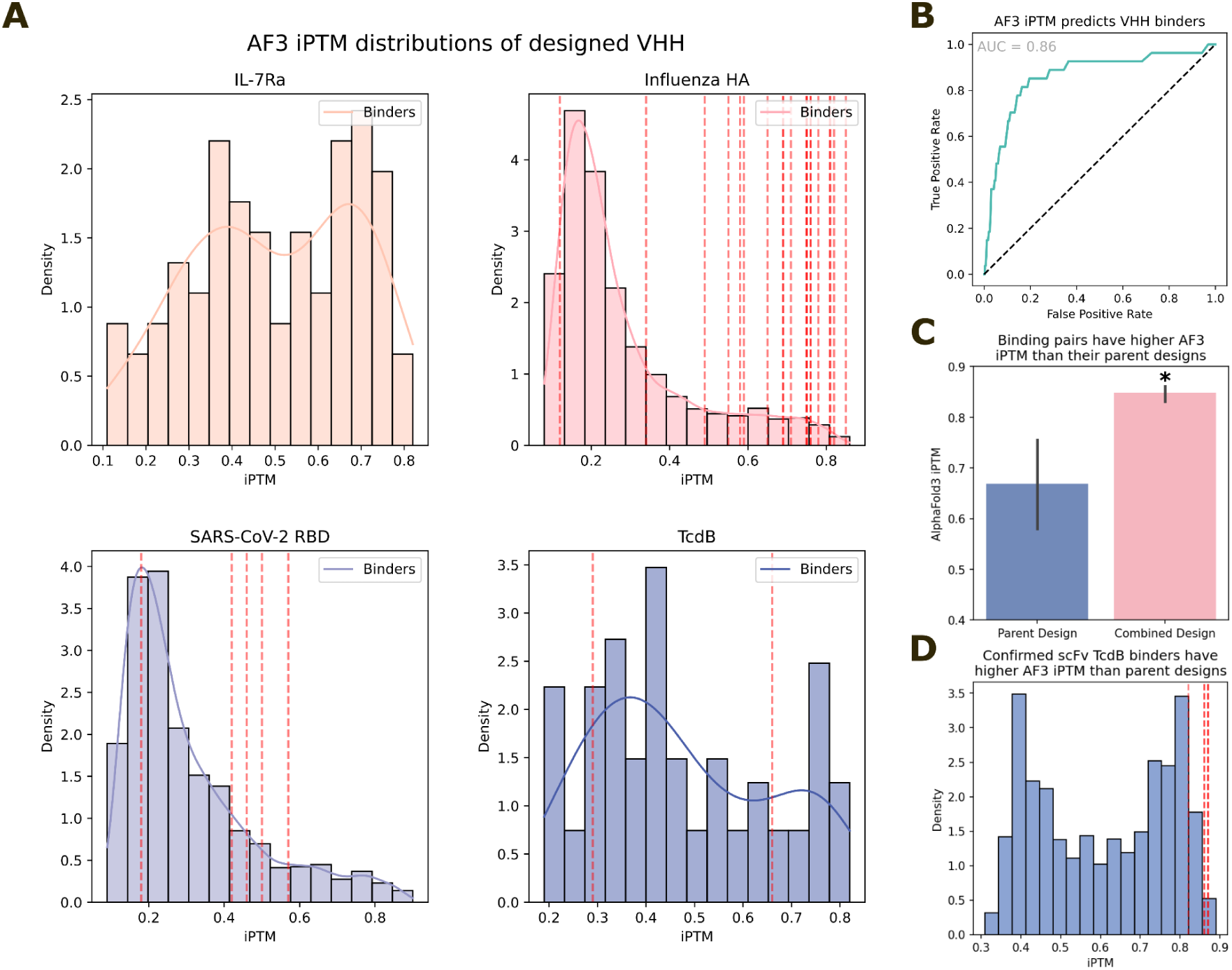
AlphaFold3 retrospectively predicts binders. **A)** iPTM distributions of design VHH libraries against 4 targets. Red lines indicate validated binders. **B**) ROC curve demonstrating strong retrospective predictive power of AF3 at discriminating designed VHH binders from non-binders (AUC=0.86). Note though that this plot is dominated by influenza HA binders, which are more numerous than confirmed binders to SARS-CoV-2 RBD and TcdB. **C**-**D**) Similar retrospective analyses of TcdB scFv binders. These binders were assembled combinatorially from structurally similar “parent” designs. The successfully-binding combined designs have significantly higher AF3 iPTM scores than the parent designs that they emanate from (**C**, 6 binders from 12 parent designs; two-sided Students t-test; *p=0.0025*), and from the parental library as a whole (**D**). These analyses further indicate the utility of AF3 for antibody design filtering.

**Extended Data Figure 19:**
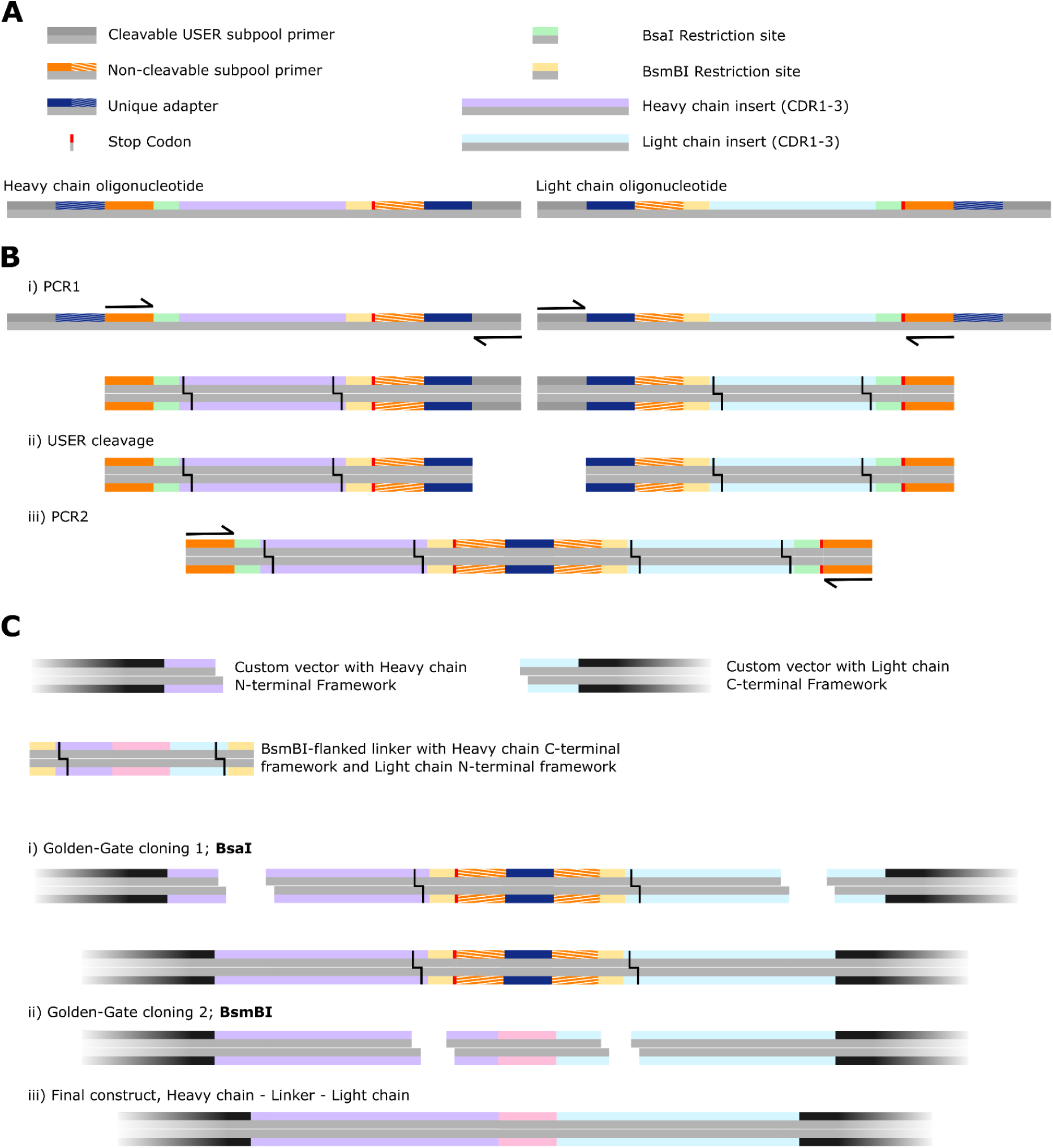
Assembly of uniquely-paired H-L scFvs from 400 bp oligonucleotides. Overview of the assembly strategy for scFvs, uniquely pairing designed heavy chains with their corresponding designed light chain. Note that this strategy permits assembly in either H-L order or L-H order, and for several linkers to be inserted between the chains (see also Extended Data Fig. 15). **A**) Overview of the synthesized oligos. Each heavy-chain oligonucleotide contains two unique “barcodes”, present also on their unique corresponding light-chain oligonucleotide. **B**) **i**) For H-L ordering of the two Fvs, oligonucleotides are amplified such that the 3’ barcode of the heavy chain is amplified (left), and the identical barcode is amplified at the 5’ end of the corresponding light chain. Subpool-specific USER-cleavable primers permit this amplification. **ii**) The USER-cleavable primers are cleaved, exposing the unique barcodes. **iii**) A second PCR is performed with only the outer (non-cleavable) primers. PCR assembly brings together the two designed Fvs over the unique barcode. **C**) **i**) Golden-Gate Assembly (GGA) with BsaI is used to clone these assembled fragments into custom vectors that provide the N-terminal heavy chain framework (up until the start of CDR1) and the C-terminal light chain framework (from the end of CDR3) fragments. Note that GGA is “scarless”, so no additional base pairs are inserted between framework and CDRs. **ii**) After amplification of custom linkers, that are flanked by the C-terminal heavy chain framework (from the end of CDR3) and the N-terminal light chain framework (up until the start of CDR1) fragments, cloning with a second GGA step is performed, with the BsmBI enzyme. Note that multiple different linkers can be included in this second step. Note also the presence of a stop codon after GGA-step 1, to ensure that failure of GGA-step 2 does not affect subsequent functional assays.

**Extended Data Figure 20:**
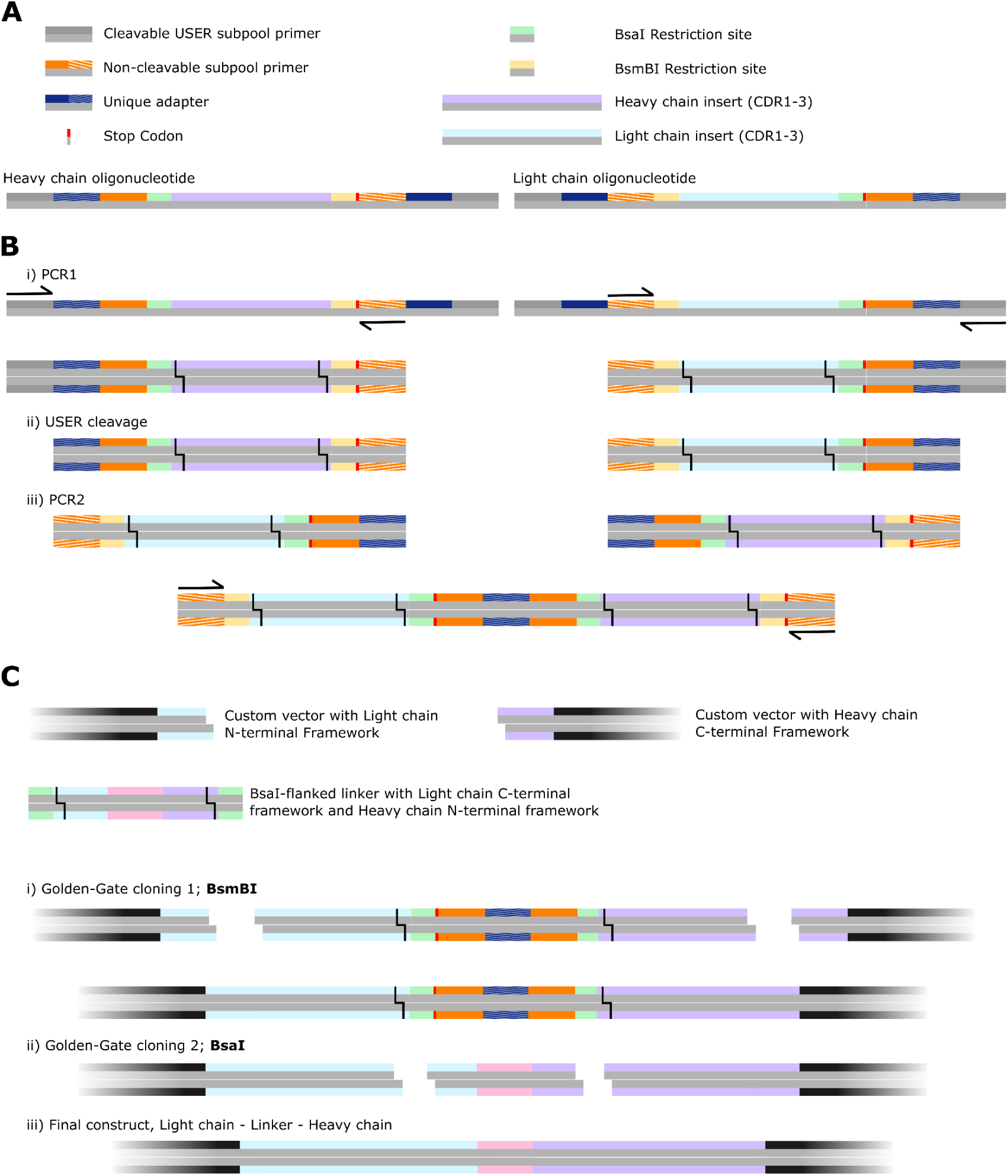
Assembly of uniquely-paired L-H scFvs from 400 bp oligonucleotides. Overview of the assembly strategy for scFvs, uniquely pairing designed heavy chains with their corresponding designed light chain. Note that this strategy permits assembly in either H-L order or L-H order, and for several linkers to be inserted between the chains (see also Extended Data Fig. 14). **A**) Overview of the synthesized oligos. These are the same oligonucleotides as in Extended Data Fig. 14. **B**) **i**) For L-H ordering of the two Fvs, oligonucleotides are amplified such that the 5’ barcode of the heavy chain is amplified (left), and the identical barcode is amplified at the 3’ end of the corresponding light chain. Subpool-specific USER-cleavable primers permit this amplification. **ii**) The USER-cleavable primers are cleaved, exposing the unique barcodes. **iii**) A second PCR is performed with only the outer (non-cleavable) primers. PCR assembly brings together the two designed Fvs over the unique barcode, in L-H ordering. **C**) **i**) Golden-Gate Assembly (GGA) with BsmBI is used to clone these assembled fragments into custom vectors that provide the N-terminal light chain framework (up until the start of CDR1) and the C-terminal heavy chain framework (from the end of CDR3) fragments. Note that GGA is “scarless”, so no additional base pairs are inserted between framework and CDRs. **ii**) After amplification of custom linkers, that are flanked by the C-terminal light chain framework (from the end of CDR3) and the N-terminal heavy chain framework (up until the start of CDR1) fragments, cloning with a second GGA step is performed, with the BsaI enzyme. Note that multiple different linkers can be included in this second step. Note also the presence of a stop codon after GGA-step 1, to ensure that failure of GGA-step 2 does not affect subsequent functional assays.

**Extended Data Figure 21:**
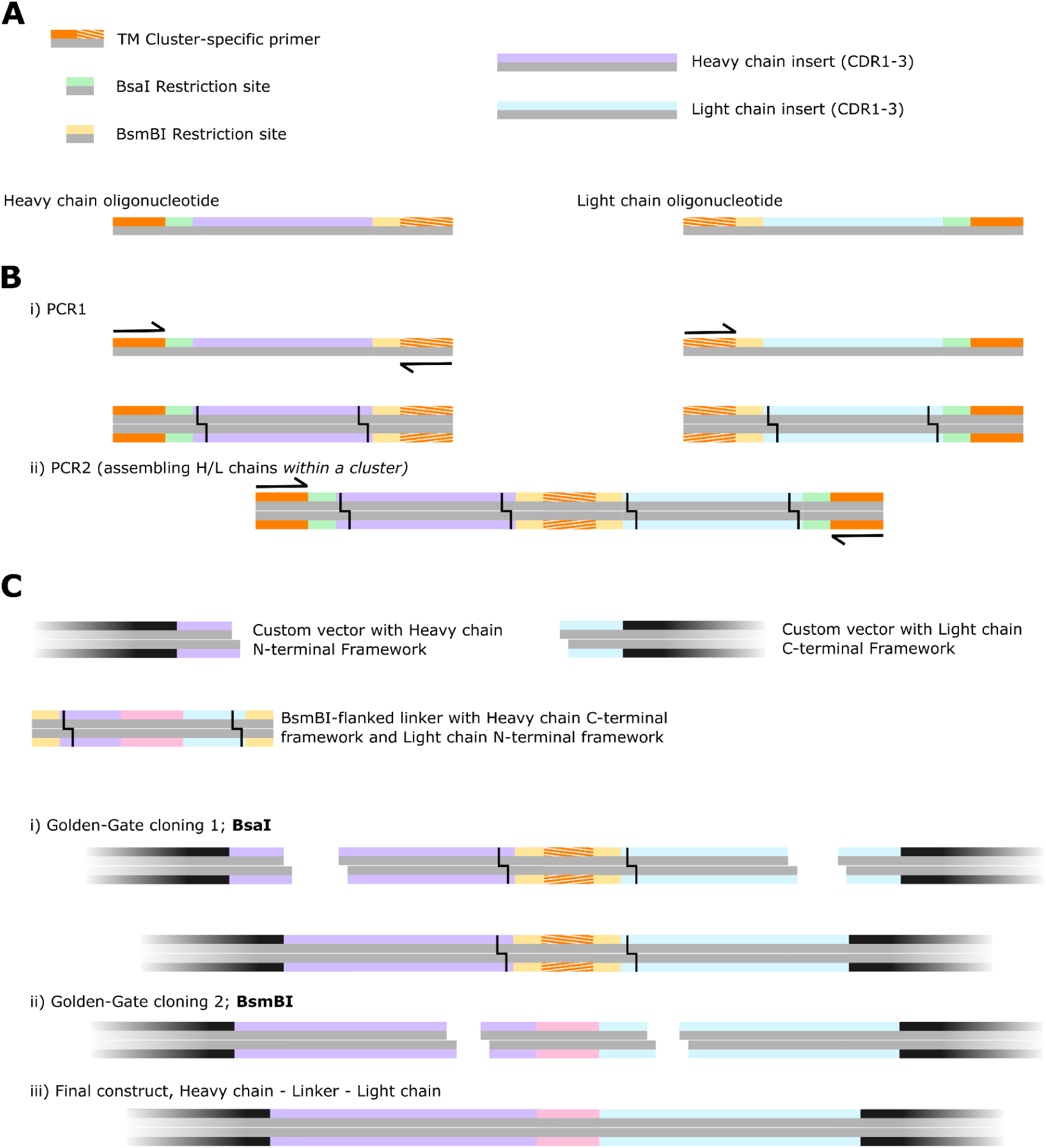
Assembly of structurally-similar clusters of heavy and light chains from 300 bp oligonucleotides. Assembly strategy for combinatorial assembly of structurally-similar, TM-clustered heavy and light chains from 300 bp oligonucleotides. **A**) Schematic of ordered oligonucleotides. Each cluster of designs gets a unique barcode (striped orange). **B**) This cluster-specific barcode is used both as a PCR primer (**i**) and as the basis for PCR assembly (**ii**). These barcodes/primers being unique to each structurally-similar cluster of designs ensures that only “compatible” heavy and light chains assemble. **C**) in an identical manner to Extended Data Figs. 14 & 15, two rounds of GGA ligate the fragment from (**B**) into a custom vector (i) and subsequently ligate in a (set of) linker(s) (ii), to yield a library of structurally-compatible heavy and light chain scFvs. Note that due to oligonucleotide constraints, only one Fv-ordering can be achieved in this 300 bp setup.

**Extended Data Figure 22:**
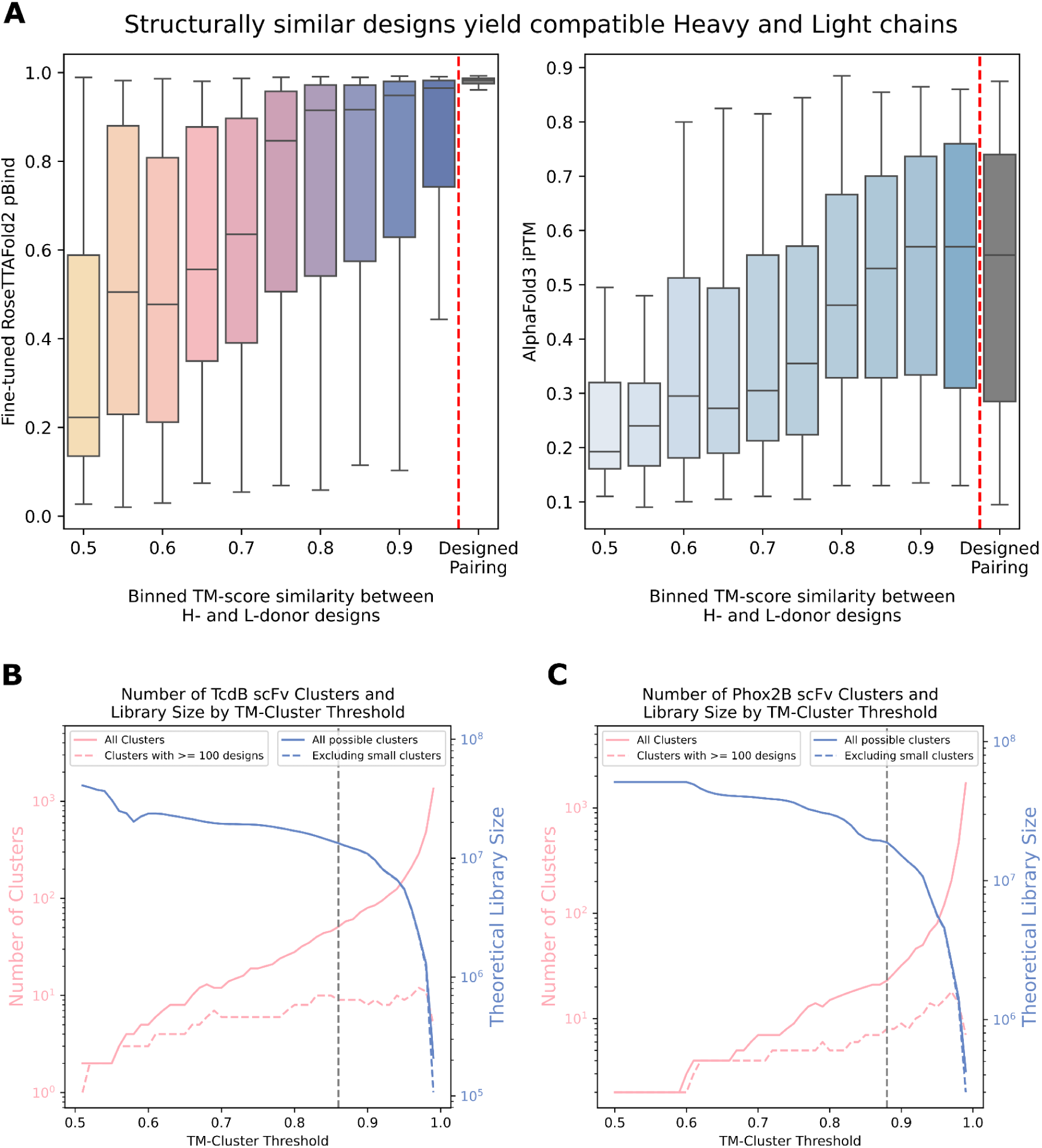
Computational validation of the structure-based combinatorial assembly strategy. Structure-based design permits the rational combinatorial assembly of heavy and light chains, assembling only heavy and light chains from structurally similar pairs. **A**) Fine-tuned RoseTTAFold (left), and AlphaFold3 (right) validate that pairing heavy and light chains from structurally similar (i.e. high pairwise TM score) designs yields scFvs that are more likely to be predicted to bind with high confidence (RF2 pBind, left; AF3 iPTM, right) than heavy and light chains from structurally-dissimilar (low pairwise TM score) designs. Note that the extremely high pBind distribution of the “designed pairings” (rightmost bar of left plot) is an artifact of those designs being specifically filtered for high pBind scores. **B-C**) combinatorial assembly leads to dramatically larger library sizes. Plots show the number of clusters (pink) at different TM score thresholds for TcdB (left) and Phox2b (right) scFvs. For the amplification strategy to work, each “cluster” becomes a PCR subpool, requiring independent PCR reactions (3 per subpool). Hence, we limit ourselves to large subpools (>= 100 designs), which maximises the combinatorial amplification for the amount of additional library assembly work. We additionally plot the theoretical library size for each target (blue), calculated as *number_of_clusters x cluster_size*^2^. Gray lines indicate the TM threshold chosen for library assembly, where library sizes approximately match the transfection efficiency of yeast (10^7^) ^58^.

**Extended Data Figure 23:**
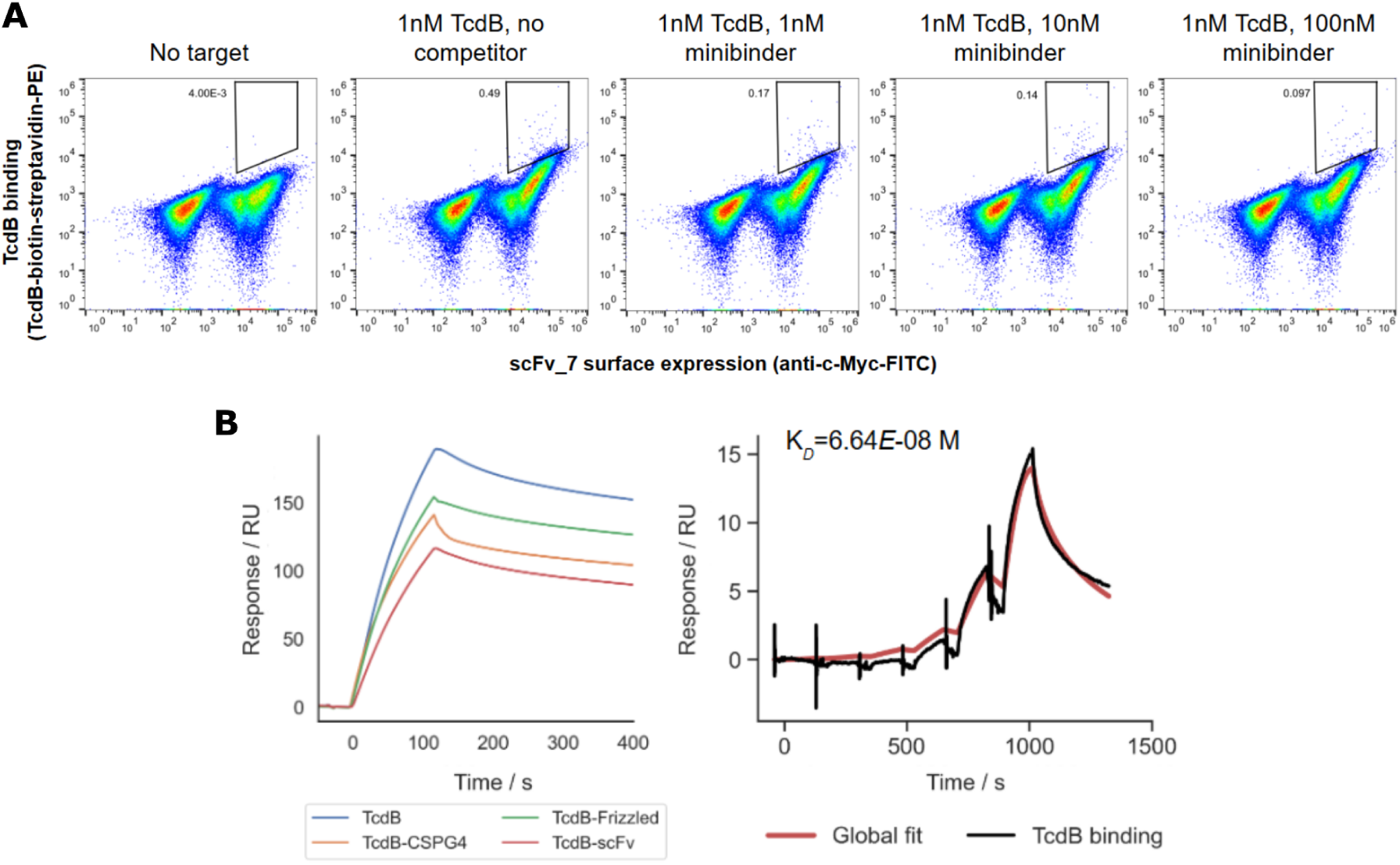
Negative data for TcdB 400 bp library. **A)** Yeast competition data for *scFv7* designed against the CSPG4-binding site of TcdB, in HL orientation. The scFv was identified by next-generation sequencing of prior FACS enrichments against TcdB and clonally expressed on EBY100 yeast for competition with a previously-characterized designed minibinder against the intended epitope^30^. Target binding was detected at nanomolar concentrations, but no significant reduction in binding signal was detected over the course of increasing competitor titrations. **B**) SPR analysis confirmed binding of the target while competition was not observed, suggesting binding at an incorrect site.

**Extended Data Figure 24:**
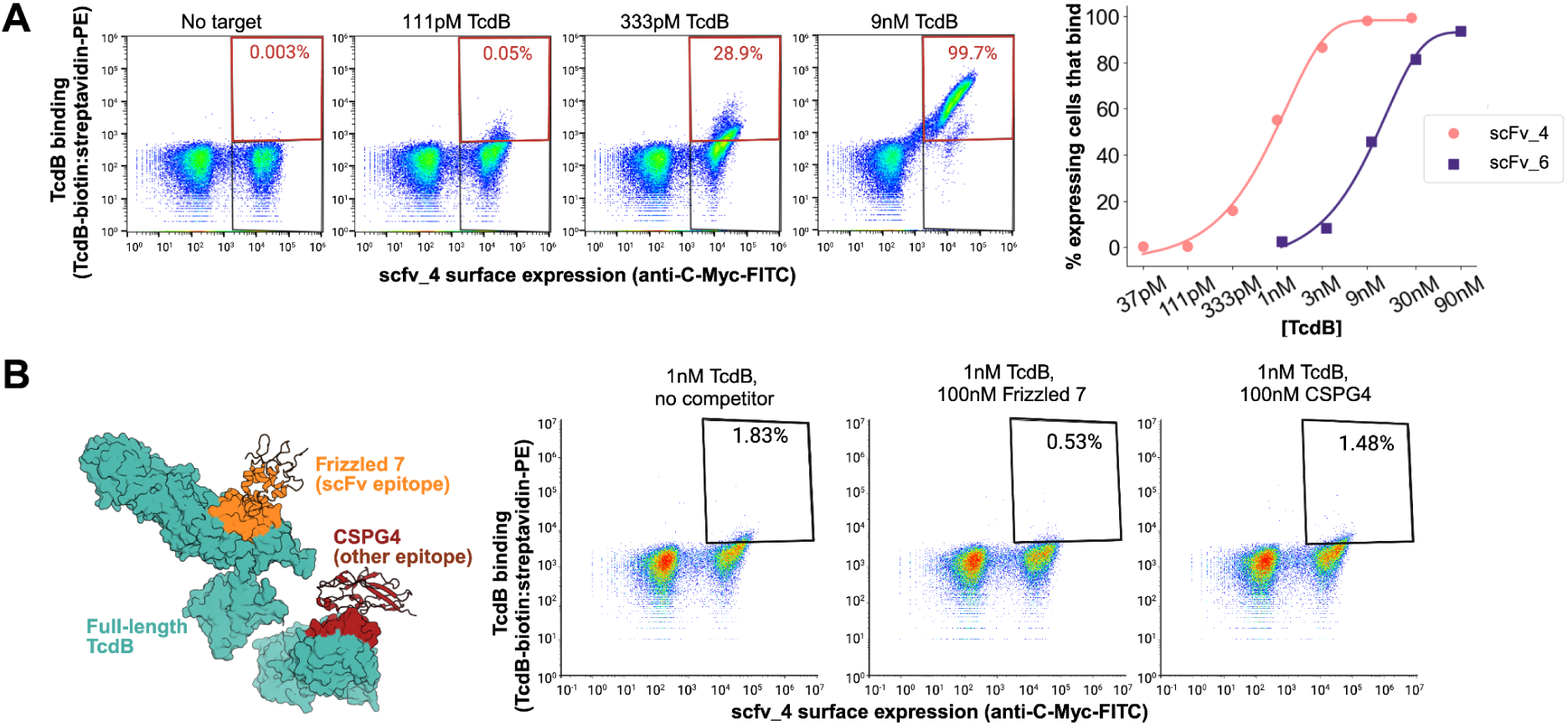
Characterization of scFvs that bind TcdB using yeast surface display flow cytometry. **A)** (left) Results of flow cytometry of yeast samples displaying *scFv4*-C-Myc construct. Each sample was treated with a titration of soluble biotinylated TcdB fragment (1401-1616) (bn-TcdB) concentrations and visualized with anti-C-Myc FITC + SAPE. (right) Percentage of expressing cells which are within the gate increase with bn-TcdB concentration. **B**) (left) scFvs were designed to bind to the Frizzled-7 epitope and therefore should compete with Frizzled-7. Designed scFvs should not compete with CSPG4, which binds at a different epitope on full-length TcdB. (right) Yeast displaying *scFv4*-C-Myc were incubated with 1 nM TcdB and either no competitor, 100 nM Frizzled-7, or 100 nM CSPG4. Binding signal decreases when Frizzled-7 is added, supporting that *scFv4* binds at the Frizzled-7 epitope. Binding signal does not significantly decrease when CSPG4 is added.

**Extended Data Figure 25:**
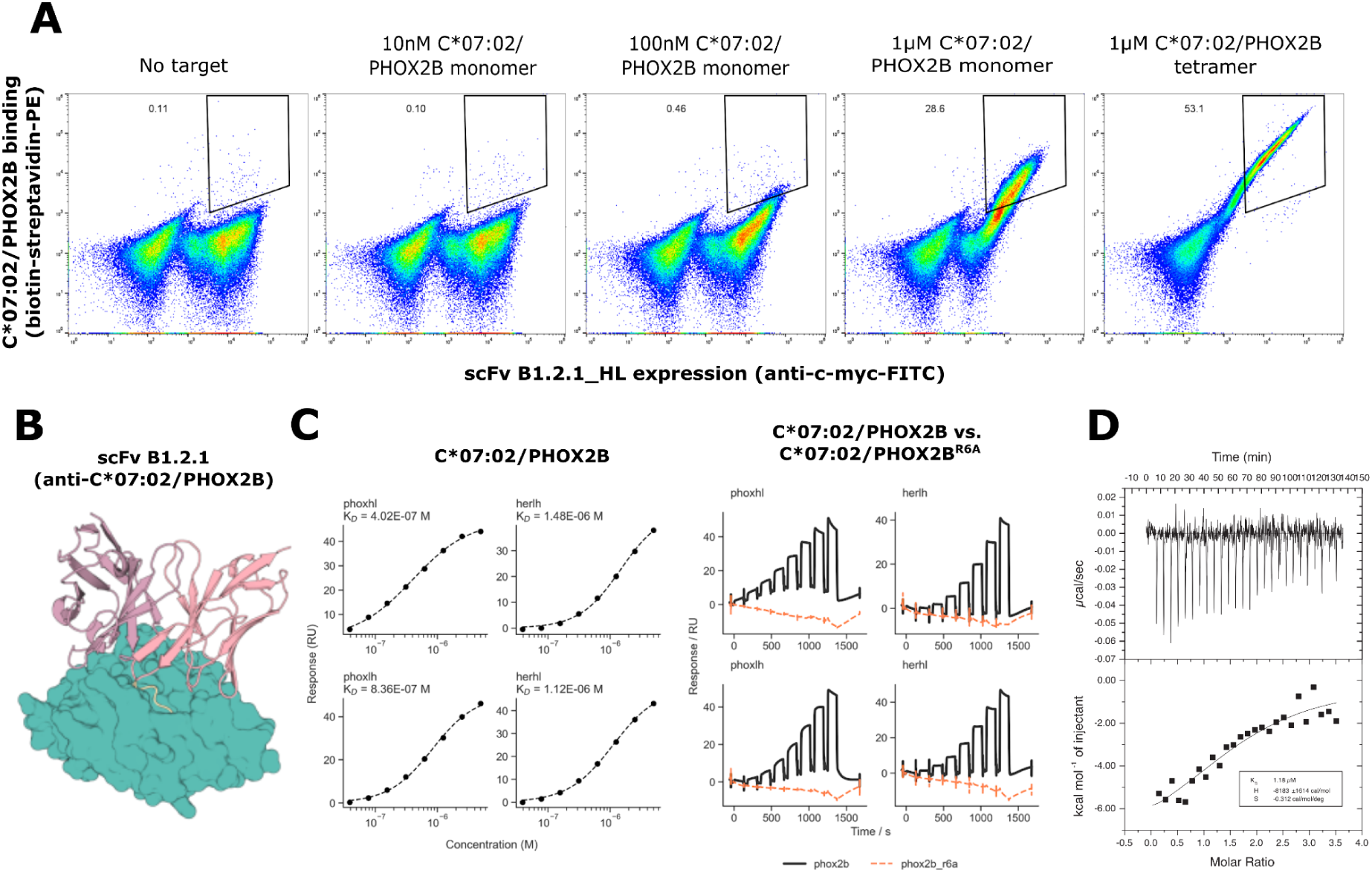
Biochemical characterization of a designed scFv specifically binding a therapeutically-relevant peptide-MHC. **A)** C*07:02/PHOX2B titration results with yeast surface display of anti-C*07:02/PHOX2B scFv B1.2.1, tested in the “HL” orientation with a (G4S)3 linker. For the tetramer condition, the biotinylated C:07:02/PHOX2B pHLA was tetramerized on streptavidin-PE (SAPE) prior to validation. The negative control is yeast incubated in the same concentrations of SAPE and FITC used in the experimental conditions in the absence of target. **B**) AF3 prediction of construct B1.2.1 docked to C*07:02/PHOX2B. **C**) Left: Surface plasmon resonance (SPR) data characterizing binding of B1.2.1 in the HL and LH orientations in the 10LH-based framework (“phox” prefix, left column) and the trastuzumab framework (“her” prefix, right column). B1.2.1 binds with approximately 1 μM affinity. Right: SPR data characterizing on-target binding of C*07:02/PHOX2B (“phox2b”) versus the same HLA bound to the R6A mutant of PHOX2B (“phox2b_r6a”). The results indicate specific binding to the intended target. **D**) Representative ITC titration of HLA-C*07:02/PHOX2B (30 μM) into a sample containing 2 μM herceptin_VLVH-His-Avi binder. Both samples contain 1 mM excess of PHOX2B peptide, to prevent the formation of empty HLA. The black line is the fit of the isotherm. Fitted values for K_D_, ΔH, and ΔS were determined using a 1-site binding model.

**Extended Data Figure 26:**
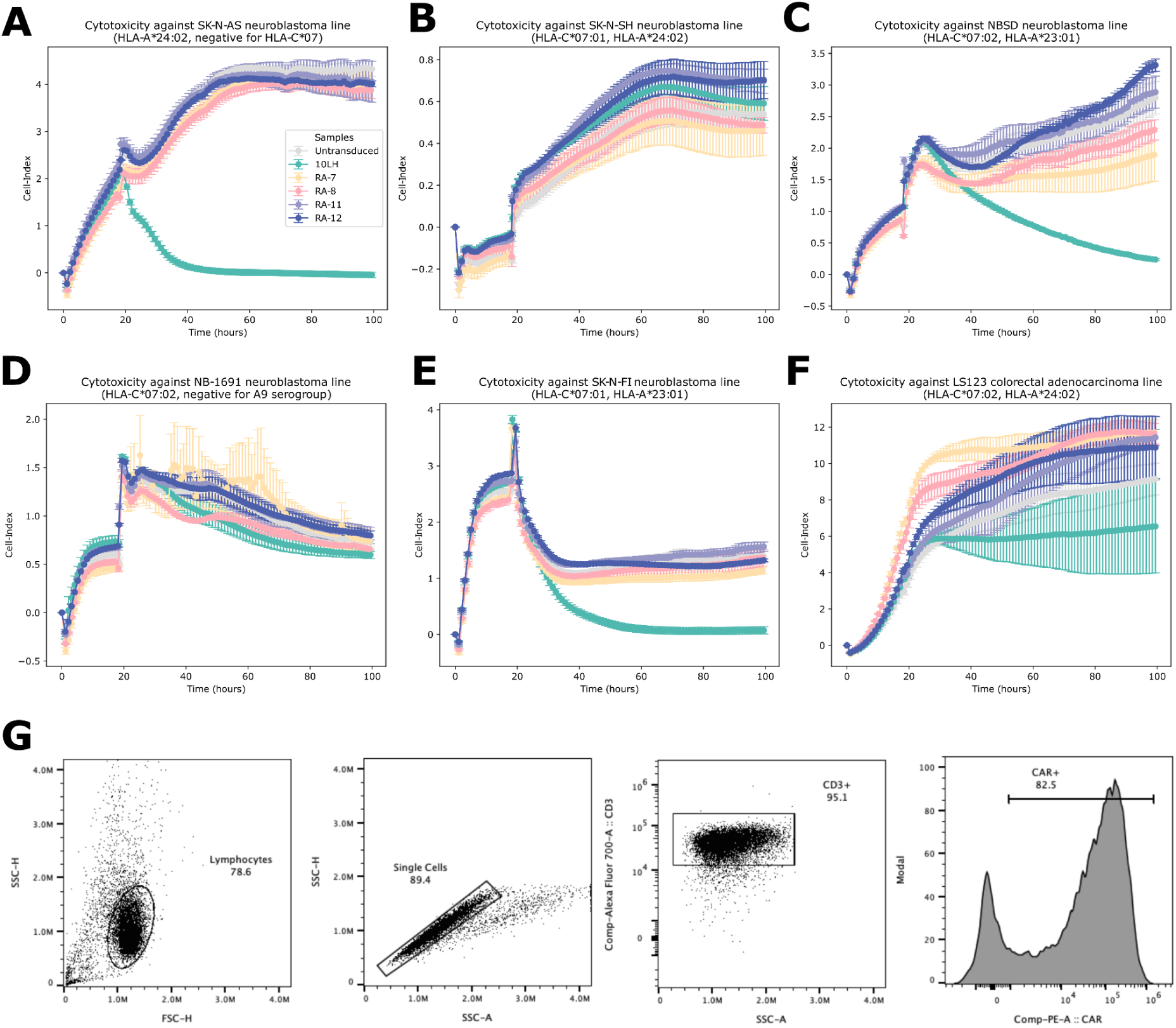
Low affinity designed Phox2B:HLA-C*07:02 scFvs do not show cytotoxicity. Cell index (CI) was monitored over time for neuroblastoma cell lines ((**A**) SK-N-AS, (**B**) SK-N-SH, (**C**) NBSD, (**D**) NB-1691, (**E**) SK-N-FI) and a colorectal adenocarcinoma cell line ((**F**) LS123) following the addition of CAR-T cells at total T cell effector-to-target ratio 15:1. CI values, representing cell viability and growth, were measured by the xCELLigence RTCA DP system at hourly intervals and normalized to baseline values to assess cell-mediated cytotoxicity. Mean and SEM are shown with each sample performed in triplicate, and rare outlier CI values were filtered. 10LH (positive control) demonstrated cytotoxicity against most Phox2b+ neuroblastoma lines expressing HLA-A*24:02 or HLA-A*23:01 and no cytotoxicity against Phox2b-LS123, as expected. In contrast, designed scFvs show no cytotoxicity against Phox2b+ neuroblastoma lines expressing HLA-C*07:02 or HLA-C*07:01. RA-7 and RA-8, as well as RA-11 and RA-12 comprise the same scFv with alternate heavy and light chain orderings. **G**) Transduced CAR T cells were stained with anti-(G_4_S)_3_ linker antibody, anti-CD3, anti-CD4 and anti-CD8 to confirm CAR expression. Staining for one example of positive control 10LH is shown.

**Extended Data Figure 27:**
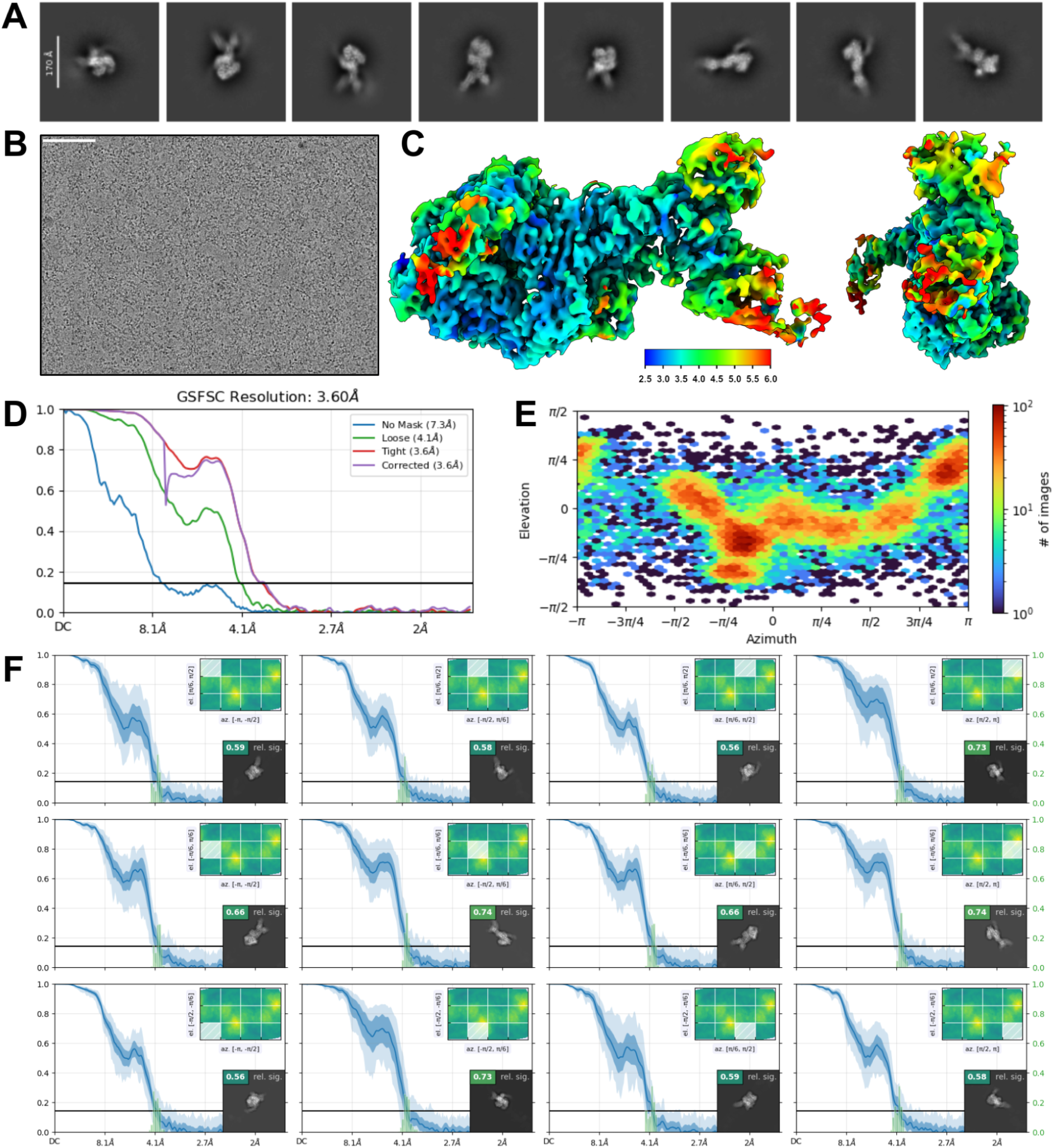
CryoEM statistics for TcdB in complex with scFv6. 10,897 movies were collected on a Glacios with a K3 detector. Thin ice was targeted for imaging, and only extended TcdB were observed. Heterogeneous refinement and apo structure were used to sort scFv bound TcdB (41,837 particles) and unbound apo TcdB (14,384 particles). **A**) 2D Class Averages. **B**) Representative micrograph **C**) Local Resolution map (Å), calculated using an FSC value of 0.143 viewed along two different angles **D**) Global Fourier Shell Correlation plot, Non Uniform Refinement **E**) Orientational distribution plots. **D**) Orientational diagnostics data.

**Extended Data Figure 28:**
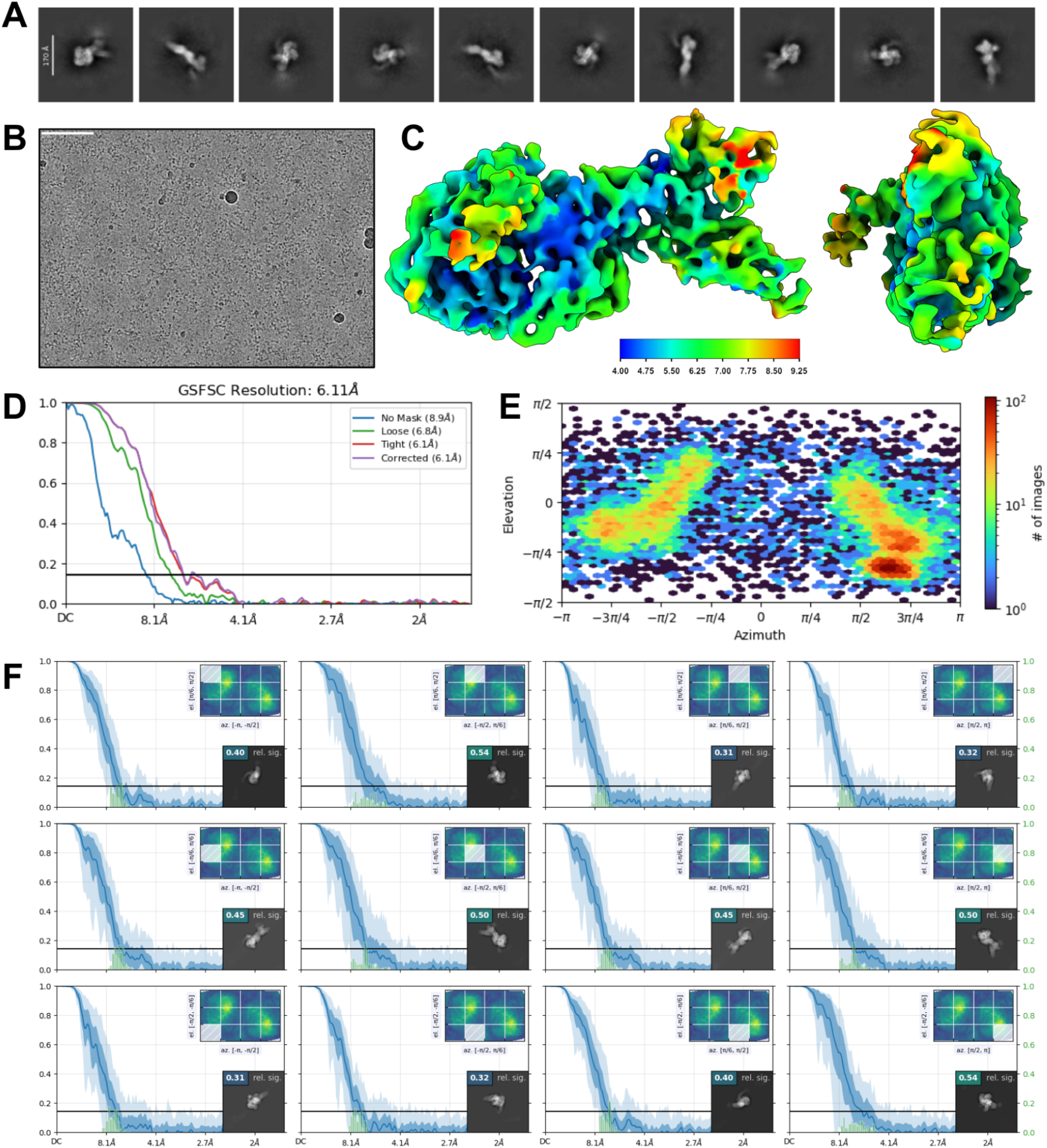
CryoEM statistics for TcdB in complex with scFv5. 8,304 movies were collected on a Glacios with a K3 detector. Thin ice was targeted for imaging, and only extended TcdB were observed. Heterogeneous refinement and apo structure were used to sort scFv-bound TcdB (12,540 particles) and unbound apo TcdB (3,667 particles). **A**) 2D Class Averages. **B**) Representative micrograph **C**) Local Resolution map (Å), calculated using an FSC value of 0.143 viewed along two different angles **D**) Global Fourier Shell Correlation plot, Non Uniform Refinement **E**) Orientational distribution plots. **D**) Orientational diagnostics data.

**Extended Data Figure 29:**
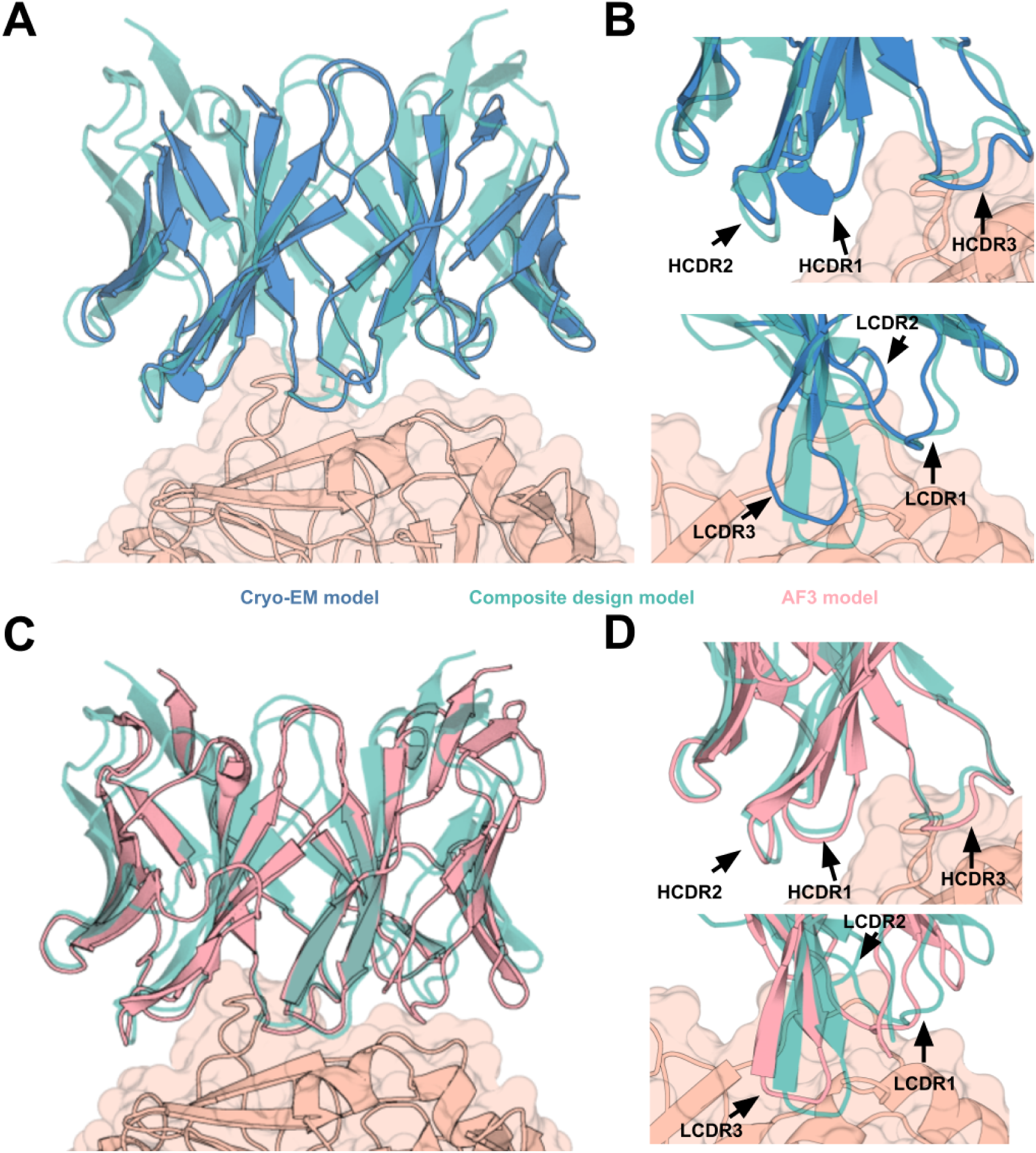
Cryo-EM structure and AF3 prediction of *scFv6* overlaid with the parent design models. *scFv6* is an assembly of heavy and light chains from two different (but near-superimposable) parent design models. **A-B**) Cryo-EM structure of *scFv6* aligned to the “composite design model” (the heavy and light chains from the respective parent designs). The structures were superimposed based on the target protein structure. This highlights that LCDRs and HCDRs from structurally clustered but independent designs retain their intended structure after combinatorial assembly. **C-D**) AlphaFold3 recapitulates the composite design model structure.

