## Supplementary Material for "Atomically accurate de novo design of antibodies with RFdiffusion"

February 2025

### Contents

I Methods for the Training of RFdiffusion and RoseTTAFold2 7

1 Training Datasets 7
1.1 SAbDab database of antibody structures 7
1.2 TCR/MHC-Peptide set of crystal and AlphaFold2-predicted structures 8
1.3 Loop-mediated interactions in the Protein Data Bank 8

2 Review of Vanilla RFdiffusion 9
2.1 Noising and Denoising 10
2.2 Losses 10
2.3 Structure Self-Conditioning 12
2.4 Training Procedure 12
2.5 Review of vanilla RFdiffusion fine-tuned on complexes 12
2.6 Dataset 13
2.7 Target Templating 13
2.8 Hotspots 13

3 RFdiffusion for Antibody Design 14
3.1 Datasets 14
3.2 Template Provision 14
3.3 Losses 16
3.4 Sequence Self Conditioning 16
3.5 Interface Hotspots 17
3.6 Antibody RFdiffusion Training Details 19

4 Review of RoseTTAFold2 19
4.1 RF2 Auxiliary Heads 20
4.2 Predicted LDDT (pLDDT) 20
4.3 Predicted Aligned Error (pAE) 20
4.4 Predicted Bind (pBind) 20

5 Fine-Tuning RoseTTAFold2 for Antibody Complex Prediction/Design Filtering 21
5.1 Updated pBind Prediction Head 21
5.2 Provision of “Hotspot” Information 21
5.3 Provision of Target Structure 23
5.4 Cropping of Input Chains 23
5.5 Cropping of SAbDab Set 23
5.6 Cropping of T-Cell Receptor (TCR) dataset 24
5.7 Cropping of Loop-Mediated Interaction Dataset 24

### Part I

### Methods for the Training of RFdiffusion and RoseTTAFold2

#### 1 Training Datasets

Three training sets were used to train RoseTTAFold2 and RFdiffusion, all representing primarily loop-based interactions with "target" proteins.

##### 1.1 SAbDab database of antibody structures

X-ray crystallographic and cryo-electron microscopy (cryo-EM) protein structures from the Structural Antibody Database (SAbDab) [1] deposited prior to January 13th, 2023 were used, with a resolution cutoff of 5Å. Sequences of the complementarity-determining regions (CDRs), as defined by the Chothia numbering system [2], were concatenated and clustered by 70% sequence similarity using MMseqs2 [3] to generate training clusters, yielding a median of 2 structures per cluster (95th percentile = 15). A 90:10 training:test split was sampled randomly from these clusters and kept consistent for all training runs (for both RoseTTAFold2 and RFdiffusion). This yielded 2235 training and 245 test clusters from 6205 total protein structures. Of these, 4936 had antigen present in the structure. Note that in many cases, multiple copies of an antibody are present in the protein structure. Each copy was included in the training set, and randomly sampled during training.

A final set of structures was used for validation, generated by combining two sets of structures. The first set were antibody-antigen complex structures released after 13th January, 2023, with resolution $< 5$Å and target sequence similarity $< 30\%$ to any target in the training set. The second set came from the original test set used during training, with additional verification that the target shared < 30% sequence similarity to any target in the training set. From each of these sets, a single example from each "cluster" was sampled, to prevent overrepresentation of particular complexes. To prevent an overrepresentation of antibody-peptide binders given this clustering strategy, we only included complexes where the target chain were between 100 and 500 residues These final sets totalled 104 and 47 structures respectively.

#### 1.2 TCR/MHC-Peptide set of crystal and AlphaFold2-predicted structures

To construct the TCR structure distillation dataset, experimentally validated TCR:pMHC binding interactions were downloaded from the VDJdb database [4] and modeled using the TCRdock software [5]. The VDJdb dataset was filtered to Class I-restricted, human and mouse complexes with paired TCR chains and fully resolved V, J, and CDR3 gene and sequence information. TCR:pMHC pairings from the large dextramer dataset from 10x Genomics were excluded due to potential non-specific dextramer binding. Epitopes with fewer than 10 paired TCRs were excluded. For epitopes with more than 50 paired TCRs, the associated TCRs were downsampled to 50 representatives using the distance-based approach described in the TCRdock manuscript [5].

The final dataset included 129 crystal structures (from the PDB) and 2574 AF2-predicted structures, covering 39 unique MHC subtypes. A 90:10 training:test split was randomly sampled based on MHC subtype, yielding 35 training clusters and 4 test clusters. Note that only crystal structures were used for the test set (AF2 models for these MHC subtypes were excluded from training).

#### 1.3 Loop-mediated interactions in the Protein Data Bank

The Protein Data Bank (PDB) [6] was mined for hetero-dimeric complexes where at least one side of the interface was primarily loop-based. Pyrosetta [7] was used to determine the secondary structure of all chains in the PDB (date cutoff August 2nd, 2021). Hetero-dimeric complexes (excluding structures present in SAbDab) were then identified and X-ray and cryo-EM structures with resolution $< 5$Å were selected. Interface residues were then defined as residues being $< 10$Å $C\alpha - C\alpha$ distance between the two chains. The proportion of these residues within loops was calculated, and complexes where $> 70\%$ of interacting residues were within loops, and these loops summed to $> 10$ residues in length (to filter for structures where interacting loops were comparable in length to antibodies) were selected. The protein in the hetero-dimeric complex with the loop-based interface was designated the "binder". To more closely mimic the antibody dataset, we also filtered for hetero-dimeric complexes where both chains were reasonably structured ($> 40\%$ non-loop secondary structure). This was decided because many loop-based interactions in the PDB are generally unstructured (with structure dependent on non-protein molecules such as DNA or RNA). Finally, in line with the fact that RFdiffusion, when trained on hetero-dimeric complexes, trains on complete protein chains as the "binder" side (rather than cropping them), hetero-dimeric complexes where the "binder" was $\leq 250$ residues in length were used for training (such that the "binder" was not cropped during training). Clusters were determined in line with the clusters used to train RoseTTAFold2 [8] (30% sequence similarity), and a 90:10 train:test set was randomly sampled, and kept constant in all training runs. This yielded 2010 training and 254 test clusters, with 18767 total hetero-dimeric complexes.

### 2 Review of Vanilla RFdiffusion

The antibody design version of RFdiffusion is fine-tuned from the vanilla RFdiffusion [9] base model and shares many similarities. Here, we briefly review the fundamentals of vanilla RFdiffusion, as they relate to the antibody variant described here. A thorough description of the training and architecture of vanilla RFdiffusion can be found elsewhere [9].

#### 2.1 Noising and Denoising

RFdiffusion is a Denoising Diffusion Probabilistic Model (DDPM), a class of generative model first described in Sohl-Dickstein et al. [10] and expanded later [11]. DDPMs formulate sampling from an unknown data distribution as an iterative denoising task; an initial sample $x^{(T)}$ is drawn from a known distribution (typically a Gaussian distribution), this sample is then progressively denoised by the DDPM until a completely denoised sample $\hat{x}^{(0)}$ is obtained, that closely matches the data distribution. DDPMs are trained by taking samples $x^{(0)}$ from the data distribution (in our case protein structures) and applying noise to these samples according to a defined noising schedule. DDPMs are trained to remove the added noise and produce the original sample.

Following the work of AlphaFold2 [12], RFdiffusion represents protein backbones as a collection of residue frames where each frame has both a translation and a rotation. RFdiffusion formulates the noising process as a joint process in both translation and rotation space. Translations are noised by the addition of three-dimensional Gaussian noise and rotations are noised with Brownian motion on SO3.

We use the same procedure to sample from the reference distribution and to interpolate between $\hat{x}^{(t)}$ and $x^{(t-1)}$ as vanilla RFdiffusion. As such, we provide these algorithms (which originally appeared in the vanilla RFdiffusion Supplement) as Algorithm 1 for reference.

#### 2.2 Losses

In contrast to structure prediction models such as AF2 and RF2 which use the Frame Aligned Point Error (FAPE), first described in the AF2 paper, RFdiffusion uses a Mean Squared Error (MSE) loss over predicted residue frames as the primary training objective. A crucial difference between FAPE and MSE is that FAPE is invariant to global translations and rotations, whilst MSE is not (MSE is also an L2 vs L1 loss, which drives RFdiffusion to match the data distribution). It was found empirically that the MSE loss yielded a model which performed substantially better

**Algorithm 1** Vanilla RFdiffusion Reference Functions

**function** SampleReference(L)
    ▷ Random initial structure for $L$ residues
    **for** $l = 1, \ldots, L$ **do**
        $r_l^{(T)} \sim \text{Uniform}(SO(3))$
        $z_l^{(T)} \sim \mathcal{N}(0, I_3)$
        $x_l^{(T)} = (r_l^{(T)}, x_l^{(T)})$
    **end for**
**return** $x^{(T)}$
**end function**

**function** ReverseStep($x^{(t)}, \hat{x}^{(0)}$)
    ▷ One step of reverse diffusion
    **for** $l = 1, \ldots, L$ **do**
        $(r_l^{(t)}, z_l^{(t)}) = x_l^{(t)}$
        $(\hat{r}_l^{(0)}, \hat{z}_l^{(0)}) = \hat{x}_l^{(0)}$
        ▷ Update translations
        $z_l^{(t-1)} \sim \mathcal{N}(\frac{\sqrt{\bar{\alpha}^{(t-1)}}\beta^{(t)}}{1-\bar{\alpha}^{(t)}}\hat{z}_l^{(0)} + \frac{\sqrt{\alpha^{(t)}}(1-\bar{\alpha}^{(t-1)})}{1-\bar{\alpha}^{(t)}} z_l^{(t)}, \beta^{(t)} I_3)$

        ▷ Update rotations
        $s_l =$ RotationScoreApproximation($r_l^{(t)}, \hat{r}_l^{(0)}, \sigma_t^2$)
        $\epsilon_{l,1}, \epsilon_{l,2}, \epsilon_{l,3} \overset{iid}{\sim} \mathcal{N}(0, 1)$
        $r_l^{(t-1)} = r_l^{(t)} \exp_{I_3}\left\{(\sigma_t^2 - \sigma_{t-1}^2) {r_l^{(t)}}^\top s_l + \sqrt{\sigma_t^2 - \sigma_{t-1}^2} \sum_{d=1}^3 \epsilon_{l,d} f_d\right\}$
        $x_l^{(t-1)} = (r_l^{(t-1)}, z_l^{(t-1)})$
    **end for**
**return** $x^{(t-1)}$
**end function**

than the same model trained with FAPE loss. A separate MSE loss over both the translations and rotations of each frame is applied.

#### 2.3 Structure Self-Conditioning

Traditionally, DDPMs are parametrized to take as input only the noised example from the previous time step. RFdiffusion is trained to also take as input the previous timestep's prediction of the ground-truth example; this is a technique called self-conditioning that has been shown in other work to dramatically improve the quality of DDPM outputs [13]. Self-conditioning also dramatically improves the quality of vanilla RFdiffusion outputs, and has subsequently been used to improve performance of a number of other generative design models [14, 15, 16].

#### 2.4 Training Procedure

Vanilla RFdiffusion is trained on the set of monomer structures in the PDB which were used for RoseTTAFold2 training. Only structures with a total number of residues $\leq 384$ were used in training. At each step of training of vanilla RFdiffusion, a sample $x^{(0)}$ is selected from the dataset and $t$ steps of noise are added to obtain a corrupted input, $x^{(t)}$. Vanilla RFdiffusion is trained unconditionally (no ground truth structure and sequence given) 20% of the time and 80% of the time the model is trained with an unnoised motif (region of known structure and sequence) provided to the model. RFdiffusion is tasked with predicting the ground truth structure of the noised region. The base vanilla RFdiffusion model is trained for 5 epochs, where each epoch contains 25,600 structures.

#### 2.5 Review of vanilla RFdiffusion fine-tuned on complexes

The complex fine-tuned version of vanilla RFdiffusion is described in detail elsewhere [9]. Here, we briefly review the aspects of the training procedure which are relevant to the antibody variant of

RFdiffusion described in this work.

#### 2.6 Dataset

The complex fine-tuned model of vanilla RFdiffusion was trained starting from a version of the aforementioned vanilla RFdiffusion (trained only on monomer structures). The model was then trained on a mixture of 50% monomer examples and 50% complex examples (the monomer and complex examples are from the PDB datasets used in the training of RoseTTAFold2 [8]).

#### 2.7 Target Templating

In keeping with the precedent of other binder design pipelines [17, 18], vanilla RFdiffusion assumes that the target backbone structure does not change upon binding and can be held fixed during design. To accomplish this, when the network is given a complex example, one side of the complex is noised and the other side has both its backbone, sidechains, and sequence provided to the model. The model learns to only denoise the binder chain and to keep the target chain fixed.

#### 2.8 Hotspots

The complex fine-tuned version of vanilla RFdiffusion is trained to take information about which surface site on a target protein to interact with. This allows the model to be controllable by the user at inference time. Hotspot residues are defined as any residue on the unnoised (target) chain which is within 8Å $C_\beta - C_\beta$ distance of the noised (binder) chain. During training time, $0 - 20\%$ of hotspots are provided to the model as a one-hot vector appended to the t1d feature.

### 3 RFdiffusion for Antibody Design

#### 3.1 Datasets

For antibody design, RFdiffusion is trained on the three datasets described above (SabDab, TCR structures, and loops at interfaces from the PDB) in a ratio of 50% SabDab, 10% TCR, 40% loops at interfaces. For antibody structures without a resolved target structure, the whole structure (i.e. the whole antibody) is noised. In all other cases, the target chain backbone and sequence (but not sidechains; this contrasts to vanilla RFdiffusion) is provided to RFdiffusion (for TCRs, the target chain is the peptide-MHC, and for the PDB examples the target structure is the chain with the non-loop-based interface). The antibody/TCR/loop-based interaction chain is then noised. RFdiffusion is tasked with predicting the ground truth structure of the entire system.

#### 3.2 Template Provision

RFdiffusion is trained to keep the non-CDR region of the antibody/TCR fixed. This is done by providing the model with the structure of the non-CDR framework region through the t2d (template) input. These template entries in t2d are marked with a one-hot vector in the 44th dimension of t2d. In the case of the non-antibody loop-based interfaces dataset, the loop-interfaces are treated as above (like CDRs), and the rest of the protein is treated as framework.

The structure self-conditioning track of the model also runs through the t2d track so when the model is processing self-conditioning information, the templated framework region of the binder is corrected to be the input template region and marked in the 44th dimension as being templated. The other entries (from the self-conditioning input) are marked in the 45th dimension as being from self-conditioning, this is described in Algorithm 2.

**Algorithm 2** t2d Self Conditioning

**function** T2D_NoStrSelfCond($template\_dgram_{ij}, framework\_mask_{ij}$)
▷ Generate Initial xyz_t and t2d features for self conditioning track

▷ $template\_dgram_{ij} \in \mathbb{R}^{44}$
▷ $framework\_mask_{i,j} \in \{0, 1\}$

▷ $xyz\_t$ always initialized to zeros
$xyz\_t = \vec{0}$ ▷ $xyz\_t \in \mathbb{R}^{L \cdot 27 \cdot 3}$

$t2d_{ij} = \vec{0}$ ▷ $t2d_{ij} \in \mathbb{R}^{45}$

$t2d_{ij1:43} = template\_dgram_{ij1:43} * framework\_mask_{i,j}$
$t2d_{ij44} = framework\_mask_{i,j}$ ▷ Indicate which contacts are from the provided framework
**return** $xyz\_t, t2d$
**end function**

**function** T2D_WithStrSelfCond($t2d\_prev_{ij}, template\_dgram_{ij}, framework\_mask_{ij}$)
▷ Correct t2d framework entries. t2d_prev is the distogram representation of $\hat{x}^{(0)}_{\text{prev.}}$
▷ $t2d\_prev_{ij} \in \mathbb{R}^{44}$
▷ $template\_dgram_{ij} \in \mathbb{R}^{44}$
▷ $framework\_mask_{i,j} \in \{0, 1\}$

$t2d_{ij} = \vec{0}$ ▷ $t2d_{ij} \in \mathbb{R}^{45}$

$t2d_{ij1:43} = template\_dgram_{ij1:43} * framework\_mask_{i,j}$

$t2d_{ij44} = framework\_mask_{i,j}$ ▷ Indicate which contacts are from the provided framework
$t2d_{ij45} = not\ framework\_mask_{i,j}$ ▷ Indicate which contacts are from the self conditioning input
**return** $t2d$
**end function**

| Loss | Description | Weight |
|---|---|---|
| $\mathcal{L}_{\text{Frame}}$ | MSE loss on frames | 1.0 |
| $\mathcal{L}_{\text{2D}}$ | Loss on distogram prediction | 1.0 |
| $\mathcal{L}_{\text{MLM}}$ | Masked language modelling loss | 1.0 |

Supplementary Methods Table 1: Losses used to train RFdiffusion.

##### 3.3 Losses

In addition to the vanilla RFdiffusion losses described above, RFdiffusion is trained with a masked sequence prediction loss. This loss is implemented as a categorical cross entropy loss between the true sequence and the predicted sequence in the masked region. This loss is added to training to encourage the model to build an explicit understanding of sequence which is critical for the sequence self-conditioning technique we describe below.

Descriptions and weights of all of the losses applied during RFdiffusion training are provided in Supplementary Methods Table 1 (note that the losses besides $\mathcal{L}_{\text{MLM}}$ are identical to those used in vanilla RFdiffusion).

##### 3.4 Sequence Self Conditioning

RFdiffusion has a sequence self-conditioning track in addition to the structure self-conditioning track which was reviewed in Section 2.3. At inference time, the sequence self-conditioning track takes the previous timestep's final sequence embedding and provides this as input to the next denoising step. This allows the model to jointly reason over sequence and structure. RFdiffusion generation using the sequence self-conditioning track is shown in Algorithm 3.

The "base" version of vanilla RFdiffusion used in this study to fine-tune the antibody variant was trained with this sequence self-conditioning. It trained with the same hyperparameters as "vanilla" RFdiffusion for protein complexes [9] (and Section 2.5), except for the provision of sequence self-conditioning 50% of the time during training, and the application of the amino acid cross entropy

**Algorithm 3** RFdiffusion Generation

**function** SAMPLE(L)
    ▷ RFdiffusion generation of $L$-residue backbone structure
    $x^{(T)} = \text{SampleReference}(L)$
    $\hat{x}^{(0)}_{\text{prev.}} = \vec{0}$ ▷ Initialize structure self-conditioning
    $\hat{s}^{(0)}_{\text{prev.}} = \vec{0}$ ▷ Initialize sequence self-conditioning; $\hat{s}^{(0)}_{\text{prev.}} \in \mathbb{R}^{L \cdot C_m}$
    **for** $t = T, \ldots, 1$ **do**
        $\hat{x}^{(0)}, \hat{s}^{(0)} = \text{RFDIFFUSION}(x^{(t)}, \hat{x}^{(0)}_{\text{prev.}}, \hat{s}^{(0)}_{\text{prev.}})$
        $x^{(t-1)} = \text{REVERSESTEP}(x^{(t)}, \hat{x}^{(0)})$
        $\hat{x}^{(0)}_{\text{prev.}} = \hat{x}^{(0)}$
        $\hat{s}^{(0)}_{\text{prev.}} = \hat{s}^{(0)}$
    **end for**
**return** $\hat{x}^{(0)}$
**end function**

loss ($\mathcal{L}_{\text{MLM}}$). This cross entropy loss is preferentially applied at timesteps close to $t = 0$ by weighting it with a sigmoid weighting, fit empirically such that the loss is mostly applied in the $t < 50$ period. This model trained for 5 epochs with 25600 examples per epoch.

#### 3.5 Interface Hotspots

To allow the targeting of specific epitopes, "interface hotspots" are provided in a manner similar to in vanilla RFdiffusion. The hotspot definition deviates slightly from that of vanilla RFdiffusion, with a target residue classified as a hotspot if the residue has an average $C_\beta$ distance to the closest 5 antibody CDR residues of less than 8Å. This definition was used so as to encourage CDR-mediated interactions with the epitope, rather than framework-mediated interactions (which was initially a problem, especially for VHH design).

For each example provided to the model at training time, a random fraction between $0 - 100\%$ of the hotspot residues are provided to the model as a one-hot vector appended to the t1d feature.

| Input Name (Shape) | Description | Same as in vanilla RFdiffusion? |
|---|---|---|
| msa_masked (1,1,L,48) | The masked sequence (20aa, zeros (1), mask (1), repeat aa (20), zeros (1), repeat mask (1), zeros (2), N-term/C-term (2)) | yes |
| msa_full (1,1,L,25) | The masked sequence (20aa, mask (1), zeros (1), N-term/C-term (2)) | yes |
| seq (1,L,22) | The masked sequence (20aa, zeros (1), mask (1)) | yes |
| xyz_prev (L,27,3) | The coordinates of all atoms in $x^t$ (N-Ca-C-O backbone (4), 10 sidechain atoms (zeros if no atom present), 13 hydrogen atoms (zeros if no atom present)) | yes |
| idx_pdb (L) | The integer index of each residue. Used to create a positional encoding. | yes |
| t1d (1,L,23) | The one-dimensional features associated with $x^t$ (20 amino acids, mask (1), timestep (1), hotspot boolean (1)). The timestep is set to 1 for all target residues, and to $1 - t/T$ in all other positions. | yes |
| t2d (1,L,L,45) | The two-dimensional, residue-pair features associated with the self-conditioning feature Section 2.3. The intra-chain contacts of the antibody framework regions and the target region are corrected as described in Section 3.2, whether each position is from a correction or self-conditioning is noted as a boolean in dimensions 44 and 45, respectively (36 distance bins (2-20Å, 0.5Åbins,) + 1 final distance bin (> 20Å), angle maps (sine and cosine of omega, theta, and phi angle) (6), corrected contact mask (1), self-conditioning contact mask (1)) | no |
| xyz_t (1,L,27,3) | The self-conditioning feature. Described in Section 2.3. This feature is immediately converted to a distogram and anglegram representation by the model; only the first 3 atom coordinates (corresponding to N, CA, and C backbone atoms) are used. | yes |
| alpha_t (1,L,30) | The sidechain torsions provided to the model. No sidechain information is provided to the model in RFdiffusion and this feature is always all zeros. | no |
| msa_prev (L,$C_m$) | The MSA embedding self-conditioning feature. As described in Section 3.4. $C_m = 256$ | no |
| pair_prev (L,L,$C_p$) | The 2D embedding recycling information. Recycling is not used in RFdiffusion-antibody and this feature is always all zeros. $C_p = 128$ | yes |
| state_prev (L,$C_s$) | The 1-D embedding recycling information. Recycling is not used in RFdiffusion-antibody and this feature is always all zeros. $C_s = 16$ | yes |

Supplementary Methods Table 2: Description of Features Input to RFdiffusion

##### 3.6 Antibody RFdiffusion Training Details

The RFdiffusion used in this manuscript was trained starting from the sequence self-conditioning version of vanilla RFdiffusion described in Section 3.4. We trained with a crop size of 768 and a pseudo-batch size of 16. RFdiffusion was trained for 70 epochs with 512 examples per epoch. At the start of training, the learning rate is linearly increased (warmed up) from 0 to 0.0005 over the first 100 gradient steps. All other training settings used were the same as in vanilla RFdiffusion.

### 4 Review of RoseTTAFold2

RoseTTAFold2 [8] (RF2) is an updated version of the original RoseTTAFold architecture [8]. The training of RF2 is described in detail in Baek et al., 2023 [8]. Here, we briefly review RF2 to give context for the fine-tuned version we use for antibody design filtering.

The RF2 architecture comprises three "tracks"; a 1D multiple sequence alignment (MSA) track, a 2D residue-pair track, and a 3D structure track, in which residue frames are iteratively refined into the final output 3D structure. Inputs to RF2 comprise the query sequence, processed MSAs and templates of homologous structures from the PDB. These inputs are embedded and used to initialise the three tracks. 36 "structure blocks" are then performed, where the three tracks are sequentially updated from their previous values and from projections from the other tracks. Four final blocks of the 3D track are carried out, with frozen 1D and 2D data from the 36th layer of RF2. Following AF2, RF2 "recycles" the output from the previous pass (through the 40 blocks) 0-3 times each training iteration.

RF2 trains on three training datasets. Firstly, single chains from the PDB (deposited prior to April 30th, 2020) with resolution $< 4.5$Å are clustered with an e-value cutoff of 1e-3 into approximately 20,000 clusters representing 280,000 chains. Secondly, a "distillation" set of monomers was created using AF2. 12 million UniRef50 [19] sequences were folded by AF2 [20]. High confidence (pLDDT $> 80$) structures of $> 200$ residues in length were used. Structures were clustered as before into 1.0 million clusters from 3.6 million total structures. Structures with sequence similarity $> 30\%$ to anything in the PDB-derived training set was excluded. The third set was dimeric structures in the PDB. These represent every pair of contacting chains in a biological assembly, and is split into homomeric and heteromeric interactions. These sets were sampled 25:50:25 during training.

#### 4.1 RF2 Auxiliary Heads

In addition to predicting the structure to which the provided amino acid sequence will fold, RF2 also predicts several other values. These other values are predicted from the main tracks of the model using small networks called "auxiliary heads". We review the auxiliary heads here.

#### 4.2 Predicted LDDT (pLDDT)

Following AF2, RF2 predicts the per-residue LDDT [21] of the predicted structure.

#### 4.3 Predicted Aligned Error (pAE)

Following AF2, RF2 predicts the aligned error for each pair of residues. This is predicted from the 2D residue-pair track embedding (124 dimensions) projected down to 64 pAE logits.

#### 4.4 Predicted Bind (pBind)

RF2 also predicts whether two chains will form a complex or not through the pBind head. The pBind head projects from the 64 pAE logits of the interchain contacts down to a single value. This value is converted to a probability of binding by applying a sigmoid function (Algorithm 4).

**Algorithm 4** RF2 pBind Auxiliary Head

**function** CALCULATE_PBIND($pAE_{ij}, sameChain_{ij}$)
  ▷ $pAE_{ij} \in \mathbb{R}^{64}$
  ▷ $sameChain_{ij} \in \{0, 1\}$

  $L_{nonzero} = \sum_{i=1}^{L} \sum_{l=1}^{L} sameChain_{ij}$

  $mean\_bins = \frac{1}{L_{nonzero}} * \sum_{i=1}^{L} \sum_{l=1}^{L} pAE_{ij} * sameChain_{ij}$

  $pBind = \text{Sigmoid}(\text{Linear}(mean\_bins))$ ▷ $pBind \in \mathbb{R}$
**return** $pBind$
**end function**

### 5 Fine-Tuning RoseTTAFold2 for Antibody Complex Prediction/Design Filtering

We fine-tuned from RF2 on the same datasets used to fine-tune RFdiffusion for antibody design, with modifications described below.

#### 5.1 Updated pBind Prediction Head

We reasoned that the model could more effectively learn to predict pAE and pBind if they were predicted by separate auxiliary heads. We developed an updated pBind prediction model which projects off of all three tracks of the model (Algorithm 5).

#### 5.2 Provision of "Hotspot" Information

Hotspots were defined as described above. A random proportion of these (0-100%) were provided to the network during training, as a one-dimensional one-hot feature describing which residues on the target are contacted by the antibody/TCR/"binder" chain.

**Algorithm 5** RF2 Updated pBind Auxiliary Head

**function** CALCULATE_PBIND($pair_{ij}, rbf_{ij}, state_i, sameChain_{ij}$)
    ▷ $pair_{ij} \in \mathbb{R}^{128}$
    ▷ $rbf_{ij} \in \mathbb{R}^{64}$
    ▷ $state_i \in \mathbb{R}^{32}$
    ▷ $sameChain_{ij} \in \{0, 1\}$

    $attn_{i*j} = \text{flatten}(\text{Linear}(rbf_{ij}))$ ▷ $attn_{i*j} \in \mathbb{R}$

    $[left_{i1}]_j = [state_{i1} \text{ for } j \in L]$
    $[right_{1i}]_j = [state_{1i} \text{ for } j \in L]$

    $left_{ij} = \text{concat}_j([left_{i1}]_j)$
    $right_{ji} = \text{concat}_j([right_{1i}]_j)$

    $feat_{ij} = \text{concat}(pair_{ij}, left_{ij}, right_{ji})$ ▷ $feat_{ij} \in \mathbb{R}^{128+32+32}$
    $logits_{i*j} = \text{flatten}(\text{Linear}(feat_{ij}))$ ▷ $logits_{i*j} \in \mathbb{R}$

    ▷ Mask out residue pairs which are on the same chain
    $flatSameChain_{i*j} = \text{flatten}(sameChain_{ij})$
    $masked\_attn_l = attn_{i*j}[flatSameChain_{i*j} == 0]$
    $masked\_logits_l = logits_{i*j}[flatSameChain_{i*j} == 0]$

    $masked\_logits_l = \text{Softmax}(masked\_logits_l)$
    $logits_{inter} = \text{mean}_l(masked\_attn_l * masked\_logits_l)$ ▷ $logits_{inter} \in \mathbb{R}$

    $pBind = \text{Sigmoid}(logits_{inter})$
**return** $pBind$
**end function**

#### 5.3 Provision of Target Structure

During training, the cropped (Section 5.4) target structure is provided to the network, through both the two-dimensional and three-dimensional track of RF2. Sidechain positions, however, were not provided.

#### 5.4 Cropping of Input Chains

Due to GPU memory constraints, cropping of the input sequence/structure was required. This was performed slightly differently for each of the sets used during training.

#### 5.5 Cropping of SAbDab Set

For antibody structures, both the antibody and target chains were cropped (when necessary). The heavy chain (IgH) was cropped from the first residue up to residue 105-115 (inclusive, randomly sampled during training) based on the Chothia numbering format. This approximately corresponds to the variable domain ($V_H$). This cropping was also used for nanobody structures. The heavy chain (IgL) was similarly cropped, from the first residue up to residue 100-110 (inclusive, randomly sampled during training), again based on the Chothia numbering format. This approximately corresponds to the variable domain ($V_L$).

The antigen/target chain was then cropped spatially to $MaxLength - (len(IgH) + len(IgL))$. The maximum length was typically set to 384. The centroid residue of the spatial crop was randomly sampled from the calculated "hotspot" residues on the target chain. The closest N residues of the target chain were included, and provided to the network (see above).

#### 5.6 Cropping of T-Cell Receptor (TCR) dataset

During preparation of the TCR/pMHC dataset, the TCR is cropped to the $V_\alpha$ and $V_\beta$ domains and the MHC is cropped to the $\alpha_1$ and $\alpha_2$ subunits (for MHC class I) or $\alpha_1$ and $\beta_1$ subunits (for MHC class II). The peptide is not cropped. The maximum total length of this cropped TCR/pMHC complex is 436 residues and no further cropping is performed at training time.

#### 5.7 Cropping of Loop-Mediated Interaction Dataset

As aforementioned, the loop-mediated interaction dataset was curated such that the "binder" chain was $< 250$ residues in length. This chain was therefore not cropped. The "target" chain was cropped identically to how antigen/target chains are cropped in the SAbDab case (spatially around a randomly-selected "hotspot" residue).

#### 5.8 Provision of "Framework" Structure for Loop Dataset

For antibody and TCR cases, we reasoned that the geometry of the two Ig domains ($V_H/V_L$; $V_\alpha/V_\beta$) is sufficiently constrained to be accurately predicted from single amino acid sequence. However, for the set of loop-mediated hetero-dimeric complexes from the PDB, we predicted that the structure of the "framework" (non-loop, non-interacting) part of the "binder" chain would not be necessarily accurately predicted from amino acid sequence alone (as both AF2 and RF2 cannot generally accurately predict native protein structures without multiple sequence alignment (MSA)/template inputs [12, 22]). Rather than providing MSA information for these regions, we instead simply provided the structure of the "binder framework" as a template input to the model, in the two-dimensional track of RF2. Note that this does not provide the rigid-body orientation of the "binder" chain with respect to the "target" chain, and does not provide information about the interface loop structure.

#### 5.9 Generation of Negative Examples

As well as predicting structure, RF2 predicts whether two chains bind (with the pBind prediction head; Section 5.1). As such, it is trained both with interacting ("positive") and non-interacting ("negative") pairs of proteins. To make use of this predictor in the context of antibody design filtering, during antibody fine-tuning, RF2 was similarly trained with both positive and negative examples (sampled with 50% probability). The negative examples were generated stochastically during training. Three methods for generating such negative examples were explored, with subtle modifications for the three datasets. These are detailed in Sections 5.10 - 5.12

#### 5.10 Mis-Matching Binder/Target Chains

For antibody-antigen pairs, negative examples were generated by sampling an antibody from a different antibody cluster (clusters generated by $< 70\%$ CDR sequence similarity; Section 1.1). The sampled cluster is always from the same training/test partition. To be sufficiently confident the "negative" antibody would not bind to the target chain, these antibodies were sampled only from the set of antibody structures where the antigen was present, and where the antigen shared $< 20\%$ sequence similarity (by MMseqs2) to the "positive" antigen.

For the TCR set, a "negative" TCR was sampled by sampling a different cluster (MHC subtype) from the same train/test partition, and subsequently sampling a TCR sequence from that cluster.

For the loop-mediated interaction set, a "negative" binder sequence was sampled from a different cluster (but same train/test partition) but same species.

#### 5.11 Swapping CDR Loops

The second way we explored to generate negative examples for the antibody and TCR sets was to swap key interacting loops with unrelated sequences. We assume that swapping the H3 CDR loop of antibodies, and the $\alpha 3 + \beta 3$ loops of TCRs would abolish binding, given that these typically mediate the key interactions between antibodies/TCRs and antigens [23]. For each of these loops, a "negative" was found that met the following criteria:

1. The same length as the "positive" loop
2. $< 40\%$ sequence similarity to the "positive" loop
3. From the same train:test partition

Note that for the $\alpha 3 + \beta 3$ loops, the two replacement CDRs were not necessarily from the same TCR, but each individually met the above criteria.

#### 5.12 De Novo Miniprotein Binder Set

A third way we explored for providing the model with negative examples was to task the model with predicting whether de novo designed miniprotein binders bound experimentally (based on in-house yeast surface display data). When receiving one of these examples, the model was provided the full target sequence and structure, interface hotspots on the target derived from the AF2-predicted miniprotein binder-target dock, and the sequence of the miniprotein binder. A loss was applied on whether the model correctly predicted the miniprotein binder as binder or nonbinder, no loss was applied on whether the model predicted the complex structure correctly (as no "true" structure exists for these examples).

#### 5.13 Sampling of Training Examples from Clusters

During training, clustering is used to prevent over-representation of certain highly similar proteins in the training set. For "positive" examples, for each training example, a cluster is randomly sampled (without replacement within a training epoch), and from that cluster a specific structure is sampled. For the "negative" examples generated by random pairing of an unrelated "binder" and “target”, the “negative” cluster was first sampled, and then a replacement “binder” was sampled from the set of structures within this cluster. This similarly prevents over-representation of any specific cluster in the negative training set.

#### 5.14 Losses

The losses used during fine-tuning of RF2 are the same as those used to train RF2 originally. These losses are presented in Table 3. For positive training examples, structural losses ($\mathcal{L}_{\text{dist}}$, $\mathcal{L}_{\text{FAPE}}$, $\mathcal{L}_{\text{bond}}$, $\mathcal{L}_{\text{vdW}}$, $\mathcal{L}_{\text{pAE}}$) were applied over all residues resolved in the structure, over the whole complex. For negative examples, structural losses were only applied over the individual “binder” and “target”, but not across the complex (as there is no “correct” rigid body orientation). In cases where negative examples were generated by swapping key CDR loops or residues, no structural loss was applied over these residues.

#### 5.15 RoseTTAFold2 Fine-Tuning Settings

We fine-tuned RF2 in two stages:

In the first stage, we fine-tuned RF2 starting from published RF2 weights [8]. We trained on 45% SabDab examples, 10% TCR examples, and 45% loop-mediated interfaces from the PDB (see Section 1 for more information on the datasets). We trained using the updated pBind auxiliary head (Algorithm 5) and with 50% of examples being mis-matched binder/target chains (Section 5.10). We used a pseudo batch size of 64 and a learning rate of 0.001 that was decayed by 0.95 after every 10,000 optimizer steps. We used the losses outlined in Table 3. RF2 was trained for 400 epochs with 512 examples per epoch. All other training settings used are the same as in the published RF2.

In the second stage, we started from the set of weights output by the first stage, and trained on 45% SabDab examples, 10% TCR examples, 10% loop-mediated interfaces from the PDB, and

| Loss | Description | Weight |
|---|---|---|
| $\mathcal{L}_{\text{dist}}$ | Loss on distogram | 1.0 |
| $\mathcal{L}_{\text{FAPE}}$ | Frame Aligned Point Error loss on predicted structure, split 50:50 on backbone FAPE and All-Atom FAPE | 10.0 |
| $\mathcal{L}_{\text{pLDDT}}$ | Loss on accuracy estimation (pLDDT) | 1.0 |
| $\mathcal{L}_{\text{bond}}$ | Loss on bond geometry | 0.02 |
| $\mathcal{L}_{\text{vdW}}$ | Loss on van der Waals energy, as estimated from the Rosetta Lennard-Jones potential (LJ, 12-6). | 0.02 |
| $\mathcal{L}_{\text{MLM}}$ | Masked language modelling loss (note no sequence masking occurs during antibody fine-tuning) | 3.0 |
| $\mathcal{L}_{\text{pAE}}$ | Loss on the predicted alignment error | 0.01 |
| $\mathcal{L}_{\text{bind}}$ | Loss on whether a complex is a positive or negative example | 1.0 |

Supplementary Methods Table 3: Losses used to fine-tune RF2 on antibodies.

35% de novo miniprotein binder examples (Section 5.12). We trained with a psuedo batch size of 32. The other training settings were the same as the first stage. For this stage, we trained for 207 epochs with 512 examples per epoch.

### 6 Computational Methods

#### 6.1 Attempting Antibody Design in Vanilla RFdiffusion

To confirm that fine-tuning of RFdiffusion was required for antibody design, we attempted to design antibodies in vanilla RFdiffusion. We tried two strategies. In the first, we provided the sequence of the antibody framework to the version of RFdiffusion that can condition on sequence input alone [24]. Additionally, we provided the target structure and "hotspot" information specifying the epitope. The second strategy was to additionally provide "fold" information about the antibody

| Input Name (Shape) | Description |
|---|---|
| msa_masked (1,1,L,48) | The sequence (20aa, zeros (2), repeat aa (20), zeros (6)) |
| msa_full (1,1,L,25) | The sequence (20aa, zeros (5)) |
| seq (1,L) | The sequence as an index tensor |
| idx_pdb (L) | The integer index of each residue. Used to create a positional encoding. A 200 residue offset is inserted to indicate a chain break. |
| t1d (T,L,23) | The 1-D features of the templated regions. T=1 if no binder template information is provided. T=2 if binder template information is provided. (20 amino acids, mask (1), template-match confidence (1), hotspot boolean (1)) |
| t2d (T,L,L,44) | The 2-D features of the templated regions. T=1 if no binder template information is provided. T=2 if binder template information is provided. (36 distance bins ($2 - 20$Å, 0.5Å bins) + 1 final distance bin ($>20$Å), angle maps (sine and cosine of omega, theta and phi angle) (6), mask (1)) fine-tuning) |
| xyz_t (T,L,27,3) | The 3-D coordinates of the templated regions. T=1 if no binder template information is provided. T=2 if binder template information is provided. (4 backbone atoms N, CA, C, O (4), 10 heavy sidechain atoms (zeros if not present), 14 hydrogen atoms (zeros if not present)). This feature is immediately converted to a distogram and anglegram by the model so only the backbone atoms are used. |
| alpha_t (T,L,30) | Sidechain torsion information for the templated regions. T=1 if no binder template information is provided. T=2 if binder template information is provided. Initially T, L, 10, 2, with sine and cosine of (omega, phi, psi angles (3), (up to) 4 torsion angles, $C_\beta$ bend (1), $C_\beta$ twist (1), $C_\gamma$ bend (1)). This is concatenated with a mask (T, L, 10, 1) indicating which torsion angles are present for a given amino acid, and reshaped to T, L, 30. When training with no sidechain information, all entries in this feature are zeros and are masked. |
| t2d (T,L,L,44) | The 2D features of the templated regions. T=1 if no binder template information is provided. T=2 if binder template information is provided. (36 distance bins ($2 - 20$Å, 0.5Å bins) + 1 final distance bin ($>20$Å), angle maps (sine and cosine of omega, theta and phi angle) (6), mask (1)) fine-tuning) |
| same_chain (L,L) | 2D binary feature indicating whether a pair of residues belong to the same chain or not. |
| xyz_prev (L,27,3) | The 3-D recycling information. (4 backbone atoms N, CA, C, O (4), 10 heavy sidechain atoms (zeros if not present), 14 hydrogen atoms (zeros if not present)) |
| msa_prev (L,$C_m$) | The MSA embedding recycling information. Zeros at the first cycle and then passed between recycles. $C_m = 256$ |
| pair_prev (L,L,$C_p$) | The 2-D embedding recycling information. Zeros at the first cycle and then passed between recycles. $C_p$ = 128 |
| state_prev (L,$C_s$) | The 1-D embedding recycling information. Zeros at the first cycle and then passed between recycles. $C_s$ = 16 |

Supplementary Methods Table 4: Description of Features Input to Fine-Tuned RF2.

framework (secondary structure and “block-adjacency” inputs [9]). 100 designs were made to four separate targets, for both VHHs and scFvs.

#### 6.2 In Silico Analysis of True/Decoy Discrimination in RoseTTAFold2

To determine the extent to which fine-tuned RF2 can distinguish between true (native) antibody-antigen pairs vs decoy pairs (Extended Data Fig. 2), we took the validation set described above (which shares no target homology to the training set), and predicted the structures of either true antigen-target pairs, or 5 randomly selected antibody-decoy pairs. In the case of decoys, the hotspots for the decoy target with its native antibody were extracted and provided, along with that decoy target structure and the (now non-matching) antibody sequence. 10 recycles were used in RF2, with the sample with the highest pLDDT saved.

To assess specificity of designed VHHs (Extended Data Fig. 4) and scFvs (Extended Data Fig. 6) for the targets they were designed to bind, we performed the same test; predicting the structure of designed sequences against either the correct or decoy targets (500 designs were generated for each target with identical settings, with one ProteinMPNN sequence designed per backbone). For VHHs, we performed this on the set of targets for which binders were successfully identified. For scFvs, we performed this analysis both on the set of VHH-successful targets, and also a second set of five unrelated epitopes. We plotted the proportion of designs with RF2 pAE $< 10$, normalized to the proportion of pAE $< 10$ predictions in the true VHH-target case for that given target. 10 recycles were used in RF2, with the sample with the highest pLDDT saved.

#### 6.3 Comparing RoseTTAFold2 Antibody Monomer Prediction to IgFold

To compare the performance of RF2 at predicting antibody monomer structures to the performance of IgFold [25], we used the part of the validation set released after the fine-tuned RF2 date cutoff

(January 13th, 2023; 104 structure, of which 29 are VHHs), which cannot be in either the RF2 nor IgFold training dataset, nor share homology to their training sets. For RF2, we predicted the structure using 10 recycles. For IgFold, we used the recommended settings including Rosetta refinement. Backbone R.M.S.D. to the native structure was performed both over the whole Fv ($\leq$ residue 110 of the heavy chain; $\leq$ residue 105 of the light chain; Chothia numbering [2]) and over just the CDR H3 loop (as defined by the Chothia numbering system).

#### 6.4 Comparing Designed Interfaces to Native Interfaces

To assess the quality of designed antibody interfaces (orthogonally to assessment with fine-tuned RF2), we used Rosetta [26]. For the native set, we took all non-redundant VHH complexes (1296 structures) and 1000 non-redundant randomly sampled Fv complexes from the PDB. We compared these to the VHHs (Extended Data Fig. 4C) and scFvs (Extended Data Fig. 6D) described in Extended Data Fig. 4B and 6C respectively. These were predicted with RF2 and filtered to designs with R.M.S.D. to the design model < 2Å and pAE < 10. These two sets of structures were relaxed over the interface with Rosetta FastRelax [26]. Rosetta ddG was subsequently calculated over the interface. Additionally, the Surface Aggregation Propensity (SAP) score, developed originally to measure solubility/developability of antibodies [27, 28], was measured.

#### 6.5 Assessment of Designed VHH Structural Similarity to the PDB

The structural similarity of the highest affinity VHH to each target and the structurally-characterized VHH to influenza HA was assessed. For each target, PDB entries containing the target were identified using Blastp [29, 30] against the entire PDB, blast hits with $\geq 25$ "query coverage" and $\geq 90$ "percent identity" were accepted. Entries were then filtered to only those containing a chain within the 30% sequence similarity cluster of immunoglobulin domains (the first and largest cluster in the 30% clustering of all sequences in the PDB provided at

| Target | PDB Accession | Hotspot Residues (<chain><idx>) |
|---|---|---|
| HIV Env | 2NY7 | A371, A375, A435, A475 |
| SARS-CoV-2 RBD | 6M0J | B492, B493, B494, B495, B496, B497 |
| RSV-F Site I | 7LVW | D469, D384 |
| RSV-F Site III | 7LUC | T305, T456 |
| Influenza Hemagglutinin | 5VLI | B146, B170, B177 |
| TcdB | 6C0B | A1433, A1435, A1437, A1438, A1493 |
| IL-7 Receptor $\alpha$ | 3DI3 | B81, B139, B192 |

Supplementary Methods Table 5: VHH Design Campaigns.

https://cdn.rcsb.org/resources/sequence/clusters/clusters-by-entity-30.txt). This truncated set of complexes was searched visually for designs showing similarity to the design model. No significant structural similarity was found. Results of this analysis are shown in Extended Data Fig. 5.

To assess similarity of the CDR loops to CDR loops in the PDB, designed VHH sequences were queried against the PDB using Blastp [29, 30]. The top identified hit was then aligned to the designed CDRs, and sequence similarity ($N\_Matching/CDR\_Length$) of each CDR was measured.

#### 6.6 VHH Design Campaigns

We designed VHHs against 6 targets in total. These, along with their PDB accessions and "hotspot" residues used to define the epitope are detailed in Table 5.

#### 6.7 TcdB scFv Design Campaigns

We designed two scFv libraries against TcdB, the target PDB accessions and "hotspot" residues used to define the epitope are detailed in Table 6.

| Target | PDB Accession | Hotspot Residues (<chain><idx>) |
|---|---|---|
| scFv TcdB - Unique Pairing | 7ML7 | A1816, A1818, A1819, A1823, A1831 |
| scFv TcdB - Combinatorially Assembled | 6C0B | A1433, A1435, A1437, A1438, A1493 |

Supplementary Methods Table 6: TcdB scFv Design Campaigns.

#### 6.8 Phox2b Peptide scFv Design Campaigns

To design specific scFv binders against PHOX2B:HLA-C*07:02 in the absence of an experimentally-determined structure for this pMHC complex, we first used AlphaFold2 to predict the PHOX2B:HLA-C*07:02 structure. We used a previously published variant of AlphaFold2 fine-tuned on pMHC structures and binding data [31]. We also used a slightly different AlphaFold2 variant similarly fine-tuned on pMHC structures and binding data, but with more equal weighting of the fine-tuning training data across different peptide backbone conformations in PDB structures [32]. The two models' predicted structures for this pMHC differed significantly in the peptide backbone's bond angles, with each pointing the critical I5 and R6 residues in opposite directions towards each $\alpha1$ and $\alpha2$ groove of HLA-C*07:02. We chose to design antibodies were designed against both pMHC structures. Hotpsots were sampled randomly from the set of all peptide residues and the residues on the MHC adjacent to the peptide. All designs, regardless of target hotspot or target AlphaFold2 model, were mixed together in terms of structure-based library construction.

#### 6.9 In Silico Analysis of Combinatorially-Assembled Heavy and Light Chains

To assess the predicted quality of combinatorially assembled heavy and light chains, we clustered designs by structural similarity and predicted the complex structure of a randomly sampled subset of heavy and light chains, along with the target. TcdB scFv designs, pre-filtered with fine-tuned

RoseTTAFold2, were compared structurally through pairwise TM-align

#### 6.10 Retrospective Analysis of AF3 iPTM Predictivity of Binders

##### 6.10.1 VHH

Single seed AlphaFold3 predictions using version 1 were run locally using the default MSA and template generation pipeline for the target (JackHMMR [33]) and template generation only for the VHH (the MSA field was left empty). The input targets were cropped for computational efficiency. See Extended Data Fig. 18A,B. The sequences of the input targets are shown in 7

##### 6.10.2 scFv

scFv structures were predicted using AlphaFold3 version 2 with MSA (JackHMMR) and template for the target structure and only template for the heavy and light chains of the scFv. Linker sequences were not included between the heavy and light chain of the scFv during prediction, as thus they were treated as separate chains.10 seeds were used for each scFv prediction in the initial design library and the max iPTM score was taken from across the 10 seeds. See Extended Data Fig 18C,D.

| **Target** | **Sequence** |
|---|---|
| TcdB | FVSLTFSILEGINAIIEVDLLSKSYKLLISGELKILMLN SNHIQQKIDYIGFNSELQKNIPYSFVDSEGKENGF INGSTKEGLFVSELPDVVLISKVYMDDSKPSFG YYSNNLKDVKVITKDNVNILTGYYLKDDIKI SLSLTLQDEKTIKLNSVHLDESGVAEILKFMNRK GSTNTSDSLMSFLESMNIKSIFVNFLQSNIKFI LDANFIISGTT |
| IL-7R$\alpha$ | DYSFSCYSQLEVNGSQHSLTCAFEDPDVNTTNLEF EICGALVEVKCLNFRKLQEIYFIET KKFLLIGKSNICVKVGEKSLTCKKI DLTTIVKPEAPFDLSVVYREGAND FVVTFNTSHLQKKYVKVLMHDVAY RQEKDENKWTHVNLSSTKLTLLQ RKLQPAAMYEIKVRSIPDHYFKGF WSEWSPSYYFRTP |
| SARS-CoV-2 RBD | NLCPFGEVFNATRFASVYAWNRKRISNCVAD YSVLYNSASFSTFKCYGVSPTKLNDLCFTNV YADSFVIRGDEVRQIAPGQTGKIADYNYKLP DDFTGCVIAWNSNNLDSKVGGNYNYLYRLFR KSNLKPFERDISTEIYQAGSTPCNGVEGFNC YFPLQSYGFQPTNGVGYQPYRVVVLSFELLH APATVCG |
| Influenza HA | TICIGYHANNSTDTVDTVLEKNVTVTHSVNLL EDSHNGKLCRLKGIAPSWSYISSVSSSRGFGS GIITSNASMHECNTKCQTPLGAINSSLPYQNI HPVTIGECPKYVRSAKLRMVTGLRNIPGLFGA IAGFIEGGWTGMIDGWYGYHHQNGSGYAADQK STQNAINGITNKVNTVIEKMNIQFTAVGKLNK KVDDGFLDIWTYNAELLVLLENERTLDFHDSN VKNLYEKVKSQLKAKEIGNGCFECDNECMESV RNGTYDY |

Supplementary Methods Table 7: Sequences of Targets used in AF3 Analysis

### Part II

### Experimental Methods

#### 7 VHH Yeast Surface Display

##### 7.1 Library Preparation

Designed VHH sequences were padded to the same length via poly-serine at the C-terminus and then codon optimized for expression in *S. cerevisiae* using DNAWorks 2.0 [34]. 5' and 3' adapter sequences encoding glycine-serine linkers were added to enable subpool amplification (5' adapters: TCGTCTGGTAGTTCAGGC or GGTGGATCAGGAGGTTCG, 3' adapters: GGAAGCGGTG-GAAGTGG or GGTTCTAGTGGCTCATC).

The libraries were amplified using Kapa Hifi polymerase (Roche) in 25$\mu$l reactions first to determine how many cycles to reach half of maximum yield then a second production run which was subsequently loaded onto an agarose gel and purified using a Qiaquick kit (Qiagen). This purified product was then reamplified a second time to generate 6$\mu$g of DNA insert which was co-transformed into EBY100 with 3$\mu$g of linearized pETcon3 using a previously described electroporation protocol [35]. Transformed yeast cultures were grown in C-Trp-Ura media (with 2% glucose w/v) at 30°C.

##### 7.2 Cell Sorting

The first round of FACS was with only anti-myc FitC to enrich the yeast population expressing myc (at the C-terminus of each design) on the cell surface. 10ml of SGCAA was inoculated with 200$\mu$l of yeast culture and grown at 30°C for 18 hours. Cells were harvested by centrifugation at 4000 x g for 4 minutes and then resuspended in 2ml of PBS with 1% BSA (PBSF). 200$\mu$l of the resuspended culture was then washed once in PBSF prior to resuspension in 50$\mu$l PBSF with 1% anti-myc FitC antibody (ICL CMYC-45F) for 10 min at room temperature prior to washing 3 x in PBSF and resuspending in 1000$\mu$l immediately prior to sorting. Approximately 10 million cells were sorted during the expression sort and grown in CTUG media overnight.

Sort 2 and sort 3 followed the same method as sort 1 but were resuspended in 50$\mu$l in PBSF with 1% anti-myc FitC and 1$\mu$l of biotinylated target protein in complex with streptavidin-PE (Life Technologies S866). Cells were incubated at room temperature for 1 hour in the presence of target protein prior to washing thrice with PBSF and again resuspending in 1000$\mu$l PBSF immediately prior to sorting. Here only the double positive (+FitC and +PE) were collected and expanded for the next round of sorting.

The final sort was a titration sort where cells were first stained with target protein alone starting at 1000$\mu$l and then a 3-fold dilution series. After a one hour incubation at room temperature, the cells were washed once in PBSF and then resuspended in 50$\mu$l of PBSF with 1% anti-myc FitC and 1% streptavidin-PE. After a ten minute incubation on ice, these cells were washed thrice in PBSF before resuspension in 1000$\mu$l immediately prior to sorting. Again, only the double positive population was collected.

### 8 scFv Assembly and Screening

#### 8.1 Assembly of scFvs with Exact Pairing of Heavy Chains with their Corresponding Light Chains

##### 8.1.1 Concept

Here, we develop a multi-step assembly strategy for assembling large (tens of thousands) libraries of scFvs, where the designed heavy and light chains from a single design assemble uniquely, with (theoretically) no assembly of heavy and light chains from different designs. Additionally, to minimize the variability that is introduced by arbitrary choices of chain ordering (heavy-light vs light-heavy) and inter-chain linker, we built a system that could assemble in either order with multiple different linkers. The approach builds upon previous work, utilizing overlap PCR [36] and GoldenGate assembly [37], as well as large sets of previously-published orthogonal DNA barcodes [38]. The approach is schematically depicted in Extended Data Figs. 19 and 20.

##### 8.1.2 Oligonucleotide Design

The goal of this protocol was to order the minimal-necessary section of DNA to encode the designed heavy and light chains. We therefore "fix" the scFv framework, and order only the designed segment (CDR1-CDR3). Note, the interspersing framework sequences (FR2, FR3) are not designed. The heavy- and light-chain fragments must be ordered on separate DNA fragments (because of oligonucleotide sythesis limitations), and thus, must be brought together. Unlike in de novo proteins, where sufficient sequence diversity between designs makes assembly with overlap PCR trivial (assembly occurs over a unique part of the designed sequence) [36], the high degree of sequence homology in antibody sequences makes overlap PCR impracticable in this manner. We therefore use DNA barcodes, specially designed to be orthogonal to each other [38], to assemble the two chains. Each designed heavy- light- chain pair receive the same barcode, such that (theoretically) only they should assemble in the overlap PCR step.

To allow for assembly in either order (H-L vs L-H), we add two sets of unique barcodes to each oligo; one on each side of the designed sequence. This permits the overlap PCR to occur in either order. Finally, we control which fragment we amplify through the use of USER cleavable primers [39]. This also limits the number of PCR steps to *2 orders* × *number of subpools*. In other words, the oligonucleotides are ordered as:

< *USER primer 1* >< *barcode 1* >< *primer 2* >< *designed fragment* >< *primer 1* >< *barcode 2* >< *USER primer 2* >

Which, with USER primer 1 and primer 1 can amplify:

*< USER primer 1 >< barcode 1 >< primer 2 >< designed fragment >< primer 1 >*

which post USER-cleage will reveal:

*< barcode 1 >< primer 2 >< designed fragment >< primer 1 >*

with barcode 1 available for assembly PCR, with this designed fragment at the C-terminus of the combined oligonucleotide.

Alternatively, with USER primer 2 and primer 2, we amplify:

*< primer 2 >< designed fragment >< primer 1 >< barcode 2 >< USER primer 2 >*

which post USER-cleavage will reveal:

*< primer 2 >< designed fragment >< primer 1 >< barcode 2 >*

with barcode 2 available for assembly PCR, with this designed fragment at the N-terminus of the combined oligonucleotide.

##### 8.1.3 Choice of Barcodes

Barcodes were chosen from a previously described set of designed orthogonal 25bp DNA barcodes [38]. Because of oligonucleotide-length limitations, only the central 18bp were taken (sufficient for PCR amplification). Further, we filtered this set of barcodes such that all barcodes had a predicted melting temperature (Tm) between 58-61°C (as assessed by the Wallace method).

##### 8.1.4 Cloning Step 1

By amplifying the two fragments (heavy and light chain), and subsequently combining them with overlap PCR, we end up with either:

< *primer* >< *designed H fragment* >< *unused primer* >< *barcode* >< *unused primer* >< *designed L fragment* >< *primer* >

or

< *primer* >< *designed L fragment* >< *unused primer* >< *barcode* >< unused primer >< *designed H fragment* >< *primer* >

We then clone this into a custom vector using GoldenGate cloning [37]. This vector brings FR1 and FR4 of the N- and C-terminal chain, respectively.

##### 8.1.5 Cloning Step 2

After a first cloning step, we use a second GoldenGate step to remove the “unused primers” and the “barcode”, and replace this with FR4 and FR1 of the N- and C-terminal chain, along with a linker between them. This is a pooled reaction, so multiple linker sequences can be mixed to increase library complexity and hopefully reduce the bias induced by linker choice.

##### 8.1.6 Technical Protocol

Each qPCR reaction is performed in duplicate. The first reaction is used to assess the amplification dynamics for each subpool reaction, such that in the second reaction, DNA can be removed in the log phase of PCR amplification, which limits off-target amplification [36].

**qPCR1:** The single stranded oligonucleotide array is thawed, and diluted to 2.5ng/$\mu$l. This is amplified with the KAPA HiFi HotStart Uracil+ kit with Eva green used to monitor amplification. For each subpool, forward and reverse primers are added at 10$\mu$M. Note that the primer amplifying the USER adapter contains the necessary uracils for downstream cleavage. Amplification was performed with 95°C for 2 minutes, followed by the subpool-appropriate number of cycles of 98°C (20s) - 63°C (15s) - 72°C (45s).

**PCR cleanup 1:** PCR cleanup was performed in parallel with magnetic beads (AMPure XP), according to the manufacturer's instructions. DNA was eluted in Qiagen buffer EB.

**USER primer cleavage and end repair:** USER primer cleavage was performed with addition of 2$\mu$l of USER enzyme (NEB) was added to each qPCR product, and incubated for 37°C (15 minutes) followed by 22°C (15 minutes). 1X NEBNext End Repair Buffer and 5$\mu$l NEBNext End Repair Enzymes were added (100$\mu$l total reaction volume), and incubated at 20°C for 30 minutes.

**PCR cleanup 2:** PCR cleanup was performed in parallel with magnetic beads (AMPure XP), according to the manufacturer's instructions. DNA was eluted in Qiagen buffer EB.

**qPCR2:** 50ng each of cleaned heavy-chain fragments and their corresponding light-chain fragments were mixed. 4$\mu$l of this mixed pool was amplified with the KAPA HiFi HotStart kit (without uracil), with Eva green used to monitor amplification. For each subpool, forward and reverse primers are added at 10$\mu$M. Note that these primers do not contain uracil, and are used to perform overlap PCR between the two fragments. Amplification was performed with 95°C for 2 minutes, followed by the subpool-appropriate number of cycles of 98°C (20s) - 63°C (15s) - 72°C (45s).

**Gel Purification:** Amplified fragments were run on a 1% agarose gel with 1X SYBR Safe used to stain the DNA. Fragments at the appropriate length were subsequently purified using the Qiagen gel extraction kit.

**GoldenGate Cloning 1:** All subpools of the same order (H-L and L-H) are pooled (adding equimolar concentrations of each to the pool). An insert:vector ratio of 3:1 is used, aiming for 100-300ng of insert DNA per reaction. 1X freshly-thawed T4 ligase buffer (Thermo) is combined with 1.2$\mu$l BsaI (H-L order) or 2.4$\mu$l BsmBI (L-H order), along with 2$\mu$l T4 ligase (Thermo) (50$\mu$l total reaction volume). The reaction was incubated for 37°C for 1h, followed by 60°C for 5 minutes. PCR purification (Qiagen; QIAquick) was performed, with elution in ultrapure water. 4 parallel reactions were performed and pooled.

**E. coli Electroporation:** Electrocompetent NEB 5-alpha Electrocompetent E. coli were thawed on ice just prior to transformation. 1$\mu$g assembled DNA was added to 25$\mu$l E. coli, and electroporated with the BioRad Pulser, according to the manufacturer's instructions. Immediately post-electroporation, cells were gently resuspended in SOC medium and left shaking at 37°C for 1h. E. coli were then grown overnight in Lysogeny broth (LB) + 50mg/ml carbenicillin at 37°C.

**Midiprep:** 50ml of culture was midiprepped using the Qiagen Plasmid Plus Midi kit, according to the manufacturer's instructions.

**GoldenGate Cloning 2:** Linker sequences flanked by FR4 (upstream) and FR1 (downstream), along with the second GoldenGate enzyme (BsmBI for H-L; BsaI for L-H), were amplified by qPCR as described above. GoldenGate reactions were subsequently setup with a 3:1 insert:vector ratio, as above. The reaction was incubated at 37°C for 1h, followed by 55°C for 1h and 60°C for 5 minutes. The 55°C incubation is necessary because the lethal gene which eliminates E. coli carrying misassembled plasmid is excised during GoldenGate assembly 1. The extra, high-temperature incubation step enables optimal BsmBI digestion while mostly inactivating T4 ligase, ensuring that any plasmids where BsmBI sites were not excised and replaced with linker are converted to linear DNA. This acts as a form of selection against misassembled plasmids, since only correctly assembled, circular plasmids will be taken up by E. coli during the electroporation step that follows. PCR cleanup (Qiagen, QIAquick) was performed, with elution in ultrapure water. E. coli electroporation and midiprep were performed as above to obtain enough correctly assembled, purified plasmid for yeast electroporation.

**Yeast Electroporation:** Yeast electroporation was performed based on a protocol adapted from [40] and [41]. Briefly, 3-4$\mu$g of assembled DNA were combined with sheared salmon sperm in a 1:50 ratio, and this mixture was electroporated into 400$\mu$L of freshly permeabilized electrocompetent EBY100 *S. cerevisiae* cells suspended in electroporation buffer (1M sorbitol + 1mM $CaCl_2$). Electroporation was performed at 2500V using 2mm gap electroporation cuvettes with the BioRad Pulser, according to the manufacturer's instructions. Immediately post-electroporation, cells were gently resuspended in recovery medium (yeast peptone dextrose (YPD) broth + 0.5M sorbitol) and recovered by shaking at 250 rpm at 30°C for 1 hour. Cells were pelleted and resuspended in 100mL C-Trp-Ura medium with 2% (w/v) glycerol (CTUG) and incubated for 2-3 days by shaking at 250 rpm at 30°C.

#### 8.2 Combinatorial Assembly of scFvs from Heavy and Light Chains from Designs within Structural Clusters

##### 8.2.1 Concept

In addition to the unique-chain pairing strategy we detail above, which relies on 400bp oligonucleotides, we also developed a strategy for pairing heavy and light chains from designs within specific structural clusters (Extended Data Fig. 21). This can be performed with (cheaper) 300bp oligonucleotides, although the ability to sample both H-L and L-H chain ordering is lost. This combinatorial assembly strategy permits the massive increase in library size ($\textit{library complexity} = \textit{N clusters} \times \textit{cluster size}^2$), while retaining high in silico success rates (Extended Data Fig. 22A).

##### 8.2.2 Oligonucleotide Design

The concept closely follows that of the unique assembly protocol, where overlap PCR combines appropriate heavy and light chains, and two sequential GoldenGate assembly steps are used to 1) add this fragment to a vector, and 2) add the remaining framework regions and linker. The key differences are that: One chain ordering (H-L or L-H) is chosen for each structural cluster. This is chosen based on which N- and C-terminus pair is closer (smaller is preferred) No USER cleavable primers are used. Instead, the inner and outer primer specify the design subpool/cluster. Thus, the number of initial qPCRs performed is 2 chains x N clusters. These primers are used as the complementary region for the overlap PCR step.

The oligonucleotides are ordered (for H-L ordering) as:

< *primer 1* >< *H designed fragment* >< *primer 2* >

and

< *primer 2* >< *L designed fragment* >< *primer 3* >

or (for L-H ordering):

< *primer 1* >< *L designed fragment* >< *primer 2* >

and

< *primer 2* >< *H designed fragment* >< *primer 3* >

##### 8.2.3 Cloning Step 1

By amplifying the two fragments (heavy and light chain), and subsequently combining them with overlap PCR, we end up with either:

< *primer 1* >< *H designed fragment* >< *primer 2* >< *L designed fragment* >< *primer 3* >

or

< *primer 1* >< *L designed fragment* >< *primer 2* >< *H designed fragment* >< *primer 3* >

We clone this into a custom vector using GoldenGate cloning [37]. This vector brings FR1 and FR4 of the N- and C-terminal chain, respectively.

##### 8.2.4 Cloning Step 2

After a first cloning step, we use a second GoldenGate step to remove the central overlap region, and replace this with FR4 and FR1 of the N- and C-terminal chain, along with a linker between them. This is a pooled reaction, so multiple linker sequences can be mixed to increase library complexity and hopefully reduce the bias induced by linker choice.

##### 8.2.5 Technical Protocol

The protocol mirrors that of the unique pairing library assembly strategy in Section 8.1.6, except that there is no USER step. Hence, we proceed directly from qPCR into PCR cleanup and overlap qPCR2.

#### 8.3 Yeast Display Screening of scFvs

For each library, *S. cerevisiae* cultures transformed with the assembled scFv plasmid library were evaluated with rounds of fluorescence-assisted cell sorting (FACS) as described in a previous publication [18]. We determined the constructs expressed on sorted cells with next-generation sequencing (NGS) conducted on the MiSeq (Illumina).

An expression sort was conducted before performing avidity sorts. Cells were washed and then labeled with anti-c-myc-fluorescein isothiocyanate (FITC) for 15 minutes prior to the sort. All cells with high FITC signal were sorted, and this population was evaluated with NGS to determine the starting library population.

Enrichment was performed first with two rounds of avidity sorts, each conducted with a target concentration of 1 $\mu$M. These sorts were conducted using biotinylated target tetramerized on streptavidin-R-phycoerythrin (SAPE) prior to yeast incubation. Libraries failed if no cells were collected during the second avidity sort. For libraries with hits at 1 $\mu$M with avidity, multiple rounds of sorts without avidity were then conducted with decreased target concentration in each round.

Final binder sequences were determined by yeast colony PCR and nanopore sequencing. Yeast colonies were obtained by plating diluted CTUG cultures of sorted cells on C-Trp-Ura agar plates and incubating plates at 30°C. Colony PCR was conducted by combining 25 $\mu$L 2x Phire Plant Direct PCR Master Mix (Thermo Fisher Scientific), 2.5 $\mu$L forward and reverse primers (at 10 $\mu$M), 1$\mu$l CTUG culture, and 19 $\mu$L nuclease-free water. This mixture was incubated at 98°C for 5 minutes, then 40 cycles of (98°C for 5 seconds, 66.2°C for 5 seconds, 72°C for 20 seconds), then 72°C for 1 minute. Samples were purified using AMPure XP reagent (Beckman Coulter Life Sciences) according to manufacturer's directions before submission for nanopore sequencing (Plasmidsaurus, Seattle).

Sequences for scFv4, scFv5, and scFv6 were discovered in this manner. These validated sequences were produced as soluble protein commercially (GenScript). Designs were expressed in CHO-S cells and then purified by Ni-NTA affinity chromatography followed by size exclusion chromatography using high-performance liquid chromatography (SEC-HPLC).

### 9 Biochemical Characterization

#### 9.1 VHH Expression and Purification

For soluble protein expression and purification, each design was ordered as a linear DNA fragment (eblocks from Integrated DNA Technologies) with flanking BsaI cut sites and golden gate compatible overhangs, and then cloned into LM627 [42]. The conditions for each cloning reaction were as follows:

- 225nl water
- 100nl 10x T4 buffer (New England Biolabs B0202S)

- 150nl LM627 (100ng/$\mu$l) (Addgene 191551)
- 50nl BsaI (1.2U/rxn) (New England Biolabs R3733L)
- 100nl T4 ligase (40U/rxn) (New England Biolabs M0202L)
- 375nl DNA insert (4ng/ul)

After 30 min at 37°C, the reaction was transformed into BL21 *E. coli* followed by overnight growth in 4x1ml cultures in autoinduction media. After 20h at 37°C, cultures were harvested by centrifugation and lysed in 400$\mu$l BPER with lysozyme (0.1mg/ml), benzonase (25U/ml) and PMSF (1mM). Lysates were then clarified by centrifugation and applied to 50$\mu$l of Ni-NTA resin. After 3 washes with 20mM Tris, 300mM NaCl, 25mM imidazole, pH 8, proteins were eluted in 200$\mu$l of elution buffer (20mM Tris, 300mM NaCl, 300mM imidazole). Each protein then underwent size exclusion chromatography on an S75 5-150 column (Cytiva) in HBS-EP+ (0.01M HEPES pH 7.4, 0.15M NaCl, 3mM EDTA, 0.005% v/v Surfactant P20).

#### 9.2 scFv Expression and Purification

scFvs were expressed by Genscript in Chinese Hamster Ovary (CHO) cells with a C-terminal His-tag. scFvs were affinity purified with Ni-NTA chromatography with AmMag Ni magnetic beads. scFvs were subsequently further purified using high performance liquid chromatography using a Phenomenex Biozen column into Phosphate Buffered Saline (PBS).

#### 9.3 TcdB Expression

Plasmid pHis1522 encoding his-tagged TcdB VP10463 was a gift from Hanping Feng (University of Maryland Dental School, Baltimore, MD, 21201, USA.). The plasmid for TcdB 027 in pHis1522 was synthesized by Genscript USA. TcdB was purified from Bacillus megaterium (MoBiTec, Germany) carrying the vector pHIS1522 encoding C. difficile TcdB VP10463 or 027 fused to a C-terminal His tag.

Overnight starter cultures of B. megaterium were grown in lysogeny broth (LB) with tetracycline selection. After 12h, 1-2 L of terrific broth (TB) with tetracycline selection were inoculated with starter culture. Cultures were grown at 37°C, 180 RPM until the OD600 reached 0.8 or higher, upon which xylose (0.5% w/v final concentration) was added to induce expression of TcdB. Induced cultures were incubated overnight at 30°C, 180 RPM. Cells were pelleted at 4000 x g for 12 min and lysed in 20 mM Tris pH8, 0.1 M NaCl, 1 mg/ml lysozyme, 1% v/v protease inhibitor cocktail P8849 (MilliporeSigma), and 100 U/ml Pierce universal nuclease (88701, ThermoFisher). An EmulsiFlex C3 microfluidizer (Avestin) at 15,000 psi was then used to lyse the cells. Cell lysate was clarified by centrifugation at 14,000 × g, 4°C for 20 min and supernatants were filtered with a 0.2 $\mu$M filter. Next, affinity chromatography with a 5 ml HisTrap FF Ni-NTA column was used to purify the toxin on an Äkta FPLC systems (Cytiva). Proteins were eluted with 500 mM imidazole and a single peak was collected, and then further purified on a 1 ml HiTrap Q column (Cytiva). Proteins were eluted using a gradient of buffer with 100 mM to 1 M NaCl. Purified protein flash frozen in liquid nitrogen with 10% glycerol and stored at -80°C. For biotinylation, TcdB was incubated with a 15-fold molar excess of maleimide biotin (ThermoFisher) overnight at 4°C in 20mM Tris pH7.5, 150mM NaCl, 5% glycerol. The biotinylated TcdB was then purified on a 1 ml HiTrap Q column (Cytiva). Alternatively, TcdB 1285-1804 with C-terminal Avitag (RBD toxin fragment) was generated, purified and biotinylated in the same manner as the full length protein.

#### 9.4 Surface Plasmon Resonance (SPR) Binding Experiments

For initial binding screens, SARS-CoV-2 receptor binding domain and influenza hemagglutinin were purchased commercially from Sino Biological (40592-V08B-B, 11055-V08B-B respectively). RSV F protein was also purchased commercially from Acro Bio (RSF-V82E7). Biotinylated TcsL

(Sema6A binding fragment) and TcdB (Frizzled-7 binding fragment) were a gift from Dr. Roman Melynk at the Hospital for Sick Children (Toronto, Canada).

All SPR experiments used the same buffer prepared for SEC (HBS-EP). Target protein was captured using biotin CAPture kit (Cytiva) with a target Rmax of the analyte of 50 - 150 response units. 6-step, 5 fold dilutions starting at 5$\mu$M of each nanobody were tested in single cycle kinetics with 120 s association and 300 s dissociation at a flow rate of 30$\mu$l/min. The chip was regenerated after each cycle using the manufacturer's recommended regeneration buffer (1:3 1 M NaOH and 8M guanidinium HCl). For TcdB where target protein was limited, all nanobodies were first screened at a single concentration and the chip was regenerated by allowing the nanobodies to fully dissociate (given the short half life of these designs, the signal returned to baseline in $<$ 1 minute). Follow up titrations for the most promising hits, the nanobodies H2 and B9 used 6-step single cycle kinetics but with a 2-fold dilution and an upper concentration of 2$\mu$M. Both steady state affinity and global kinetic fitting using langmuir 1:1 interaction was used to determine the $Kd$ and generally agreed with each other. The values reported here are from kinetic fits.

For scFv experiments, scFvs were captured on a CM5 chip through amin conjugation. Single cycle kinetics were measured in HBS-EP across 6- or 8-fold dilution series using the same conditions for association and dissociation as above. Starting concentration and dilution steps were dependent on anticipated affinity and are specified in the relevant figure legends.

### 10 TcdB Nanobody

#### 10.1 Nanobody Evolution using OrthoRep

Yeast strain yAP196 (MATa AGA1:: pER-AGA1-HygR ura3-52::synTF-URA3 trp1 leu2delta1 his3delta200 pep4::HIS3 prb1delta1.6R can1 GAL met15; p1-MET15 landing pad; CENARS-Badboy3-Leu2) [43] was used for OrthoRep-driven affinity maturation [43, 44] of designed nanobodies. yAP196 strain expresses the orthogonal error-prone DNA polymerase, BadBoy357, encoded on a nuclear CEN/ARS plasmid. BadBoy3 replicates a cytoplasmic orthogonal p1 plasmid, also in the strain, at a high mutation rate of 10-4 substitutions per base while sparing genomic and nuclear DNA from hypermutation [45, 46]. yAP196's genome encodes AGA1 under the control of promoter pER, inducible by $\beta$-estradiol (Sigma Aldrich, E8875) through a synthetic transcription factor [43].

To encode the designed nanobody TcdB-H2 on p1 for hypermutation and yeast display, the nanobody sequence TcdB-H2 was first cloned onto the p1 donor plasmid, pAP19131, using Golden Gate Assembly, resulting in plasmid pYY42. In this plasmid, the TcdB-H2 nanobody is encoded as an N-terminal fusion to an HA tag and the AGA2 gene. The TRP1 and NatMX selection markers are also present. This plasmid was then digested with the restriction enzyme, ScaI, or PCR amplified to generate a linear DNA product with homologous flanks for integration onto the p1 landing pad in yAP196 strain using a yeast transformation method previously described58.

A colony from the transformation plate (SC without histidine (H), leucine (L), uracil (U), tryptophan (W), methionine (M), and cysteine (C) and with nourseothricin (Nat)) was picked into 3 ml of liquid media (SC-HLUW) and grown to semi-saturation at 30°C with shaking at 200 rpm. The culture was then diluted to an OD600 of 0.05 in 10 ml of induction media (SC-HLUW plus 200nM of $\beta$-estradiol) and grown for 16-24h or diluted to an OD600 of 0.5 in 10ml of induction media and grown for 2-4h at 30°C with shaking at 200rpm. Approximately $5 \times 10^7$ induced yeast cells were harvested from the induction culture and washed twice with HBSBM buffer (20 mM Tris-HCl pH 7.5, 100mM NaCl, 0.1% BSA, and 5mM maltose). The cells were subsequently stained in 250$\mu$L of primary staining solution (HBSBM buffer with biotinylated TcdB) at 4°C for 1 hour with rotation. After primary staining, the cells were washed with HBSBM buffer and stained with 250$\mu$l of secondary staining solution (HBSBM buffer with 0.5$\mu$l of mg/ml streptavidin-AF647 (ThermoFisher, S32357) and 2.5$\mu$l of 0.1 mg/ml anti-HA-AF488 (R&D system, IC6875G)) for 20 minutes at 4°C with rotation. Next, 1$\mu$l of 1mg/ml propidium iodide (Sigma Aldrich, 81845) was added and incubated for 1min on ice to assess cell viability. The cells were then washed twice with HBSBM buffer and resuspended in 4ml of HBSBM buffer for sorting on a Sony SH800 cell sorter. The first gate was set to separate live cells (PI negative) from dead cells (PI positive). Next, live cells were gated based on forward scatter area (FSC-A) and side scatter area (SSC-A) to exclude clumps of cells and debris artifacts. To ensure singlet selection, cells with appropriate sizes were further gated based on FSC-A and forward scatter height (FSC-H). Finally, a two-parameter density plot was used, with the y-axis representing the AF488 signal and the x-axis representing the AF647 signal, to sort high-affinity binders. A diagonal gate was applied based on the display level measured via anti-HA-AF488, while TcdB binding was assessed using streptavidin-AF647. The diagonal gate was positioned to sort cells with the best binding relative to high display levels, as shown in the figure (Extended Data Fig. 10A). 300 – 500 cells from the gated population were sorted into 3ml of SC-HLUW media. Sorted cells were grown at 30°C with shaking at 200rpm, during which the p1 plasmid encoding the nanobody autonomously underwent hypermutation through its replication by BadBoy3. When semi-saturation was reached (approximately 2-3 days), the process of induction, staining, sorting, and growth was repeated, constituting the next cycle of evolution. A total of 15 cycles of evolution were carried out. The concentration of biotinylated TcdB was reduced from cycle to cycle, progressively challenging the evolving nanobody to bind with higher and higher affinity.

The population of evolved nanobody sequences from the 15th cycle was amplified by PCR following a yeast GC prep and cloned into a standard CEN/ARS yeast display plasmid, pYY22, that does not hypermutate. This plasmid population was transformed into yAP174 (MATa AGA1::pER-AGA1-HygR ura3-52::synTF-URA3 trp1 leu2delta1 his3delta200 pep4::HIS3 prb1delta1.6R can1 GAL) [43] and the resulting population was twice sorted using labeling by 3nM of biotinylated TcdB to isolate the best clones. The sorted population was plated, and random colonies were picked and sequenced, leading to the identification of individual clones that achieved high-affinity binding of TcdB. See Table 8 for the full sequences of the plasmids used in this study.

| Plasmid Name | Description | Sequence |
|---|---|---|
| pYY22 | CEN/ARS plasmid for β-estradiol inducible yeast surface display with a LEU2 auxotrophic selection marker for yeast and an AmpR marker for propagation in E. coli | tcgaggagaacttctagtatatctacatacctaatattattgccttattaaaaatggaatcccaacaattacatcaaaatccacattctcttcaaaatcaattgtcctgtactt<br>ccttgttcatgtgtgttcaaaaacgttatatttataggataattatactctatttctcaacaagtaattggttgtttggccgagcggtctaaggcgcctgattcaagaaatatctt<br>gaccgcagttaactgtgggaatactcaggtatcgtaagatgcaagagttcgaatctcttagcaaccattatttttttcctcaacataacgagaacacacaggggcgctat<br>cgcacagaatcaaattcgatgactggaaattttttgttaatttcagaggtcgcctgacgcatatacctttttcaactgaaaaattgggagaaaaaggaaaggtgagagc<br>gccggaaccggcttttcatatagaatagagaagcgttcatgactaaatgcttgcatcacaatacttgaagttgacaatattatttaaggacctattgtttttccaataggtg<br>gttagcaatcgtcttactttctaacttttcttaccttttacatttcagcaatatatatatatatatttcaaggatataccattctaatgtctgcccctaagaagatcgtcgttttgcca<br>ggtgaccacgttggtcaagaaatcacagccgaagccattaaggttcttaaagctatttctgatgttcgttccaatgtcaagttcgatttcgaaaatcatttaattggtggtgc<br>tgctatcgatgctacaggtgttccacttccagatgaggcgctggaagcctccaagaaggctgatgccgttttgttaggtgctgtgggtggtcctaaatggggaaccggta<br>gtgttagacctgaacaaggtttactaaaaatccgtaaagaacttcaattgtacgccaacttaagaccatgtaactttgcatccgactctcttttagacttatctccaatcaa<br>gccacaatttgctaaaggtactgacttcgttgttgtcagagaattagtgggaggtatttactttggtaagagaaaggaggacgatggtgatggtgtcgcttgggatagtga<br>acaatacaccgttccagaagtgcaaagaatcacaagaatggccgctttcatggccctacaacatgagccaccattgcctatttggtccttggataaagctaatgttttgg<br>cctcttcaagattatggagaaaaactgtggaggaaaccatcaagaacgaatttcctacattgaaggttcaacatcaattgattgattctgccgccatgatcctagttaag<br>aacccaacccacctaaatggtattataatcaccagcaacatgtttggtgatatcatctccgatgaagcctccgttatcccaggttccttgggtttgttgccatctgcgtccttg<br>gcctctttgccagacaagaacaccgcatttggtttgtacgaaccatgccacggttctgctccagatttgccaaagaataaggtcaaccctatcgccactatcttgtctgct<br>gcaatgatgttgaaattgtcattgaacttgcctgaagaaggtaaggccattgaagatgcagttaaaaaggttttggatgcaggtatcagaactggtgatttaggtggttcc<br>aacagtaccaccgaagtcggtgatgctgtcgccgaagaagttaagaaaatccttgcttaacctggaggacccttctctttagactattctactcttatgcacgtaaaaaa<br>ttctaggaaatatgtattaactaggagtaaaataaccggctagtggcattcatatagccgtctgtttacatctacatcacacatttcgagtgtatatctcgcaacgttggcgtt<br>aaataggcagatcacgtgctataaaaataattataatttaaattttttaatataaatatataaattaaaaatagaaagtaaaaaaagaaattaaagaaaaaatagtttttg<br>ttttccgaagatgtaaaagactctaggggatcgccaacaaatactaccttttatcttgctcttcctgctctcaggtattaatgccgaattgtttcatcttgtctgtgtagaaaa<br>ccacacacgaaaatcctgtgattttacattttacttatcgttaatcgaatgtatatctatttaatctgcttttcttgtctaataaatatatatgtaaagtacgctttttgttgaaattttttt<br>aaacctttgtttattttttttttcttcattccgtaactcttctaccttctttatttactttctaaaatccaaatacaaaacataaaaataaataaacacagagtaaattcccaaattatt<br>ccatcattaaaagatacgaggcgcgtgtaagttacaggcaagcgatccgtcccgagcggccgcgattatcaaaaaggatcttcacctagatccttttaaattaaaaat<br>gaagttttaaatcaatctaaagtatatatgagtaaacttggtctgacagttaccaatgcttaatcagtgaggcacctatctcagcgatctgtctatttcgttcatccatagttgc<br>ctgactccccgtcgtgtagataactacgatacgggagggcttaccatctggccccagtgctgcaatgataccgcgggacccacgctcaccggctccagatttatcag<br>caataaaccagccagccggaagggccgagcgcagaagtggtcctgcaactttatccgcctccatccagtctattaattgttgccgggaagctagagtaagtagttcg<br>ccagttaatagtttgcgcaacgttgttgccattgctacaggcatcgtggtgtcacgctcgtcgtttggtatggcttcattcagctccggttcccaacgatcaaggcgagttac<br>atgatcccccatgttgtgcaaaaaagcggttagctccttcggtcctccgatcgttgtcagaagtaagttggccgcagtgttatcactcatggttatggcagcactgcataat<br>tctcttactgtcatgccatccgtaagatgcttttctgtgactggtgagtactcaaccaagtcattctgagaatagtgtatgcggcgaccgagttgctcttgcccggcgtcaat<br>acgggataataccgcgccacatagcagaactttaaaagtgctcatcattggaaaacgttcttcggggcgaaaactctcaaggatcttaccgctgttgagatccagttc<br>gatgtaacccactcgtgcacccaactgatcttcagcatcttttactttcaccagcgtttctgggtgagcaaaaacaggaaggcaaaatgccgcaaaaaagggaataa<br>gggcgacacggaaatgttgaatactcatactcttcctttttcaatattattgaagcatttatcagggttattgtctcatgagcggatacatatttgaatgtatttagaaaaataa<br>acaaataggggttccgcgcacatttccccgaaaagtgccacctgtcatgaccaaaatcccttaacgtgagttttcgttccactgagcgtcagaccccgtagaaaagat<br>caaaggatcttcttgagatcctttttttctgcgcgtaatctgctgcttgcaaacaaaaaaaccaccgctaccagcggtggtttgtttgccggatcaagagctaccaactcttt<br>ttccgaaggtaactggcttcagcagagcgcagataccaaatactgttcttctagtgtagccgtagttaggccaccacttcaagaactctgtagcaccgcctacatacctc<br>gctctgctaatcctgttaccagtggctgctgccagtggcgataagtcgtgtcttaccgggttggactcaagacgatagttaccggataaggcgcagcggtcgggctga<br>acggggggttcgtgcacacagcccagcttggagcgaacgacctacaccgaactgagatacctacagcgtgagctatgagaaagcgccacgcttcccgaaggga<br>gaaaggcggacaggtatccggtaagcggcagggtcggaacaggagagcgcacgagggagcttccagggggaaacgcctggtatctttatagtcctgtcgggtttc<br>gccacctctgacttgagcgtcgatttttgtgatgctcgtcaggggggcggagcctatggaaaaacgccagcaacgcggcctttttacggttcctggccttttgctggcctttt<br>tgctcacatgttctttcctgcgttatcccctgattctgtggataaccgtgcggccgcccctgaattcgcatctagactgatgagacgtggtagagccacaaacagccggta<br>caagcaacgatctccaggaccatctgaatcatgcgcggatgacacgaactcacgacggcgatcacagacattaacccacagtacagacactgcgacaacgtgg<br>caattcgtcgcaatacaacgCCTAGGTTGAGGCCAACGaTTATGGGAGatcgaatattaataTTATGGGAGatcgaatattaataTTATGG<br>GAGatcgaatattaataTTATGGGAGatcgaatattaataTTATGGGAGatcgaatattaataTTATGGGAGatcgaatattaataTTATGGGA<br>GaTGGCATGAATGGCCTTAAGcagatccgccaggcgtgtatatatagcgtggatggccaggcaactttagtgctgacacatacaggcatatatatatgtg<br>tgcgacgacacatgatcatatggcatgcatgtgctctgtatgtatataaaactcttgttttcttcttttctctaaatattctttccttatacattaggacctttgcagcataaattact<br>atacttctatagacacacaaacacaaatacacacactaaATGAGATTCCCATCTATCTTCACCGCTGTTTTGTTCGCTGCTTCTTCT<br>GCTTTGGCTGCTCCAGCTAACACCACCACCGAAGACGAAACCGCTCAAATCCCAGCTGAAGCTGTTATCGACTAC<br>TCTGACTTGGAAGGTGACTTCGACGCTGCTGCTTTGCCATTGTCTAACTCTACCAACAACGGTTTGTCTTCTACCA<br>ACACCACCATCGCTTCTATCGCTGCTAAGGAAGAAGGTGTTCAATTGGACAAGAGAGAAGCTGACGCAggtaccgag<br>accgaaagtgaaacgtgatttcatgcgtcattttgaacattttgtaaatcttatttaataatgtgtgcggcaattcacatttaatttatgaatgttttcttaacatcgcggcaactc<br>aagaaacggcaggttcggatcttagctactagagaaagaggagaaatactagatgcgtaaaggcgaagagctgttcactggtgtcgtccctattctggtggaactgg<br>atggtgatgtcaacggtcataagttttccgtgcgtggcgagggtgaaggtgacgcaactaatggtaaactgacgctgaagttcatctgtactactggtaaactgccggtt<br>ccttggccgactctggtaacgacgctgacttatggtgttcagtgctttgctcgttatccggaccatatgaagcagcatgacttcttcaagtccgccatgccggaaggctat<br>gtgcaggaacgcacgatttcctttaaggatgacggcacgtacaaaacgcgtgcggaagtgaaatttgaaggcgataccctggtaaaccgcattgagctgaaaggc<br>attgactttaaagaggacggcaatatcctgggccataagctggaatacaattttaacagccacaatgtttacatcaccgccgataaacaaaaaaatggcattaaagc<br>gaattttaaaattcgccacaacgtggaggatggcagcgtgcagctggctgatcactaccagcaaaacactccaatcggtgatggtcctgttctgctgccagacaatca<br>ctatctgagcacgcaaagcgttctgtctaaagatccgaacgagaaacgcgatcatatggttctgctggagttcgtaaccgcagcgggcatcacgcatggtatggatg<br>aactgtacaaatgaccaggcatcaaataaaacgaaaggctcagtcgaaagactgggcctttcgttttatctgttgtttgtcggtgaacgctctctactagagtcacactg<br>gctcaccttcgggtgggcctttctgcgtttataggtctcgtcgacGGTTCCaaggacaatagctcgacgattgaaggtagatacccatacgacgttccagactacg<br>ctctgcaggctagtggtggtggtggttctggtggtggtggttctggtggtggtggttctgctagcgacgtcCAGGAACTGACAACTATATGCGAGCAAA<br>TCCCCTCACCAACTTTAGAATCGACGCCGTACTCTTTGTCAACGACTACTATTTTGGCCAACGGGAAGGCAATGC<br>AAGGAGTTTTTGAATATTACAAATCAGTAACGTTTGTCAGTAATTGCGGTTCTCACCCCTCAACAACTAGCAAAGG<br>CAGCCCCATAAACACACAGTATGTTTTTTGAtagctcgagatctgataacaacagtgtagatgtaacaaaatcgactttgttcccactgtacttttag<br>ctcgtacaaaatacaatatacttttcatttctccgtaaacaacatgttttcccatgtaatatccttttctatttttcgttccgttaccaactttacacatactttatatagctattcactt<br>ctatacactaaaaaactaagacaattttaattttgctgcctgccatatttcaatttgttataaattcctataatttatcctattagtagctaaaaaaagatgaatgtgaatcgaat<br>cctaagagaatt |
| pYY42 | Plasmid for generation of a linear | gagatacctacagcgtgagctatgagaaagcgccacgcttcccgaagggagaaaggcggacaggtatccggtaagcggcagggtcggaacaggagagcgca<br>cgagggagcttccagggggaaacgcctggtatctttatagtcctgtcgggtttcgccacctctgacttgagcgtcgatttttgtgatgctcgtcaggggggcggagcctat<br>ggaaaaacgccagcaacgcggcctttttacggttcctggccttttgctggccttttgctcacatgttctttcctgcgttatcccctgattctgtggataaccgtattaccgcctttt<br>gagtgagctgataccgctcgccgcagccgaacgaccgagcgcagcgagtcagtgagcgaggaagcggaagagcgcccaatacgcaaaccgcctctccccgc<br>gcgttggccgattcattaatgcagctggcacgacaggtttcccgactggaaagcgggcagtgagcgcaacgcaatagtactataatatatgaattacattattaattta<br>aaaacgcgtgatgacctatacataggaagatctatagaaacaaaaagattaataactttcaaatatcagaaaaatatagaaacatgtgataagctcatagacatAta |

| | | |
|---|---|---|
| | integration cassette for integration onto p1 where the cassette encodes a VHH_TcdB_H2 nanobody-HAtag-AGA2 fusion with TRP1 and NatMX selection markers. The integration cassette is generated by linearization with ScaI or by PCR. This plasmid contains an AmpR marker for propagation in E. coli | aaggatccATGAGATTCCCATCTATCTTCACCGCTGTTTTGTTCGCTGCTTCTTCTGCTTTGGCTGCTCCAGCTAACACCACCACCGAAGACGAAACCGCTCAAATCCCAGCTGAAGCTGTTATCGACTACTCTGACTTGGAAGGTGACTTCGACGCTGCTGCTTTGCCATTGTCTAACTCTACCAACAACGGTTTGTCTTCTACCAACACCACCATCGCTTCTATCGCTGCTAAGGAAGAAGGTGTTCAATTGGACAAGAGAGAAGCTGACGCAggtaccCAGGTTCAGTTGGTAGAAAGCGGTGGTGGGCTTGTTCAACCTGGAGGATCTCTGCGTCTTAGTTGCGCCGCCAGCGGTGAAAGCCTGGCGAATATGATCTCATTAGGGTGGTTTAGGCAGGCACCAGGGCAAGGGCTTGAGGCGGTGGCGGCGATCTCTGGGAACGGGAAGTTCACATATTATGCGGATTCAGTAAAAGGTAGATTTACCATTAGTAGGGATAATTCCAAAAATACGTTGTATCTACAAATGAACTCCTTACGTGCAGAAGATACTGCTGTGTACTACTGCGCCGCGTGGGGCAGGAAGTTATACTGGGGACAAGGCACACTTGTCACTGTTTCAgtcgacGGTTCCaaggacaatagctcgacgattgaaggtagatacccatacgacgttccagactacgctctgcaggctagtggtggtggtggttctggtggtggtggttctggtggtggtggttctgctagcgacgtcCAGGAACTGACAACTATATGCGAGCAAATCCCCTCACCAACTTTAGAATCGACGCCGTACTCTTTGTCAACGACTACTATTTTGGCCAACGGGAAGGCAATGCAAGGAGTTTTTGAATATTACAAATCAGTAACGTTTGTCAGTAATTGCGGTTCTCACCCCTCAACAACTAGCAAAGGCAGCCCCATAAACACACAGTATGTTTTTTGAAGAAGACTTGtcatgaAAAAAAAAAAAAAAAAAAAAAAAAAAAAAAAAAAAAAAAAAAAAAAAAAAAAAAAAAAAAAAAAAAAAAAAAAAACCTGTCACCGGATGTGTTTTCCGGTCTGATGAGTCCGTGAGGACGAAACAGGgaattcgcgaatttcttatgatttatgatttttattattaaataagttataaaaaaaataagtgtatacaaattttaaagtgactcttaggttttaaaacgaaaattcttattcttgagtaactctttcctgtaggtcaggttgctttctcaggtatagcatgaggtcgctcgagctcttaagggcaaggcatagacatatacaaggcctgttcaccgtcagatgctgttccatcatataaagcagtatctaaaccacataatgtgaaacccattcttctataagcatggatagcaggggcgttaacattagtaacttctaaccataaatgaccggctcctctctctcttgcaaattctgtagccaaacccatcaaagctcttcctacaccatgacctctatgctctggggcgacttctatatcttcaacggtcaaccttctattccaacctgaataagaaacgactacaaaaccagccaaatcaccgtcatcaccgtatgctacgaaagttcttgaatctgggtcaccatcttctccgtcgtcgctttcgtcatcagattcatcatctggaaagactttagttaatggaggatcgactgggacttctcttaaggtaaatccatcaccggtggcggttactctaaagacggtgtcggtggtgaatgaaccgtctaaagcctcaattgcttcagcatcaccagggactgatgttctgtatctgtaggctgtatcatctaaggtagtacccattttataattatataaaaactttcatatagagctccgtttctattatgaatttcatttataaagtttatgtacaaatatcataaaaaaagagaatcttttttcttagcatttttgacgaaatttgctattttgttagagtcttttacaccatttgtctccacacctccgcttacatcaacaccaataacgccatttaatctaagcgcatcaccaacattttctggcgtcagtccaccagctaacataaaatgtaagctttcggggctctcttgccttccaacccagtcagaaatcgagttccaatccaaaagttcacctgtcccacctgcttctgaatcaaacaagggaataaacgaatgaggtttctgtgaagctgcactgagtagtatgttgcagtcttttggaaatacgagtcttttaataactggcaaaccgaggaactcttggtattcttgccacgactcatctccatgcagttggacgatatcaatgccgtaatcattgaccagagccaaaacatcctccttaggttgattacgaaacacgccaaccaagtatttcggagtgcctgaactatttttatatgcttttacaagacttgaaattttccttgcaataaccgggtcaattgttctctttctattgggcacacatataatacccagcaagtcagcatcggaatctagagcacattctgcggcctctgtgctctgcaagccgcaaactttcaccaatggaccagaactacctgtgaaattaataacagacattttataattattataaaaactttcatataagagatataaaatttaatatggaaaaaataagacaagaaagatacaaccaaatgaaagaagctctaaatagtgttgaaggttataaaggaaaaattgtagcctcagactcagattggtgtttcaaagatcctCAAGGCAATAGAATAACAGATTTTGATAGTactttaagccagccccgacacccgccaacacccgctgacgcgccctgacgggcttgtctgctcccggcatccgcttacagacaagctgtgaccgtctccgggagctgcatgtgtcagaggttttcaccgtcatcaccgaaacgcgcgagacgaaaggcaaaaaaaaaggctccaaaaggagcctttaattgtatcggtttaccaatgcttaatcagtgaagcccctatttcagcaatctgtctatttcgttcgtccatagttgcctgactacccgtcgtgtagataactacgatacgggaaggcttaccatctggtccaagtgcggcaatgataccccgtgaaccacgctctcctgccccagatttatcagcaataaaccagccagcaggaagtgccgaacgcagtagtggtcctgcaactttatccgcttcaagccagtctattaattgttgccgtgaagccagagtaagtagttcgccagttaatagtttgcgtaatgttgttgccattgccgcaggcattgtggtgtctcgctcgtcgtttggtatggcttcattcagttctggttcccaacggtcaaggcgagttacatggtctcccatattgtgcaaaaaagcggttaattccttcggtccaccgatggttgtcagaagtaagttcgccgcagtgttatcactcatcgttatggcggcactacataattctcttaccgtcattccgtccgtaaggtgcttttctgtcactggggagtattcaaccaagtcattctgcgaatagtgtattcgtcgtccgagttgctcttgccccgcatctacacgggataatactgctccacatagcagaactttgaaagtgctcatcattggaaaacgctcttcagggcgaaaactctcaaggattttaccgctattcaagtccagttctatgtatcccactcgtgcccccaattggtcctctgcgtcttttactttcacgagcgtttctgggtgagcaaaaacaggaaggcaaaatgccgcaaaaaagggaataagggcgacacggaaatgttgaatactcatactcttcctttttcaatattattgaagcatttatcagggttattgtctgatgagcggatacatatttgaatgtatttagaaaaataaacaaataggggaattaaaaaaaagcccgctcattaggcgggctactagtcggatatagttcctcctttcagcaaaaaacccctcaagacccgtttagaggccccaaggggttatgctagttattgctcagcggtggcagcagccaactcagcttcctttcgggctttgttagcagccggatcttctagaagaccccgtagaaaagatcaaaggatcttcttgagatcctttttttctgcgcgtaatctgctgcttgcaaacaaaaaaaccaccgctaccagcggtggtttgtttgccggatcaagagctaccaactctttttccgaaggtaactggcttcagcagagcgcagataccaaatactgttcttctagtgtagccgtagttaggccaccacttcaagaactctgtagcaccgcctacatacctcgctctgctaatcctgttaccagtggctgctgccagtggcgataagtcgtgtcttaccgggttggactcaagacgatagttaccggataaggcgcagcggtcgggctgaacggggggttcgtgcacacagcccagcttggagcgaacgacctacaccgaact |
| pAP191 | Plasmid for generation of a linear integration cassette for integration onto p1 where the cassette encodes GOI-HA-Aga2 fusion with | actacgatacgggaaggcttaccatctggtccaagtgcggcaatgataccccgtgaaccacgctctcctgccccagatttatcagcaataaaccagccagcaggaagtgccgaacgcagtagtggtcctgcaactttatccgcttcaagccagtctattaattgttgccgtgaagccagagtaagtagttcgccagttaatagtttgcgtaatgttgttgccattgccgcaggcattgtggtgtctcgctcgtcgtttggtatggcttcattcagttctggttcccaacggtcaaggcgagttacatggtcAcccatattgtgcaaaaaagcggttaattccttcggtccaccgatggttgtcagaagtaagttcgccgcagtgttatcactcatcgttatggcggcactacataattctcttaccgtcattccgtccgtaaggtgcttttctgtcactggggagtattcaaccaagtcattctgcgaatagtgtattcgtcgtccgagttgctcttgccccgcatctacacgggataatactgctccacatagcagaactttgaaagtgctcatcattggaaaacgctcttcagggcgaaaactctcaaggattttaccgctattcaagtccagttctatgtatcccactcgtgcccccaattggtcctctgcgtcttttactttcacgagcgtttctgggtgagcaaaaacaggaaggcaaaatgccgcaaaaaagggaataagggcgacacggaaatgttgaatactcatactcttcctttttcaatattattgaagcatttatcagggttattgtctgatgagcggatacatatttgaatgtatttagaaaaataaacaaataggggaattaaaaaaaagcccgctcattaggcgggctactagtcggatatagttcctcctttcagcaaaaaacccctcaagacccgtttagaggccccaaggggttatgctagttattgctcagcggtggcagcagccaactcagcttcctttcgggctttgttagcagccggatcttctagaagaccccgtagaaaagatcaaaggatcttcttgagatcctttttttctgcgcgtaatctgctgcttgcaaacaaaaaaaccaccgctaccagcggtggtttgtttgccggatcaagagctaccaactctttttccgaaggtaactggcttcagcagagcgcagataccaaatactgttcttctagtgtagccgtagttaggccaccacttcaagaactctgtagcaccgcctacatacctcgctctgctaatcctgttaccagtggctgctgccagtggcgataagtcgtgtcttaccgggttggactcaagacgatagttaccggataaggcgcagcggtcgggctgaacggggggttcgtgcacacagcccagcttggagcgaacgacctacaccgaactgagatacctacagcgtgagctatgagaaagcgccacgcttcccgaagggagaaaggcggacaggtatccggtaagcggcagggtcggaacaggagagcgcacgagggagcttccagggggaaacgcctggtatctttatagtcctgtcgggtttcgccacctctgacttgagcgtcgatttttgtgatgctcgtcaggggggcggagcctatggaaaaacgccagcaacgcggcctttttacggttcctggccttttgctggccttttgctcacatgttctttcctgcgttatcccctgattctgtggataaccgtattaccgcctttgagtgagctgataccgctcgccgcagccgaacgaccgagcgcagcgagtcagtgagcgaggaagcggaagagcgcccaatacgcaaaccgcctctccccgcgcgttggccgattcattaatgcagctggcacgacaggtttcccgactggaaagcgggcagtgagcgcaacgcaatagtactataatatatgaattacattattaatttaaaaacTcgtgatgacctatacataggaagatctatagaaacaaaaagattaataactttcaaatatcagaaaaatatagaaacatgtgataagctcatagacatAtaaaaaTATGAGATTCCCATCTATCTTCACCGCTGTTTTGTTCGCTGCTTCTTCTGCTTTGGCTGCTCCAGCTAACACCACCACCGAAGACGAAACCGCTCAAATCCCAGCTGAAGCTGTTATCGACTACTCTGACTTGGAAGGTGACTTCGACGCTGCTGCTTTGCCATTGTCTAACTCTACCAACAACGGTTTGTCTTCTACCAACACCACCATCGCTTCTATCGCTGCTAAGGAAGAAGGTGTTCAATTGGACAAGAGAGAAGCTGACGCAgagaccgaaagtgaaacgtgatttcatg |

| | | |
|---|---|---|
| | TRP1 and NatMX selection markers. The integration cassette is generated by linearization with ScaI or by PCR. This plasmid contains an AmpR marker for propagation in E. coli | cgtcattttgaacattttgtaaatcttatttaataatgtgtgcggcaattcacatttaatttatgaatgttttcttaacatcgcggcaactcaagaaacggcaggttcggatctta<br>gctactagagaaagaggagaaatactagatgcgtaaaggcgaagagctgttcactggtgtcgtccctattctggtggaactggatggtgatgtcaacggtcataagttt<br>tccgtgcgtggcgagggtgaaggtgacgcaactaatggtaaactgacgctgaagttcatctgtactactggtaaactgccggttccttggccgactctggtaacgacg<br>ctgacttatggtgttcagtgctttgctcgttatccggaccatatgaagcagcatgacttcttcaagtccgccatgccggaaggctatgtgcaggaacgcacgatttccttta<br>aggatgacggcacgtacaaaacgcgtgcggaagtgaaatttgaaggcgataccctggtaaaccgcattgagctgaaaggcattgactttaaagaggacggcaat<br>atcctgggccataagctggaatacaattttaacagccacaatgtttacatcaccgccgataaacaaaaaaatggcattaaagcgaattttaaaattcgccacaacgtg<br>gaggatggcagcgtgcagctggctgatcactaccagcaaaacactccaatcggtgatggtcctgttctgctgccagacaatcactatctgagcacgcaaagcgttct<br>gtctaaagatccgaacgagaaacgcgatcatatggttctgctggagttcgtaaccgcagcgggcatcacgcatggtatggatgaactgtacaaatgaccaggcatc<br>aaataaaacgaaaggctcagtcgaaagactgggcctttcgttttatctgttgtttgtcggtgaacgctctctactagagtcacactggctcaccttcgggtgggcctttctgc<br>gtttataggtctcGGTTCCaaggacaatagctcgacgattgaaggtagatacccatacgacgttccagactacgctctgcaggctagtggtggtggtggttctggtg<br>gtggtggttctggtggtggtggttctgctagcgacgtcCAGGAACTGACAACTATATGCGAGCAAATCCCCTCACCAACTTTAGAATCG<br>ACGCCGTACTCTTTGTCAACGACTACTATTTTGGCCAACGGGAAGGCAATGCAAGGAGTTTTTGAATATTACAAAT<br>CAGTAACGTTTGTCAGTAATTGCGGTTCTCACCCCTCAACAACTAGCAAAGGCAGCCCCATAAACACACAGTATGT<br>TTTTTGAatcctaataatGGCCGCAAAAAAAAAAAAAAAAAAAAAAAAAAAAAAAAAAAAAAAAAAAAAAAAAAAA<br>AAAAAAAAAAAAAAAAAAACCTGTCACCGGATGTGTTTTCCGGTCTGATGAGTCCGTGAGGACGAAACaggactagG<br>CTgcgaatttcttatgatttatgatttttattattaaataagttataaaaaaaataagtgtatacaaattttaaagtgactcttaggttttaaaacgaaaattcttattcttgagta<br>actctttcctgtaggtcaggttgctttctcaggtatagcatgaggtcgctcgagctcttaagggcaaggcatagacatatacaaggcctgttcaccgtcagatgctgttcca<br>tcatataaagcagtatctaaaccacataatgtgaaacccattcttctataagcatggatagcaggggcgttaacattagtaacttctaaccataaatgaccggctcctct<br>ctctcttgcaaattctgtagccaaacccatcaaagctcttcctacaccatgacctctatgctctggggcgacttctatatcttcaacggtcaaccttctattccaacctgaata<br>agaaacgactacaaaaccagccaaatcaccgtcatcaccgtatgctacgaaagttcttgaatctgggtcaccatcttctccgtcgtcgctttcgtcatcagattcatcatc<br>tggaaagactttagttaatggaggatcgactgggacttctcttaaggtaaatccatcaccggtggcggttactctaaagacggtgtcggtggtgaatgaaccgtctaaa<br>gcctcaattgcttcagcatcaccagggactgatgttctgtatctgtaggctgtatcatctaaggtagtacccattttataattattataaaaactttcatatagagctccgtttct<br>attatgaatttcatttataaagtttatgtacaaatatcataaaaaaagagaatcttttTCAtttcttagcatttttgacgaaatttgctattttgttagagtcttttacaccatttgtct<br>ccacacctccgcttacatcaacaccaataacgccatttaatctaagcgcatcaccaacattttctggcgtcagtccaccagctaacataaaatgtaagctttcggggct<br>ctcttgccttccaacccagtcagaaatcgagttccaatccaaaagttcacctgtcccacctgcttctgaatcaaacaagggaataaacgaatgaggtttctgtgaagct<br>gcactgagtagtatgttgcagtcttttggaaatacgagtcttttaataactggcaaaccgaggaactcttggtattcttgccacgactcatctccatgcagttggacgatatc<br>aatgccgtaatcattgaccagagccaaaacatcctccttaggttgattacgaaacacgccaaccaagtatttcggagtgcctgaactatttttatatgcttttacaagactt<br>gaaattttccttgcaataaccgggtcaattgttctctttctattgggcacacatataatacccagcaagtcagcatcggaatctagagcacattctgcggcctctgtgctctg<br>caagccgcaaactttcaccaatggaccagaactacctgtgaaattaataacagacattttataattattataaaaactttcatataagagatataaaatttaatatggaa<br>Gaaataagacaagaaagatacaaccaaatgaaagaagctctaaatagtgttgaaggttataaaggaaaaattgtagcctcagactcagattggtgtttcaaagatc<br>ctCAAGGCAATAGAATAACAGATTTTGATAGTactttaagccagccccgacacccgccaacacccgctgacgcgccctgacgggcttgtctgctcc<br>cggcatccgcttacagacaagctgtgaccgtctccgggagctgcatgtgtcagaggttttcaccgtcatcaccgaaacgcgcgagacgaaaggcaaaaaaaagg<br>ctccaaaaggagcctttaattgtatcggtttaccaatgcttaatcagtgaagcccctatttcagcaatctgtctatttcgttcgtccatagttgcctgactacccgtcgtgtaga<br>ta |

Supplementary Methods Table 8: Orthorep plasmids used in this study .

##### 10.2 TcdB Neutralization

Vero CSPG4-knockout cells [47] were grown in DMEM with 10% fetal bovine serum in the presence of penicillin and streptomycin (complete DMEM), at 37°C, 5% $CO_2$, and seeded in 96 well CellBIND plates (Corning) at a density of 5000 cells/well. After 24 hours, a Bravo liquid handler (Agilent) was used to add serial diluted test samples to cells, immediately followed by TcdB (ribotype VP10463) to a final concentration of 3 pM. Viability was assayed by adding alamarBlue Cell Viability Reagent (aka Resazurin, Thermofisher) after microscopic examination indicated cell rounding in the vehicle control wells (up to 48 hours post TcdB addition), and reading fluorescence signal (ex:555; em:585) 3 hours later in a Spectramax m5e plate reader (Molecular Devices). For visualization of cell rounding, Vero CSPG4 KO cells were seeded at a density of 4000 cells/well and used 24 hours post plating. Cells were preloaded with 1$\mu$M Celltracker Orange (Thermo) for 1 hour before changing media back to cDMEM. A Bravo liquid handler was used to add serial diluted test samples to cells, immediately followed by TcdB (ribotype VP10463) to a final concentration of 1.5 pM. The cell plates were returned to the incubator for 24 hours before imaging on a Cellomics Array Scan (Thermo) instrument, using a 10X objective and a sample rate of 100 objects per well.

#### 11 PHOX2B scFv

##### 11.1 Isothermal titration calorimetry (ITC)

ITC experiment between Phox2b:HLA-C*07:02 and herceptin_VLVH-His-Avi binder was obtained using a MicroCal VP-ITC system (Malvern Panalytical). All proteins were exhaustively dialyzed into buffer [150mM NaCl and 20mM sodium phosphate (pH 7.2)] overnight and filtered through a 0.22$\mu$m polyethersulfone (PES) membrane. A syringe containing pHLA-C*07:02 at 30

$\mu$M with 1mM peptide (QYNPIRTTF) was titrated into the calorimetry cell containing 2$\mu$M herceptin_VLVH-His-Avi and the same peptide (1mM). Injection volumes of 10$\mu$l were performed for a duration of 10s and spaced 220s apart to allow a complete return to baseline. Data were processed and analyzed with Origin software. Isotherms were fit using a one-site ITC binding model. The first data point was excluded from the analysis. Reported $K_D$, $\Delta$H, and $\Delta$S were determined using a 1-site binding model.

#### 11.2 Neuroblastoma Samples and Cell Lines

Human-derived neuroblastoma cell lines, including SK-N-AS, NBSD, NB-1691, SK-N-FI, and SK-N-SH were obtained from the Maris Laboratory cell line bank. Neuroblastoma cell lines were cultured in Roswell Park Memorial Institute medium (RPMI 1640) supplemented with 10% FBS, 100U/ml penicillin, 100$\mu$g/ml streptomycin and 2mM L-glutamine. Other human cancer cell lines, including LS123, were obtained from the American Type Culture Collection (ATCC). LS123 cells were cultured in EMEM supplemented with 10% FBS, 100U/ml penicillin, 100$\mu$g/ml streptomycin and 2mM L-glutamine. The packaging cell line HEK293T was obtained from ATCC. HEK293T cells were cultured in either Dulbecco's Modified Eagle Medium (DMEM) supplemented with 10% FBS, 100U/ml penicillin, 100$\mu$g/ml streptomycin and 2mM L-glutamine or RPMI supplemented with 10% FBS, 100U/ml penicillin, 100$\mu$g/ml streptomycin and 2mM L-glutamine. All cell lines were grown under humidified conditions in 5% $CO_2$ at 37°C, and samples were regularly tested for mycoplasma contamination.

#### 11.3 Primary Human T Cells

Primary human T cells were obtained from anonymous donors through the Human Immunology Core at the University of Pennsylvania (Philadelphia, Pennsylvania) under a protocol approved by the Children's Hospital of Philadelphia Institutional Review Board. Cells were cultured using complete AIM-V (Thermo Fisher Scientific) supplemented with 10% FBS, 100U/ml penicillin, 100$\mu$g/ml streptomycin, 2mM L-glutamine, 5ng/ml recombinant IL-7 (PeproTech) and 5ng/ml recombinant IL-15 (PeproTech) under humidified conditions in 5% CO2 at 37°C. T cell donors provided informed consent through the University of Pennsylvania Immunology Core.

#### 11.4 CAR Design

scFvs designed against Phox2b 9mer in complex with HLA-C*07:02 were incorporated into second-generation CAR constructs containing 4-1BB and CD3$\zeta$ co-stimulatory domains and cloned into a pJoe lentiviral vector for expression and functional screening (Genscript, and ref [34]).

#### 11.5 Viral Production and Transduction of Primary T Cells

A second-generation lentiviral system was used to produce replication-deficient lentivirus. The day preceding transfection or the morning of transfection, HEK293T cells were plated in a 15cm dish. On the day of transfection, 98$\mu$l Lipofectamine 3000 (Life Technologies, Invitrogen) was added to 2ml room-temperature Opti-MEM medium (Gibco). Concurrently, 98$\mu$l P3000 reagent (Thermo Fisher Scientific), 20.8$\mu$g psPAX2 (encoding gag/pol), 10.92$\mu$g pMD2.6 (encoding VSV-G envelope) and a matching molar quantity of transfer plasmid were added to 2ml room-temperature Opti-MEM medium. The next morning Opti-MEM medium was replaced and virus supernatant was collected after 48h and passed through a 0.45$\mu$m syringe.

Fresh human primary T cells were activated in culture for 2 days in the presence of 5 ng/ml recombinant IL-7, 5 ng/ml recombinant IL-15 and anti-CD3/CD28 beads (Dynabeads, Human T-Activator CD3/CD28, Life Technologies) at a 1.6:1 bead:T cell ratio. On day 3, 1.6 ml retroviral supernatants were added to 24-well plates pre-treated with 500$\mu$l well/retronectin (50$\mu$g/ml, Takara) and spun at 2,000g for 2h at 32°C followed by spinoculation of activated T cells at $6 \times 10^5$ cells per well at 1000g for 20 min at 32°C. On day 4, cells were collected and washed, beads were magnetically removed, and cells were expanded in complete AIM-V.

#### 11.6 CAR T Cell Staining and Flow Cytometric Analysis

Surface expression of CAR-transduced primary T cells was measured by staining with PE-conjugated rabbit anti-(G4S)3 linker antibody (Cell Signaling) along with Alexa Fluor-conjugated anti-CD3, Spark Blue 574-conjugated anti-CD4 and Spark Blue 550-conjugated anti-CD8 antibodies (BioLegend). Cells were collected from culture, washed with 2ml 2% BSA in PBS (PBSA) at 300g for 5min, incubated with 1$\mu$l anti-(G4S)3 antibody and 2$\mu$l of anti-CD3, -CD4 and -CD8 antibodies for 20min in the dark, washed twice and resuspended in 300$\mu$l 2% BSA in PBS for analysis. Typically, $1 \times 10^6$ cells were used for staining. Flow cytometry data were collected using an Aurora Spectral Flow Cytometer (Cytek). The gating strategy for staining is shown in Extended Data Fig. 26G.

#### 11.7 Real-time Cytotoxicity Analysis using xCELLigence

Target tumor cells ($2 \times 10^4$ per well [SK-N-AS, NBSD, SK-N-FI, NB-1691, LS123] or $7.5 \times 10^4$ per well [SK-N-SH]) were seeded in xCELLigence E-Plate 96 (Agilent) and allowed to adhere and reach exponential growth phase during a 24-hour pre-incubation at 37°C in a humidified 5% CO2 atmosphere. Following establishment of stable baseline impedance measurements, transduced primary effector cells (CAR-T cells) were added at 15:1 total T cell effector-to-target (E:T) ratio (corresponding to $3 \times 10^5$ cells per well). Cell-mediated cytotoxicity was monitored in real-time by measuring electrical impedance every hour for 24-72 hours using the xCELLigence RTCA DP system. Cell Index (CI) values were automatically calculated using RTCA Software Pro 2.0 (Agilent) and normalized to the t=0 baseline measurement.

### Part III

### Electron Microscopy

#### 12 Electron Microscopy

##### 12.1 In-House Production of Iowa43 HA

Natively glycosylated, trimeric influenza hemagglutinin (strain A/USA:Iowa/1943 H1N1) was produced via transient transfection of Expi293F cells (Gibco) and purified by immobilized metal ion affinity chromatography. A gene encoding Iowa43 HA, codon optimized for *Homo sapiens*, was cloned into the CMVR mammalian expression vector and transformed into NEB5$\alpha$ competent cells (New England Biolabs). Transfection-grade, endotoxin-free plasmid was prepared from the Nucleobond Xtra Maxi EF system (Machery-Nagel). Expi293F cultures were routinely maintained in Expi93F Expression Medium (Gibco) in an incubated shaker at 37°C, 70% humidity, 8% CO2 with 125 rpm oscillation. On the day of transfection, cultures in the logarithmic growth phase were diluted to 3 million cells per ml. Transfection was performed with a ratio of 1$\mu$g of plasmid DNA and 3$\mu$g of PEI-MAX (Polysciences) per ml of culture; the DNA and PEI-MAX was complexed in Opti-MEM Reduced Serum Medium (Gibco) at room temperature for 10 minutes prior to addition to the cell culture. After 3 days of expression, culture supernatants were harvested by 5 minutes of centrifugation at 4000xg, 5 minutes of incubation with high-MW PDAD-MAC solution (Sigma Aldrich) to a final concentration of 0.0375%, and a final 5 minutes of centrifugation at 4000xg. Supernatants were clarified via 0.22$\mu$m vacuum filtration and treated with Tris (pH 8.0) and NaCl to final concentrations of 50mM and 350mM, respectively. Iowa43 HA was then purified by batch-binding the clarified, treated supernatants with an appropriate volume of Ni Sepharose Excel resin (GE Healthcare). After incubating in the supernatant for 30 minutes, the resin bed was washed with 10 column volumes of 20mM Tris (pH 8.0), 300mM NaCl buffer and Iowa43 HA was eluted with 3 column volumes of 20mM Tris (pH 8.0), 300mM NaCl, 300mM imidazole buffer. This batch bind process was then repeated with half the original volume of resin. Iowa43 HA was then concentrated using a 10K MWCO Amicon Ultra centrifugal filter unit (Millipore) and sized on a Superdex 200 Increase 10/300 GL column (GE Healthcare) into a final buffer of 20mM Tris (pH 7.5), 150mM NaCl, 5% glycerol. The size exclusion chromatography eluates were re-concentrated in the same manner to 1mg/ml and evaluated for pre- and post-freeze stability via SDS-PAGE prior to characterization by negative stain electron microscopy.

#### 12.2 nsEM Characterization of Commercial and In-House HA Antigen

Grids for both commercial insect-cell-produced and in-house Iowa43 HA antigens were prepared for negative stain characterization following identical protocols. One aliquot of sample was thawed and diluted to 0.01mg/ml into 25mM TBS pH 7.4, 150mM NaCl buffer. 3$\mu$l of the solution was deposited on the carbon side of an ionized 400 mesh copper grid with a 10nm thick carbon film (EMS: CF400-Cu-TH) and particles were allowed to settle for 30 seconds prior to the remaining fluid being wicked away with blotting paper, and replaced with 3$\mu$l 2% Uranyl Formate. Uranyl Formate solution was wicked away and replaced 2 additional times before ultimately being wicked away to allow the sample to dry.

Grids were inserted into a ThermoFisher Talos L120C 120kV electron microscope equipped with a CETA camera for screening and data collection. Samples were imaged with a nominal magnification of 57,000x, which corresponded with a calculated 2.49$\AA$ pixel size. For the commercial insect-cell-produced antigen, 155 micrographs were recorded with a total dose of 55.1 $e^-/\AA^2$. Micrographs were then processed in CryoSPARC v4.4.1 to yield a final population of 150 2D class averages composed of 283,949 particles, which confirmed the presence of monomeric HA antigen. For the in-house produced Iowa43 HA antigen, 222 micrographs were recorded at 57,000x magnification with a total dose of 55.0 $e^-/\AA^2$. Micrographs were then processed in CryoSPARC v4.4.1 to yield a final population of 69 2D class averages composed of 31,616 particles which confirmed the presence of trimeric HA antigen (Extended Data Fig. 11).

#### 12.3 CryoEM Grid Sample Preparation

Flu VHH was combined with Iowa43 HA at a 3:1 molar excess ratio (VHH:HA monomer) at a concentration of 15$\mu$M and promptly prepared for cryo-EM grid freezing. Inaccurately designed VHH was combined with SARS-CoV-2 COVID at a 3:1 molar excess ratio (VHH:COVID monomer) at a concentration of 9$\mu$M and promptly prepared for cryo-EM grid freezing. TcdB at (4.2 mg/mL, 15.56 $\mu$M) was mixed with 3 fold molar excess nanobody or scFv at concentrations of 1-5 mgs. For freezing, the sample was diluted further in 150 mM NaCl, 40 mM Tris/ HCl pH 7.5 to a final concentration of 0.81 mg/mL TcdB. For VHH_TcdB_H2 against TcdB with 100 mM glycine the sample was diluted with 150 mM NaCl, 40 mM Tris/ HCl pH 7.5. For all samples 2$\mu$Ls of Antigen in complex with nanobody or antibody was applied to glow-discharged C-flat holey carbon grids 2.0/2.0-T C-flat holey carbon grids. Vitrification was performed using a Mark IV Vitrobot at 22°C with 100% humidity for all. Blotting was done using a 6.5-7.5 second blot time, a blot force of 0, and a 7.5 second wait time before being immediately plunge frozen into liquid ethane.

#### 12.4 Collection Strategy Notes for TcdB

For TcdB samples initial experiments with VHH_TcdB_H2 yielded apo structures of TcdB, no VHH_TcdB_H2 was bound. Experiment was done in duplicate as follows. The first TcdB sample with VHH_TcdB_H2 was collected on thick ice as the particles appeared more intact visually on micrographs. However the volume was resolved to 5.13 Å and the map was of poor quality. Of note the volume appeared compressed when compared to published TcdB structures, presumably a result of protein flexibility in solution. (Extended Data Figure 13 C,E,G,I) Henceforth the map is called the apo TcdB compressed map (VHH_TcdB_H2 unbound) or apo TcdB compressed map. A fresh batch of VHH_TcdB_H2 was produced to repeat this experiment and a freshly thawed aliquot of TcdB from the same batch was used. For the second collection new grids were prepared and thinner ice was targeted. A 4.00 Å map was resolved with improved map quality, secondary structure clearly visible. (Extended Data Figure 13 D,F,H,J). A notable caveat of targeting thinner ice was the noticeable degradation of a larger population of TcdB. Because of this a mixture of thin and thick ice was targeted for the next three samples. Henceforth the map is called the apo TcdB extended map (VHH_TcdB_H2 unbound) or apo TcdB extended map. As VHH_TcdB_H2 was mostly targeting TcdB through hydrophobic interactions it was hypothesized that lack of binding was the result of aggregation through nonspecific hydrophobic interactions of the VHH. To counteract this, 100 mM glycine was added to the dilution buffer. Only thick ice is targeted here, a consequence of ice conditions available (Extended Data Figure 14).

OrthoRep Affinity Matured VHH_TcdB_H2 (VHH_TcdB_H2_ortho) was complexed and diluted with 150 mM NaCl, 40 mM Tris/ HCl pH 7.5, no glycine added. A mixture of thin and thick ice was targeted. This collection strategy complicated high resolution structure determination by creating at least four states of the sample. TcdB extended VHH_TcdB_H2_ortho AM bound, TcdB compressed VHH_TcdB_H2_ortho AM bound, apo TcdB extended (VHH unbound), apo TcdB compressed (VHH unbound) (Extended Data Figure 15, Extended Data Figure 16). For TcdB + scFv 5 & 6 samples very thin ice was targeted. This collection strategy simplified the data processing and improved view angle distribution. Overall: TcdB flexibility was minimized, view angle distribution improved, and map quality was improved, one caveat was more data was required as more TcdB particles were degraded (Figure 5, Extended Data Figure 27, Extended Data Figure 28). All TcdB samples were collected at a variety of tilt angles including 0, 5, 10, 15, 20 degrees.

#### 12.5 CryoEM data collection

Data was collected automatically using SerialEM [48]. For apo TcdB compressed (VHH unbound), apo TcdB extended (VHH unbound), VHH (glycine added) against TcdB, the VHH_TcdB_H2_ortho against TcdB, VHH_flu_01 against Flu, Inaccurately designed VHH against SARS-CoV-2: a ThermoFisher Titan Krios 300kV TEM equipped with a K3 Summit direct electron detector [49] and BioQuantum Gif energy filter was used with a pixel size of 0.843. For the scFv 5 and 6 against TcdB structures a ThermoFisher Glasios 200kV TEM equipped with a standalone K3 Summit direct electron detector was used with a pixel size of 0.885. Both microscopes were operated in counting mode. Random defocus ranges spanned between -0.8 and -1.8$\mu$m using image shift: with five-shots per hole and nine holes per stage move for the Krios. And one-shot per hole with nine holes per stage move for the Glacios. Altogether 2335, 3430, 6449, 7546, 16956, 5706, 10897, 8304 movies with a dose of 47, 49, 47, 58, 53, 59, 44, 44 $e^-/\text{Å}^2$ were recorded respectively for apo TcdB compressed (VHH unbound), apo TcdB extended (VHH unbound), VHH_TcdB_H2VHH (glycine added) against TcdB, VHH_TcdB_H2_ortho against TcdB, VHH_flu_01 against Flu, Inaccurately designed VHH against SARS-CoV-2, scFv 6 against TcdB, scFv 5 against TcdB.

#### 12.6 CryoEM Data Processing

All data processing was carried out in CryoSPARC [50] (v4.2.2) and CryoSPARC Live. Alignment of movie frames was performed using Patch Motion with an estimated B-factor of 500$\text{Å}^2$, with a maximum alignment resolution set to 5. Defocus and astigmatism values were estimated using Patch CTF with default parameters. Default parameters are used unless otherwise noted below.

##### 12.6.1 VHH_flu_01 against Flu (Figure 3, Extended Data Figure 12)

Influenza A Hemagglutinin particles either bound or not bound to VHH (design_name: *VHH_flu_01*) were initially picked in a reference-free manner using Blob Picker and extracted with a box size of

340 pixels. This was followed by a round of 2D classification and subsequent template-picking using the best 2D class averages low-pass filtered to 20$\AA$. Particles were next picked with Template Picker and were manually inspected before extracting with a box size of 340 pixels for a total of 5,869,679 particles. A round of reference-free 2D classification was next performed in CryoSPARC with a maximum alignment resolution of 6$\AA$. The best classes which revealed clearly visible secondary-structural elements were used for 3D ab initio determination using the C1 symmetry operator. Subsequently, the most promising classes, displaying clearly discernible secondary-structural elements, were utilized for 3D ab initio determination employing the C1 symmetry operator. A total of 866,449 particles underwent 3D hetero refinement, resulting in their categorization into 6 distinct classes. These particles underwent non-uniform refinement, leading to a map resolution of 3.01$\AA$. After this, the curation of exposures involved the selection of the best classes showcasing bound VHHs, thereby eliminating low-quality movies and particles from the data-set, leaving 570,970 particles for further analysis. A subsequent round of non-uniform refinement improved the global resolution estimate to 2.99$\AA$. The dataset was then partitioned based on exposure groups and underwent global CTF correction. Following this, local motion correction was applied to the particles. A subset of 288,441 particles, identified as the best from an additional round of 3D hetero refinement by splitting into 3 classes, underwent further non-uniform refinement, resulting in enhanced density for at least 2 de novo designed VHHs per trimer and a map with a global resolution estimate of 3.14$\AA$. Reference-based motion correction was then conducted to mitigate beam-induced motion between movie frames for each particle. Subsequently, non-uniform 3D refinement with C1 symmetry was performed on the same subset of particles, yielding a final high-resolution map with an estimated global resolution of 3.02$\AA$ following per-particle defocus refinement showing two VHH bound per HA trimer. Local B-factor sharpening with DeepEMhancer [51], utilizing the highRes deep learning model, was employed for display and model building purposes. Local resolution estimates were determined in CryoSPARC using an FSC threshold of 0.143. Finally, the resulting 3D maps, including half maps, final unsharpened maps, and final sharpened maps, were deposited

in the EMDB under accession number EMD-49405.

##### 12.6.2 apo TcdB compressed map (VHH_TcdB_H2 unbound)(Extended Data Figure 13 C,E,G,I)

Blob picking was used initially to pick 449,781 particles with a minimum particle diameter of 200Å and maximum particle diameter of 400Å, 327,917 particles were extracted with an extraction box size of 460 pixels. On these particles 2D classification was used with: number of 2D classes 150, Number of online-EM iterations 50, Number of final full iterations 5. 6 classes with 15,400 particles were used for particle picking with Template Picker. 1,030,408 particles were extracted with a box size of 460 pixels. On these particles 2D classification was used with: number of 2D classes 150, Number of online-EM iterations 50, Number of final full iterations 5. 7 classes with 61,737 particles were used for another round of 2D classification with: number of 2D classes 24, Number of online-EM iterations 20, Number of final full iterations 5. 11 classes with 38,253 particles were used for Non Uniform Refinement with: Number of extra final passes 5. The final volume resolved to 5.13Å. (Extended Data Figure 13 C,E,G,I). The volume appeared compressed when compared to published TcdB structures. Volumes are displayed using UCSF ChimeraX [52, 53, 54].

##### 12.6.3 apo TcdB Extended Thin Ice (VHH_TcdB_H2 unbound)

(Extended Data Figure 13 D,F,H,J) The VHH_TcdB_H2 experiment was run in duplicate with a fresh batch of VHH_TcdB_H2, thinner ice was targeted this time to try and improve the resolution of TcdB. Using Template picking with 11 classes from the previous TcdB sample, 2,644,854 particles were extracted with a box size of 460, fourier cropped to 230 pixels. On these particles 2D classification was run with: number of 2D classes 300, Number of online-EM iterations 40, Number of final full iterations 10. 4 classes with 36,147 particles were used to generate an Ab Initio with: 3 volumes. The best volume with 20,284 particles was used for Non Uniform Refinement with: number of extra final passes 5. A 4.00 Å was generated. (Extended Data Figure 13 D,F,H,J) The volume appeared extended when compared to the previous volume. Volumes are displayed using UCSF ChimeraX [52].

###### 12.6.4 VHH_TcdB_H2 bound - Glycine Added (Extended Data Figure 14)

Initial blob picking identified 1,515,840 particles within a diameter range of 150–250 Å. These were sorted into 250 classes via 2D classification, and the best six classes were used as templates for a second round of template-based picking on denoised micrographs, yielding 2,178,440 particles. Manually inspected picks were extracted using a 520-pixel box size, and 1,363,334 extracted particles underwent a second round of 2D classification into 250 classes. This classification revealed additional TcdB view angles, prompting selection of the 37 best class averages for another round of template picking, inspection, and extraction (520-pixel box size), resulting in a total of 1,374,628 particles. A final round of 2D classification into 250 classes identified 342,132 well-defined TcdB particles, which were subjected to hetero-refinement into six classes, revealing a mixture of bound and apo TcdB populations.

###### 12.6.5 OrthoRep Affinity Matured VHH_TcdB_H2_ortho against TcdB (Extended Data Figure 15, Extended Data Figure 16)

Blob picking was used initially to pick 2,683,500 particles with a minimum particle diameter of 100Å and maximum particle diameter of 160Å. 2D Classification was used in cryoSPARC live with an extraction box size of 460 pixels, 4 classes were picked with 129,474 particles. Micrograph Denoiser was used and a round of template picking was used on the first 3,227 micrographs. 836,745 particles were extracted with a box size of 460 pixels. On these particles 2D classification was used with: number of 2D classes 100, Batchsize per class 400, Number of online-EM iterations 50. 6 classes with 66,042 particles were used to run an Ab Initio Job with: 3 classes. The best volume was used for a round of Non Uniform Refinement to generate a 5.18Å map with: optimize per particle defocus set to true. The 6 classes with 66,042 particles were also used for an additional round of template picking on all 7,546 denoised micrographs. 735,230 particles were extracted with a box size of 460, fourier cropped to 230 pixels. On these particles 2D classification was used with: number of 2D classes 300, Number of online-EM iterations 75, number of final full iterations 25. 25 classes were selected with 100,145 particles. The 5.18Å map and 100,145 particles were used as an input for a round of heterogeneous refinement with 2 volumes generated. A compressed state of TcdB was observed with density for the VHH with 58,101 particles and an extended state of TcdB was observed with poor density (low occupancy) for the VHH with 42,044 particles.

The compressed state of TcdB VHH with 59,101 particles was used for Non Uniform Refinement. A 4.35Å map was generated but map quality was poor, clear density for the VHH was visible at the correct binding site. Using the 4.35Å map and the previously determined apo TcdB extended 4.00Å map (Extended Figure 13 J) another round of heterogeneous refinement was used. Again a compressed state of TcdB was observed with density for the VHH now with 62,787 particles and an extended state of TcdB was observed with poor density (low occupancy) for the VHH with 37,358 particles.

A Non Uniform refinement was run with both of these volumes/particles. A 4.19Å map was generated with a compressed state of TcdB observed with strong density for the VHH and a 4.67Å map with poor density (low occupancy) for the VHH was generated. These two volumes plus the apo TcdB extended 4.00Å map (Extended Data Figure 13 D,F,H,J) were used for another round of heterogeneous refinement with 3 volumes generated. An apo TcdB extended volume was generated with 14,219 particles, compressed state of TcdB was with strong density for the VHH with 48,356 particles, and a extended state of TcdB with density for the VHH with 37,570 particles. Non Uniform Refinement was run on the two VHH bound states, generating a 4.50Å map of extended state of TcdB with density for the VHH and a 4.42Å compressed state of TcdB with good density for the VHH. Both maps were of medium to poor quality overall due to poor view angle distribution.

To address this poor angle distribution template picking was used, we've observed iterative rounds of template picking help find rare views and improve view angle distribution with TcdB. with the 25 2D classes (mentioned above) on all 7,546 denoised micrographs, 899,219 particles were extracted with a box size of 460 pixels, fourier cropped to 230 pixels. On these particles 2D classification was run with: number of 2D classes 300, Number of online-EM iterations 75, Number of final full iterations 25. 14 classes were selected with 59,727 particles and used for a final round of template picking. 1,313,636 particles were extracted with a box size of 460, fourier cropped to 230 pixels. On these particles 2D classification was run with: number of 2D classes 100, Number of online-EM iterations 75, Number of final full iterations 25, Batchsize per class 200. 15 classes were selected with 209,028 particles.

Using 4 volumes (2 previously generated apo state maps, and two maps with VHH density): the apo TcdB extended 4.00Å map (Extended Data Figure 13 D,F,H,J), apo TcdB compressed 5.13 Å map (Extended Data Figure 13 C,E,G,I), the 4.50Å map of extended state of TcdB with density for the VHH, and a 4.42Å compressed state of TcdB with density for the VHH; a final round of heterogeneous refinement was run. 4 major classes were observed: apo TcdB compressed unbound 43,943 particles, compressed TcdB VHH bound 76,997 particles, apo TcdB extended unbound 16,074 particles, TcdB extended VHH bound 72,014 particles. Homogenous refinement was run on the two volumes with TcdB and VHH resolved, 6.27Å map was resolved for TcdB extended VHH bound and 6.74Å map was resolved compressed TcdB VHH bound. Overall map quality was improved but with lower resolution as measured by the GSFSC cutoff of 0.143. Additionally the maps both resolved the presence of aggregation, likely a result of the His tag on the VHH. (Extended Data Figure 15).

The extended state of TcdB with VHH bound was further refined using local refinement. In UCSF Chimera [55] a local map was generated to remove noise from the His tag aggregation, Volume Tools was used to generate a mask around this local volume with: Dilution radius (pix) 10, and Soft padding width (pix) 10, Lowpass filter (Å) 10. Using this mask a Local Refinement Job was run with: Use pose/shift gaussian prior to alignment set to true, Re-center rotations each iteration? Set to true, Re-center shifts each iteration? Set to true. The final volume resolved to 5.71 Å using an FSC threshold of 0.143 and is reported in (Figure 3 I,J,K ; Extended Data Figure 16). Volumes are displayed using ChimeraX.

##### 12.6.6 Inaccurately designed VHH against SARS-CoV-2 (Extended Data Figure 17)

Blob picking was initially done in cryoSPARC live with: a minimum particle diameter of 150 Å, maximum particle diameter of 230 Å, and an extraction box size of 560 pixels. 2D class averaging was run using the 722,692 particles blob picked and extracted with cryoSPARC live with: Number of 2D classes 150, Number of online-EM iterations 50, Number of final full iterations 15, Batchsize per class 300. Following 2D classification 13 classes with 161,326 particles were used for 3D Ab Initio with 3 classes. 2 volumes were poorly resolved and one volume clearly looked like SARS-CoV-2 trimer. This Ab Initio volume was used for heterogeneous refinement with 3 volumes generated. The best volume showing RBD upwards density for 2 RBD domains and 123,782 particles from 2 volumes (both showing strong RBD upwards density for at least 1 RBD domain) was used for Non Uniform Refinement with: Optimize per particle defocus set to true. A 3.38 Å map was generated in C1. Clear density for one upwards RBD was resolved but no VHH density was observed in the map, likely as a result of overall RBD flexibility. (Extended Data Figure 17 C,D,E). Symmetry expansion of all SARS-CoV-2 trimers using the C3 symmetry operator, followed by C1-symmetry 3D classification focused on the RBD, allowed selection of RBDs in the "up" conformation bound to the designed VHH. A final round of local refinement on the best-resolved classes, where VHH-bound RBDs were clearly observed, further enhanced map quality in this region compared to the global refinement (Extended Data Figure 17 F,G,H).

##### 12.6.7 scFv6 against TcdB (Figure 5; Extended Data Figure 27)

10,897 micrographs were denoised with Micrograph Denoiser using a pretrained model. 30 2D classes from VHH_TcdB_H2_ortho TcdB sample were used as templates for Template Picker.

3,357,543 particles were extracted with a box size of 460 pixels and divided into 5 groups (computational reasons) for 2D class averaging with: Number of 2D classes 200, Number of online-EM iterations 75, Number of final full iterations 25. Following 2D classification 8 classes with 56,221 particles were selected. 3D Ab Initio was run and used as an input for Non Uniform Refinement with: Number of extra final passes 5. A 3.86 Å volume was generated with great map quality, TcdB was observed in the extended state as expected due to the collection strategy of targeting only thin ice. Using this map and the previously determined apo TcdB extended map (Extended Data Figure 13 D,F,H,J) a round of heterogeneous refinement was run with the 2 volumes. Two volumes were generated with 14,384 particles in the apo TcdB extended state and 41,837 particles in the TcdB extended scFv bound state. Non Uniform Refinement was run on the scFv bound state volume and corresponding particles with: Number of extra final passes 5. A 3.88 Å volume was generated and used as an input for heterogeneous refinement where 10 volumes were generated. 3,368 particles from the worst volume were removed, the remaining 35,048 particles were used for another round of Non Uniform Refinement with: Number of extra final passes 5. The resulting 3.38 Å map was used as an input for a Reference Motion Correction job. After 34,555 particles were used for another round of Non Uniform Refinement with: Number of extra final passes 5. The resolution improved to 3.57 Å.

Globally the map was well resolved but side chain density was very poor for the epitope and scFv. To overcome this a round of 3D classification was used. It was hypothesized that there was still apo TcdB not removed through heterogeneous refinement. A local map was generated in Chimera [55] using segger including the scFv and a portion for the DRBD domain on TcdB and imported into cryoSPARC, Volume Tools was used to generate a mask around this local volume with: Dilution radius (pix) 5, and Soft padding width (pix) 3. The mask was used as a focus mask and the 3.57 Å map with scFv bound and the apo TcdB extended 4.00 Å map (Extended Data Figure 13 D,F,H,J) were used as initial volumes. For 3D classification: Number of classes 2, Filter Resolution (Å) 6, Force hard classification set to true. Two classes were generated with 19,969 particles and 14,586 particles. Non Uniform refinement was run on both classes, A 4.00 Å map was generated from the volume with 14,586 particles and a 3.60 Å map was generated from the volume with 19,696. The 3.60 Å map is the final map reported (Figure 5). Surprisingly both maps had strong density for the scFv, however the 3.60 Å map had strong density for TcdB residues 1034-1058 forming two alpha helices opposite the epitope on TcdB and the 4.00 Å map had no density for this region, likely the presence of these helices stabilized the epitope. This region is implicated in pore formation in endosomes and delivery of TcdB domains (GTD and CPD) into the cytosol [56]. Additionally, Local Resolution Estimation and Orientation Diagnostics Jobs were run on the final map (Extended Data Figure 27) Volumes are displayed using ChimeraX. Finally, the resulting 3D maps, including half maps, final unsharpened maps, and final sharpened maps, were deposited in the EMDB under accession number EMD-49373.

##### 12.6.8 scFv5 against TcdB (Figure 5; Extended Data Figure 28)

8304 Micrographs were denoised with Micrograph Denoiser using a pretrained model. 30 2D classes from VHH_TcdB_H2_ortho TcdB sample were used as templates for Template Picker. 2,736,960 particles were extracted with a box size of 460 pixels and divided into 4 groups (computational reasons) for 2D class averaging with: Number of 2D classes 200, Number of online-EM iterations 75, Number of final full iterations 25. Following 2D classification 21 classes with 88,602 particles were further 2D classified with: Number of 2D classes 50, Number of online-EM iterations 75, Number of final full iterations 25. Following 2D classification 19 classes with 28,413 particles were examined with curate exposure, 1,177 exposures were removed leaving 7,123 good micrographs and 27,102 particles. Again 2D classification was run with: Number of 2D classes 50, Number of online-EM iterations 75, Number of final full iterations 25. Following 2D classification 16 classes with 16,207 particles were used for 3D Ab Initio with 2 volumes generated. The best volume and all the particles were used for Non Uniform Refinement with: Number of extra final passes 5. A 6.17 Å volume was generated with good map quality for the resolution, TcdB was observed in the extended state as expected due to the collection strategy of targeting only thin ice. Using this map and the apo TcdB extended map (Extended Data Figure 13 D,F,H,J) a round of heterogeneous refinement was run with the 2 volumes. Two volumes were generated with 3,667 particles in the apo TcdB extended state and 12,540 particles in the TcdB extended scFv bound state. Non Uniform Refinement was run on the scFv bound state volume and corresponding particles with: Number of extra final passes 5. The final map had a resolution 6.11 Å, the design was docked into the density. (Figure 5 H,I,J). Local Resolution and Orientation Diagnostics jobs were run on this final volume. (Extended Data Figure 28) Volumes are displayed using ChimeraX.

#### 12.7 CryoEM Model Building

##### 12.7.1 VHH_flu_01 in complex with Iowa43 Influenza H1 HA

The Iowa43 pandemic influenza virus H1 glycoprotein bound to an RFdiffusion-designed HA 20 minibinder (PDB ID: 8SK7) was used as an initial reference for building the cryoEM structure of the A/USA:Iowa/1943 H1N1 bound to the RFdiffusion-designed VHH reported here. The model was first manually edited and trimmed using Coot [57]. The de novo predicted design model for the VHH was used as an initial reference for building into the corresponding density. We further refined each structure in Rosetta using density-guided protocols [58]. EM density-guided molecular dynamics simulations were next performed using ISOLDE, with manual local inspection and guided correction of rotamers and clashes throughout simulated iterations. ISOLDE runs were performed at a simulated 25 Kelvin, with a round of Rosetta density-guided relaxation performed afterward. This process was repeated iteratively until convergence and high agreement with the map was achieved. Multiple rounds of relaxation and minimization were performed on the structure, followed by human inspection for errors after each step. Phenix [59] real-space refinement was subsequently performed as a final step before the final model quality was analyzed using Molprobity [60]. Figures were generated using either UCSF Chimera [55] or UCSF ChimeraX [52]. The structure is deposited in the Protein Data Bank PDB under accession number 9NH7.

##### 12.7.2 scFv6 in complex with TcdB

TcdB was built using the published 3.87 Å crystal structure as a starting model (PDB ID: 6OQ5) [61]. The three bound VHH domains in the crystal structure were removed in PyMOL, and the structure was docked into density using Chimera [55]. The initial model was refined in Coot [57] before alignment with the design model in PyMOL, the scFv docked well in density. The entire model was refined with iterative rounds in Coot [57], Interactive Structure Optimization by Local Direct Exploration (ISOLDE) [62] were performed at a simulated 25 Kelvin, and Phenix [**?** ] real-space refinement. The final model quality was analyzed using Molprobity [60]. The final model and maps are displayed using UCSF ChimeraX [52] (Figure 5). The structure is deposited in the Protein Data Bank PDB under accession number 9NFU.

| | Influenza H1 HA/VHH_flu_01 | TcdB/scFv6 |
|---|---|---|
| **Data Collection** | | |
| Microscope | Titan Krios | Glacios |
| Voltage (kV) | 300 | 200 |
| Detector | Gatan K3 | Gatan K3 |
| Energy Filter | Gatan BioQuantum Gif | None |
| Recording mode | Counting | Counting |
| Magnification | 105,000 X | 45,000 X |
| Frame exposure time (s) | 0.0667 | 0.0505 |
| Movie micrograph exposure time (s) | 5.0 | 5.0 |
| Total dose ($e^{-}/Å^{2}$) | 53 | 44 |
| Under focus range (µm) | 0.8 - 1.8 | 0.8-1.8 |
| Number of movie micrographs | 16,954 | 10,897 |
| **Map Processing** | | |
| Extraction Box Size (pix) | 340 | 460 |
| Initial particle images (no.) | 5,869,679 | 3,357,543 |
| Final particle images (no.) | 288,441 | 14,586 |
| Map resolution (Å) | 3.02 | 3.60 |
| FCS threshold | 0.143 | 0.143 |
| Map resolution range (Å) | 2.3 - 3.7 | 2.5 - 6.0 |
| **Refinement** | | |
| Initial model used | Design Model, 8SK7 | Design Model, 6OQ5 |
| Map resolution (Å) | 3.02 | 3.60 |
| FCS threshold | 0.143 | 0.143 |
| Model resolution range (Å) | 2.3 - 3.7 | 2.5 - 6.0 |
| Map sharpening B factor | 109 | 75.2 |
| Model composition | | |
| Non-hydrogen atoms | 12,976 | 13,490 |
| Protein Residues | 1656 | 1969 |
| Ligands | BMA, NAG | Na |
| *B* factors (Å) | | |
| Protein | 42.92 | 127.88 |
| Ligands | 70.14 | Na |
| R.M.S. deviations | | |
| Bond lengths (Å) | 0.004 | 0.003 |
| Bond angles (°) | 0.591 | 0.597 |
| Validation | | |
| MolProbity score | 1.27 | 1.28 |
| Clashscore | 3.32 | 2.86 |
| Rotamer Outliers (%) | 0.16 | 0.67 |
| Ramachandran plot | | |
| Favored (%) | 97.17 | 96.73 |
| Allowed (%) | 2.83 | 3.27 |
| Disallowed (%) | 0 | 0 |

Supplementary Methods Table 9: CryoEM Data Collection, Processing, and Modeling Statistics

[4] Mikhail Shugay, Dmitriy V. Bagaev, Ivan V. Zvyagin, Renske M. Vroomans, Jeremy Chase Crawford, Garry Dolton, Ekaterina A. Komech, Anastasiya L. Sycheva, Anna E. Koneva, Evgeniy S. Egorov, Alexey V. Eliseev, Ewald Van Dyk, Pradyot Dash, Meriem Attaf, Cristina Rius, Kristin Ladell, James E. McLaren, Katherine K. Matthews, E. Bridie Clemens, Daniel C. Douek, Fabio Luciani, Debbie van Baarle, Katherine Kedzierska, Can Kesmir, Paul G. Thomas, David A. Price, Andrew K. Sewell, and Dmitriy M. Chudakov. VDJdb: a curated database of T-cell receptor sequences with known antigen specificity. *Nucleic Acids Research*, 46(D1):D419–D427, January 2018. ISSN 1362-4962. doi: 10.1093/nar/gkx760.

[5] Philip Bradley. Structure-based prediction of T cell receptor:peptide-MHC interactions. *eLife*, 12:e82813, January 2023. ISSN 2050-084X. doi: 10.7554/eLife.82813.

[6] H. M. Berman, J. Westbrook, Z. Feng, G. Gilliland, T. N. Bhat, H. Weissig, I. N. Shindyalov, and P. E. Bourne. The Protein Data Bank. *Nucleic Acids Research*, 28(1):235–242, January 2000. ISSN 0305-1048. doi: 10.1093/nar/28.1.235.

[7] Sidhartha Chaudhury, Sergey Lyskov, and Jeffrey J. Gray. PyRosetta: a script-based interface for implementing molecular modeling algorithms using Rosetta. *Bioinformatics (Oxford, England)*, 26(5):689–691, March 2010. ISSN 1367-4811. doi: 10.1093/bioinformatics/btq007.

[8] Minkyung Baek, Ivan Anishchenko, Ian R. Humphreys, Qian Cong, David Baker, and Frank DiMaio. Efficient and accurate prediction of protein structure using RoseTTAFold2, May 2023. URL `https://www.biorxiv.org/content/10.1101/2023.05.24.542179v1`. Pages: 2023.05.24.542179 Section: New Results.

[9] Joseph L. Watson, David Juergens, Nathaniel R. Bennett, Brian L. Trippe, Jason Yim, Helen E. Eisenach, Woody Ahern, Andrew J. Borst, Robert J. Ragotte, Lukas F. Milles, Basile I. M. Wicky, Nikita Hanikel, Samuel J. Pellock, Alexis Courbet, William Sheffler, Jue Wang, Preetham Venkatesh, Isaac Sappington, Susana Vázquez Torres, Anna Lauko, Valentin De Bortoli, Emile Mathieu, Sergey Ovchinnikov, Regina Barzilay, Tommi S. Jaakkola, Frank DiMaio, Minkyung Baek, and David Baker. De novo design of protein structure and function with RFdiffusion. *Nature*, 620(7976):1089–1100, August 2023. ISSN 1476-4687. doi: 10.1038/s41586-023-06415-8. URL `https://www.nature.com/articles/s41586-023-06415-8`. Number: 7976 Publisher: Nature Publishing Group.

[10] Jascha Sohl-Dickstein, Eric A. Weiss, Niru Maheswaranathan, and Surya Ganguli. Deep Unsupervised Learning using Nonequilibrium Thermodynamics. In *International Conference on Machine Learning*, volume 37 of *Proceedings of Machine Learning Research*, pages 2256–2265. PMLR, March 2015. URL `https://proceedings.mlr.press/v37/sohl-dickstein15.html`. arXiv:1503.03585 [cond-mat, q-bio, stat].

[11] Jonathan Ho, Ajay Jain, and Pieter Abbeel. Denoising Diffusion Probabilistic Models. In *Advances in Neural Information Processing Systems*, volume 33, pages 6840–6851, 2020. URL https://proceedings.neurips.cc/paper_files/paper/2020/file/4c5bcfec8584af0d967f1ab10179ca4b-Paper.pdf.

[12] John Jumper, Richard Evans, Alexander Pritzel, Tim Green, Michael Figurnov, Olaf Ronneberger, Kathryn Tunyasuvunakool, Russ Bates, Augustin Žídek, Anna Potapenko, Alex Bridgland, Clemens Meyer, Simon A. A. Kohl, Andrew J. Ballard, Andrew Cowie, Bernardino Romera-Paredes, Stanislav Nikolov, Rishub Jain, Jonas Adler, Trevor Back, Stig Petersen, David Reiman, Ellen Clancy, Michal Zielinski, Martin Steinegger, Michalina Pacholska, Tamas Berghammer, Sebastian Bodenstein, David Silver, Oriol Vinyals, Andrew W. Senior, Koray Kavukcuoglu, Pushmeet Kohli, and Demis Hassabis. Highly accurate protein structure prediction with AlphaFold. *Nature*, 596(7873):583–589, August 2021. ISSN 1476-4687 0028-0836. doi: 10.1038/s41586-021-03819-2.

[13] Ting Chen, Ruixiang Zhang, and Geoffrey Hinton. Analog Bits: Generating Discrete Data using Diffusion Models with Self-Conditioning. In *International Conference on Learning Representations*, 2023.

[14] Jason Yim, Andrew Campbell, Andrew Y. K. Foong, Michael Gastegger, José Jiménez-Luna, Sarah Lewis, Victor Garcia Satorras, Bastiaan S. Veeling, Regina Barzilay, Tommi Jaakkola, and Frank Noé. Fast protein backbone generation with SE(3) flow matching, October 2023. URL http://arxiv.org/abs/2310.05297. arXiv:2310.05297 [q-bio].

[15] Jason Yim, Brian L. Trippe, Valentin De Bortoli, Emile Mathieu, Arnaud Doucet, Regina Barzilay, and Tommi Jaakkola. SE(3) diffusion model with application to protein backbone generation, February 2023. URL https://arxiv.org/abs/2302.02277v2.

[16] Chentong Wang, Yannan Qu, Zhangzhi Peng, Yukai Wang, Hongli Zhu, Dachuan Chen, and

Longxing Cao. Proteus: pioneering protein structure generation for enhanced designability and efficiency, February 2024. URL https://www.biorxiv.org/content/10.1101/2024.02.10.579791v1. Pages: 2024.02.10.579791 Section: New Results.

[17] Pablo Gainza, Sarah Wehrle, Alexandra Van Hall-Beauvais, Anthony Marchand, Andreas Scheck, Zander Harteveld, Stephen Buckley, Dongchun Ni, Shuguang Tan, Freyr Sverrisson, Casper Goverde, Priscilla Turelli, Charlène Raclot, Alexandra Teslenko, Martin Pacesa, Stéphane Rosset, Sandrine Georgeon, Jane Marsden, Aaron Petruzzella, Kefang Liu, Zepeng Xu, Yan Chai, Pu Han, George F. Gao, Elisa Oricchio, Beat Fierz, Didier Trono, Henning Stahlberg, Michael Bronstein, and Bruno E. Correia. De novo design of protein interactions with learned surface fingerprints. *Nature*, 617(7959):176–184, May 2023. ISSN 1476-4687. doi: 10.1038/s41586-023-05993-x.

[18] Longxing Cao, Brian Coventry, Inna Goreshnik, Buwei Huang, William Sheffler, Joon Sung Park, Kevin M. Jude, Iva Marković, Rameshwar U. Kadam, Koen H. G. Verschueren, Kenneth Verstraete, Scott Thomas Russell Walsh, Nathaniel Bennett, Ashish Phal, Aerin Yang, Lisa Kozodoy, Michelle DeWitt, Lora Picton, Lauren Miller, Eva-Maria Strauch, Nicholas D. DeBouver, Allison Pires, Asim K. Bera, Samer Halabiya, Bradley Hammerson, Wei Yang, Steffen Bernard, Lance Stewart, Ian A. Wilson, Hannele Ruohola-Baker, Joseph Schlessinger, Sangwon Lee, Savvas N. Savvides, K. Christopher Garcia, and David Baker. Design of protein-binding proteins from the target structure alone. *Nature*, 605(7910): 551–560, May 2022. ISSN 0028-0836, 1476-4687. doi: 10.1038/s41586-022-04654-9. URL https://www.nature.com/articles/s41586-022-04654-9.

[19] Baris E. Suzek, Yuqi Wang, Hongzhan Huang, Peter B. McGarvey, and Cathy H. Wu. Uniref clusters: a comprehensive and scalable alternative for improving sequence similarity searches. *Bioinformatics*, 31(6):926–932, November 2014. ISSN 1367-4803. doi: 10.1093/bioinformatics/btu739. URL http://dx.doi.org/10.1093/bioinformatics/btu739.

[20] Chloe Hsu, Robert Verkuil, Jason Liu, Zeming Lin, Brian Hie, Tom Sercu, Adam Lerer, and Alexander Rives. Learning inverse folding from millions of predicted structures. 162:8946–8970, 17–23 Jul 2022. URL https://proceedings.mlr.press/v162/hsu22a.html.

[21] Valerio Mariani, Marco Biasini, Alessandro Barbato, and Torsten Schwede. lddt: a local superposition-free score for comparing protein structures and models using distance difference tests. *Bioinformatics*, 29(21):2722–2728, August 2013. ISSN 1367-4811. doi: 10.1093/bioinformatics/btt473. URL http://dx.doi.org/10.1093/bioinformatics/btt473.

[22] Minkyung Baek, Frank DiMaio, Ivan Anishchenko, Justas Dauparas, Sergey Ovchinnikov, Gyu Rie Lee, Jue Wang, Qian Cong, Lisa N. Kinch, R. Dustin Schaeffer, Claudia Millán, Hahnbeom Park, Carson Adams, Caleb R. Glassman, Andy DeGiovanni, Jose H. Pereira, Andria V. Rodrigues, Alberdina A. van Dijk, Ana C. Ebrecht, Diederik J. Opperman, Theo Sagmeister, Christoph Buhlheller, Tea Pavkov-Keller, Manoj K. Rathinaswamy, Udit Dalwadi, Calvin K. Yip, John E. Burke, K. Christopher Garcia, Nick V. Grishin, Paul D. Adams, Randy J. Read, and David Baker. Accurate prediction of protein structures and interactions using a three-track neural network. *Science (New York, N.Y.)*, 373(6557):871–876, August 2021. ISSN 1095-9203 0036-8075. doi: 10.1126/science.abj8754.

[23] John L Xu and Mark M Davis. Diversity in the cdr3 region of vh is sufficient for most antibody specificities. *Immunity*, 13(1):37–45, July 2000. ISSN 1074-7613. doi: 10.1016/s1074-7613(00)00006-6. URL http://dx.doi.org/10.1016/S1074-7613(00)00006-6.

[24] Susana Vázquez Torres, Philip J. Y. Leung, Preetham Venkatesh, Isaac D. Lutz, Fabian Hink, Huu-Hien Huynh, Jessica Becker, Andy Hsien-Wei Yeh, David Juergens, Nathaniel R. Bennett, Andrew N. Hoofnagle, Eric Huang, Michael J. MacCoss, Marc Expòsit, Gyu Rie Lee, Asim K. Bera, Alex Kang, Joshmyn De La Cruz, Paul M. Levine, Xinting Li, Mila Lamb, Stacey R. Gerben, Analisa Murray, Piper Heine, Elif Nihal Korkmaz, Jeff Nivala, Lance

Stewart, Joseph L. Watson, Joseph M. Rogers, and David Baker. De novo design of high-affinity binders of bioactive helical peptides. *Nature*, 626(7998):435–442, December 2023. ISSN 1476-4687. doi: 10.1038/s41586-023-06953-1. URL http://dx.doi.org/10.1038/s41586-023-06953-1.

[25] Jeffrey A. Ruffolo, Lee-Shin Chu, Sai Pooja Mahajan, and Jeffrey J. Gray. Fast, accurate antibody structure prediction from deep learning on massive set of natural antibodies. *Nature Communications*, 14(1), April 2023. ISSN 2041-1723. doi: 10.1038/s41467-023-38063-x. URL http://dx.doi.org/10.1038/s41467-023-38063-x.

[26] Andrew Leaver-Fay, Michael Tyka, Steven M. Lewis, Oliver F. Lange, James Thompson, Ron Jacak, Kristian Kaufman, P. Douglas Renfrew, Colin A. Smith, Will Sheffler, Ian W. Davis, Seth Cooper, Adrien Treuille, Daniel J. Mandell, Florian Richter, Yih-En Andrew Ban, Sarel J. Fleishman, Jacob E. Corn, David E. Kim, Sergey Lyskov, Monica Berrondo, Stuart Mentzer, Zoran Popović, James J. Havranek, John Karanicolas, Rhiju Das, Jens Meiler, Tanja Kortemme, Jeffrey J. Gray, Brian Kuhlman, David Baker, and Philip Bradley. ROSETTA3: an object-oriented software suite for the simulation and design of macromolecules. *Methods in Enzymology*, 487:545–574, 2011. ISSN 1557-7988. doi: 10.1016/B978-0-12-381270-4.00019-6.

[27] Naresh Chennamsetty, Vladimir Voynov, Veysel Kayser, Bernhard Helk, and Bernhardt L. Trout. Design of therapeutic proteins with enhanced stability. *Proceedings of the National Academy of Sciences*, 106(29):11937–11942, July 2009. ISSN 1091-6490. doi: 10.1073/pnas.0904191106. URL http://dx.doi.org/10.1073/pnas.0904191106.

[28] Timothy M. Lauer, Neeraj J. Agrawal, Naresh Chennamsetty, Kamal Egodage, Bernhard Helk, and Bernhardt L. Trout. Developability index: A rapid in silico tool for the screening of antibody aggregation propensity. *Journal of Pharmaceutical Sciences*, 101(1):102–115,

January 2012. ISSN 0022-3549. doi: 10.1002/jps.22758. URL http://dx.doi.org/10.1002/jps.22758.

763-1 978-1-62703-764-8. doi: 10.1007/978-1-62703-764-8_5. URL https://link.springer.com/10.1007/978-1-62703-764-8_5. Series Title: Methods in Molecular Biology.

[40] Lorenzo Benatuil, Jennifer M. Perez, Jonathan Belk, and Chung-Ming Hsieh. An improved yeast transformation method for the generation of very large human antibody libraries. *Protein Engineering, Design and Selection*, 23(4):155–159, April 2010. ISSN 1741-0134, 1741-0126. doi: 10.1093/protein/gzq002. URL https://academic.oup.com/peds/article-lookup/doi/10.1093/protein/gzq002.

[41] Mira Loock, Luiza Berenguer Antunes, Rhiannon T Heslop, Antonio Alfonso De Lauri, Andressa Brito Lira, and Igor Cestari. High-Efficiency Transformation and Expression of Genomic Libraries in Yeast. *Methods and Protocols*, 6(5):89, September 2023. ISSN 2409-9279. doi: 10.3390/mps6050089. URL https://www.mdpi.com/2409-9279/6/5/89.

[42] B. I. M. Wicky, L. F. Milles, A. Courbet, R. J. Ragotte, J. Dauparas, E. Kinfu, S. Tipps, R. D. Kibler, M. Baek, F. DiMaio, X. Li, L. Carter, A. Kang, H. Nguyen, A. K. Bera, and D. Baker. Hallucinating symmetric protein assemblies. *Science*, 378(6615):56–61, October 2022. ISSN 1095-9203. doi: 10.1126/science.add1964. URL http://dx.doi.org/10.1126/science.add1964.

[43] Alexandra M. Paulk, Rory L. Williams, and Chang C. Liu. Rapidly Inducible Yeast Surface Display for Antibody Evolution with OrthoRep. *ACS Synthetic Biology*, 13(8):2629–2634, August 2024. ISSN 2161-5063, 2161-5063. doi: 10.1021/acssynbio.4c00370. URL https://pubs.acs.org/doi/10.1021/acssynbio.4c00370.

[44] Alon Wellner, Conor McMahon, Morgan S. A. Gilman, Jonathan R. Clements, Sarah Clark, Kianna M. Nguyen, Ming H. Ho, Vincent J. Hu, Jung-Eun Shin, Jared Feldman, Blake M. Hauser, Timothy M. Caradonna, Laura M. Wingler, Aaron G. Schmidt, Debora S. Marks, Jonathan Abraham, Andrew C. Kruse, and Chang C. Liu. Rapid generation of potent

antibodies by autonomous hypermutation in yeast. *Nature Chemical Biology*, 17(10):1057–1064, October 2021. ISSN 1552-4450, 1552-4469. doi: 10.1038/s41589-021-00832-4. URL https://www.nature.com/articles/s41589-021-00832-4.

[45] Gordon Rix, Rory L. Williams, Vincent J. Hu, Aviv Spinner, Alexander (Olek) Pisera, Debora S. Marks, and Chang C. Liu. Continuous evolution of user-defined genes at 1 million times the genomic mutation rate. *Science*, 386(6722):eadm9073, November 2024. ISSN 0036-8075, 1095-9203. doi: 10.1126/science.adm9073. URL https://www.science.org/doi/10.1126/science.adm9073.

[46] Arjun Ravikumar, Garri A. Arzumanyan, Muaeen K.A. Obadi, Alex A. Javanpour, and Chang C. Liu. Scalable, Continuous Evolution of Genes at Mutation Rates above Genomic Error Thresholds. *Cell*, 175(7):1946–1957.e13, December 2018. ISSN 00928674. doi: 10.1016/j.cell.2018.10.021. URL https://linkinghub.elsevier.com/retrieve/pii/S0092867418313308.

[47] Pulkit Gupta, Zhifen Zhang, Seiji N. Sugiman-Marangos, John Tam, Swetha Raman, Jean-Phillipe Julien, Heather K. Kroh, D. Borden Lacy, Nicholas Murgolo, Kavitha Bekkari, Alex G. Therien, Lorraine D. Hernandez, and Roman A. Melnyk. Functional defects in Clostridium difficile TcdB toxin uptake identify CSPG4 receptor-binding determinants. *Journal of Biological Chemistry*, 292(42):17290–17301, October 2017. ISSN 00219258. doi: 10.1074/jbc.M117.806687. URL https://linkinghub.elsevier.com/retrieve/pii/S0021925820338886.

[48] David N. Mastronarde. SerialEM: A Program for Automated Tilt Series Acquisition on Tecnai Microscopes Using Prediction of Specimen Position. *Microscopy and Microanalysis*, 9(S02):1182–1183, August 2003. ISSN 1431-9276, 1435-8115. doi: 10.1017/S1431927603445911. URL https://academic.oup.com/mam/article/9/S02/1182/6907237.

[49] Ming Sun, Caleigh M. Azumaya, Eric Tse, David P. Bulkley, Matthew B. Harrington,

Glenn Gilbert, Adam Frost, Daniel Southworth, Kliment A. Verba, Yifan Cheng, and David A. Agard. Practical considerations for using K3 cameras in CDS mode for high-resolution and high-throughput single particle cryo-EM. *Journal of Structural Biology*, 213(3):107745, September 2021. ISSN 10478477. doi: 10.1016/j.jsb.2021.107745. URL `https://linkinghub.elsevier.com/retrieve/pii/S1047847721000502`.

[50] Ali Punjani, John L Rubinstein, David J Fleet, and Marcus A Brubaker. cryosparc: algorithms for rapid unsupervised cryo-em structure determination. *Nature Methods*, 14(3): 290–296, February 2017. ISSN 1548-7105. doi: 10.1038/nmeth.4169. URL `http://dx.doi.org/10.1038/nmeth.4169`.

[51] Ruben Sanchez-Garcia, Josue Gomez-Blanco, Ana Cuervo, Jose Maria Carazo, Carlos Oscar S. Sorzano, and Javier Vargas. Deepemhancer: a deep learning solution for cryo-em volume post-processing. *Communications Biology*, 4(1), July 2021. ISSN 2399-3642. doi: 10.1038/s42003-021-02399-1. URL `http://dx.doi.org/10.1038/s42003-021-02399-1`.

[52] Eric F. Pettersen, Thomas D. Goddard, Conrad C. Huang, Elaine C. Meng, Gregory S. Couch, Tristan I. Croll, John H. Morris, and Thomas E. Ferrin. Ucsf chimerax: Structure visualization for researchers, educators, and developers. *Protein Science*, 30(1):70–82, October 2020. ISSN 1469-896X. doi: 10.1002/pro.3943. URL `http://dx.doi.org/10.1002/pro.3943`.

[53] Thomas D. Goddard, Conrad C. Huang, Elaine C. Meng, Eric F. Pettersen, Gregory S. Couch, John H. Morris, and Thomas E. Ferrin. UCSF ChimeraX: Meeting modern challenges in visualization and analysis. *Protein Science*, 27(1):14–25, January 2018. ISSN 0961-8368, 1469-896X. doi: 10.1002/pro.3235. URL `https://onlinelibrary.wiley.com/doi/10.1002/pro.3235`.

[54] Elaine C. Meng, Thomas D. Goddard, Eric F. Pettersen, Greg S. Couch, Zach J. Pearson, John H. Morris, and Thomas E. Ferrin. <span style="font-variant:small-caps;">UCSF

ChimeraX</span> : Tools for structure building and analysis. *Protein Science*, 32(11): e4792, November 2023. ISSN 0961-8368, 1469-896X. doi: 10.1002/pro.4792. URL `https://onlinelibrary.wiley.com/doi/10.1002/pro.4792`.

[55] Eric F. Pettersen, Thomas D. Goddard, Conrad C. Huang, Gregory S. Couch, Daniel M. Greenblatt, Elaine C. Meng, and Thomas E. Ferrin. Ucsf chimera—a visualization system for exploratory research and analysis. *Journal of Computational Chemistry*, 25(13):1605–1612, July 2004. ISSN 1096-987X. doi: 10.1002/jcc.20084. URL `http://dx.doi.org/10.1002/jcc.20084`.

[56] Mengqiu Jiang, Joonyoung Shin, Rudo Simeon, Jeng-Yih Chang, Ran Meng, Yuhang Wang, Omkar Shinde, Pingwei Li, Zhilei Chen, and Junjie Zhang. Structural dynamics of receptor recognition and pH-induced dissociation of full-length Clostridioides difficile Toxin B. *PLOS Biology*, 20(3):e3001589, March 2022. ISSN 1545-7885. doi: 10.1371/journal.pbio.3001589. URL `https://dx.plos.org/10.1371/journal.pbio.3001589`.

[57] Paul Emsley and Kevin Cowtan. *Coot* : model-building tools for molecular graphics. *Acta Crystallographica Section D Biological Crystallography*, 60(12):2126–2132, December 2004. ISSN 0907-4449. doi: 10.1107/S0907444904019158. URL `https://journals.iucr.org/paper?S0907444904019158`.

[58] Julia Koehler Leman, Brian D. Weitzner, Steven M. Lewis, Jared Adolf-Bryfogle, Nawsad Alam, Rebecca F. Alford, Melanie Aprahamian, David Baker, Kyle A. Barlow, Patrick Barth, Benjamin Basanta, Brian J. Bender, Kristin Blacklock, Jaume Bonet, Scott E. Boyken, Phil Bradley, Chris Bystroff, Patrick Conway, Seth Cooper, Bruno E. Correia, Brian Coventry, Rhiju Das, René M. De Jong, Frank DiMaio, Lorna Dsilva, Roland Dunbrack, Alexander S. Ford, Brandon Frenz, Darwin Y. Fu, Caleb Geniesse, Lukasz Goldschmidt, Ragul Gowthaman, Jeffrey J. Gray, Dominik Gront, Sharon Guffy, Scott Horowitz, Po-Ssu Huang, Thomas Hu-

ber, Tim M. Jacobs, Jeliazko R. Jeliazkov, David K. Johnson, Kalli Kappel, John Karanicolas, Hamed Khakzad, Karen R. Khar, Sagar D. Khare, Firas Khatib, Alisa Khramushin, Indigo C. King, Robert Kleffner, Brian Koepnick, Tanja Kortemme, Georg Kuenze, Brian Kuhlman, Daisuke Kuroda, Jason W. Labonte, Jason K. Lai, Gideon Lapidoth, Andrew Leaver-Fay, Steffen Lindert, Thomas Linsky, Nir London, Joseph H. Lubin, Sergey Lyskov, Jack Maguire, Lars Malmström, Enrique Marcos, Orly Marcu, Nicholas A. Marze, Jens Meiler, Rocco Moretti, Vikram Khipple Mulligan, Santrupti Nerli, Christoffer Norn, Shane Ó'Conchúir, Noah Ollikainen, Sergey Ovchinnikov, Michael S. Pacella, Xingjie Pan, Hahnbeom Park, Ryan E. Pavlovicz, Manasi Pethe, Brian G. Pierce, Kala Bharath Pilla, Barak Raveh, P. Douglas Renfrew, Shourya S. Roy Burman, Aliza Rubenstein, Marion F. Sauer, Andreas Scheck, William Schief, Ora Schueler-Furman, Yuval Sedan, Alexander M. Sevy, Nikolaos G. Sgourakis, Lei Shi, Justin B. Siegel, Daniel-Adriano Silva, Shannon Smith, Yifan Song, Amelie Stein, Maria Szegedy, Frank D. Teets, Summer B. Thyme, Ray Yu-Ruei Wang, Andrew Watkins, Lior Zimmerman, and Richard Bonneau. Macromolecular modeling and design in Rosetta: recent methods and frameworks. *Nature Methods*, 17(7):665–680, July 2020. ISSN 1548-7105. doi: 10.1038/s41592-020-0848-2.

[59] Paul D. Adams, Pavel V. Afonine, Gábor Bunkóczi, Vincent B. Chen, Ian W. Davis, Nathaniel Echols, Jeffrey J. Headd, Li-Wei Hung, Gary J. Kapral, Ralf W. Grosse-Kunstleve, Airlie J. McCoy, Nigel W. Moriarty, Robert Oeffner, Randy J. Read, David C. Richardson, Jane S. Richardson, Thomas C. Terwilliger, and Peter H. Zwart. *PHENIX* : a comprehensive Python-based system for macromolecular structure solution. *Acta Crystallographica Section D Biological Crystallography*, 66(2):213–221, February 2010. ISSN 0907-4449. doi: 10.1107/S0907444909052925. URL https://journals.iucr.org/paper?S0907444909052925.

[60] Vincent B. Chen, W. Bryan Arendall, Jeffrey J. Headd, Daniel A. Keedy, Robert M. Immormino, Gary J. Kapral, Laura W. Murray, Jane S. Richardson, and David C. Richard-

son. *MolProbity* : all-atom structure validation for macromolecular crystallography. *Acta Crystallographica Section D Biological Crystallography*, 66(1):12–21, January 2010. ISSN 0907-4449. doi: 10.1107/S0907444909042073. URL https://journals.iucr.org/paper?S0907444909042073.

[61] Peng Chen, Kwok-ho Lam, Zheng Liu, Frank A. Mindlin, Baohua Chen, Craig B. Gutierrez, Lan Huang, Yongrong Zhang, Therwa Hamza, Hanping Feng, Tsutomu Matsui, Mark E. Bowen, Kay Perry, and Rongsheng Jin. Structure of the full-length Clostridium difficile toxin B. *Nature Structural & Molecular Biology*, 26(8):712–719, August 2019. ISSN 1545-9993, 1545-9985. doi: 10.1038/s41594-019-0268-0. URL https://www.nature.com/articles/s41594-019-0268-0.

[62] Tristan Ian Croll. *ISOLDE* : a physically realistic environment for model building into low-resolution electron-density maps. *Acta Crystallographica Section D Structural Biology*, 74(6):519–530, June 2018. ISSN 2059-7983. doi: 10.1107/S2059798318002425. URL https://journals.iucr.org/paper?S2059798318002425.
